# The origin of asexual brine shrimps

**DOI:** 10.1101/2021.06.11.448048

**Authors:** Nicolas Olivier Rode, Roula Jabbour-Zahab, Loreleï Boyer, Élodie Flaven, Francisco Hontoria, Gilbert Van Stappen, France Dufresne, Christoph Haag, Thomas Lenormand

## Abstract

Determining how and how often asexual lineages emerge within sexual species is central to our understanding of sex-asex transitions and the long-term maintenance of sex. Asexuality can arise “by transmission” from an existing asexual lineage to a new one, through different types of crosses. The occurrence of these crosses, cryptic sex, variation in ploidy and recombination within asexuals greatly complicates the study of sex-asex transitions, as they preclude the use of standard phylogenetic methods and genetic distance metrics. In this study we show how to overcome these challenges by developing new approaches to investigate the origin of the various asexual lineages of the brine shrimp *Artemia parthenogenetica*. We use a large sample of asexuals, including all known polyploids, and their sexual relatives. We combine flow cytometry with mitochondrial and nuclear DNA data. We develop new genetic distance measures and methods to compare various scenarios describing the origin of the different lineages. We find that all diploid and polyploid *A. parthenogenetica* likely arose within the last 80,000 years through successive and nested hybridization events that involved backcrosses with different sexual species. All *A. parthenogenetica* have the same common ancestor and therefore likely carry the same asexuality gene(s) and reproduce by automixis. These findings radically change our view of sex-asex transitions in this group, and show the importance of considering asexuality “by transmission” scenarios. The methods developed are applicable to many other asexual taxa.

## Introduction

Understanding why sexual reproduction is so widespread among eukaryotes, despite the well-known costs of sex (Maynard Smith 1978; Otto and Lenormand 2002; Meirmans et al. 2012), requires an understanding of how and how often sex-asex transitions can occur (Delmotte et al. 2001; Simon et al. 2003; Archetti 2004; Lenormand et al. 2016; Engelstädter 2017; Haag et al. 2017; Boyer et al. 2021). Here we refer to asexual reproduction as reproduction without “syngamy” (i.e. without the fusion of male and female gametes, but with the possibility of recombination). Most theoretical models on the origin and frequency of sex-asex transitions adopt a simplistic view and assume that an asexual clone can emerge immediately from a sexual ancestor. Yet, although it is true that extant parthenogenetic species are derived from sexual ancestors, most of these transitions likely occured in several steps and include non-clonal modes of asexual reproduction, which can impact the fitness of asexual lineages at both short and long evolutionary timescales (Asher 1970; Suomalainen et al. 1987; Archetti 2004, 2010; Engelstädter 2017). Historically, cytologists distinguished automictic and apomictic asexuals based on the presence or absence of “mixis” (i.e. fusion of meiotic products). From a genetic standpoint, apomixis refers to clonal reproduction, which is functionally equivalent to mitosis. Cytologic and genetic definitions are contradictory because mixis is not always required for non-clonal reproduction, e.g. when either meiosis I or II is aborted (Asher 1970). Here, we use the genetic definition, and refer to automixis as any modification of meiosis that leads to non-clonal inheritance (i.e., maintenance of non-zero meiotic recombination).

In this study, we focus on sex-asex transitions in animals, aiming at a better understanding of both the origin of parthenogenetic lineages and their genomic evolution. We distinguish among four different types of origin of asexuality. Asexuality can arise either spontaneously (e.g., by mutation, “spontaneous origin”, Simon et al. 2003), through the presence of endosymbionts (“symbiotic origin”; Simon et al 2003), through hybridization between two different sexual species (“hybrid origin”, Cuellar 1987; Moritz et al. 1989; Simon et al. 2003; Kearney 2005), or through transmission from an existing asexual lineage to a new one (hereafter “origin by transmission”). Such transmission events may occur through rare males produced by asexual lineages that may transmit asexuality genes by mating with related sexual females (“contagious asexuality”, Hebert and Crease 1983; Simon et al. 2003; Paland et al. 2005; Jaquiéry et al. 2014). Yet, transmission through crosses between asexual females and sexual males are also possible: either asexual females may rarely produce reduced eggs by meiosis and undergo rare sex, or their unreduced eggs may be fertilized, leading to an elevated ploidy level in the new lineage (e.g., production of a triploid lineage through the fertilization, by a haploid sperm, of an unreduced diploid egg produced by an asexual female).

Because of this diversity of possible scenarios, many simple methods fail to provide a robust approach to elucidate how different asexual lineages emerge and how they relate to each other. We can identify five major hurdles that need to be addressed for a comprehensive understanding of sex-asex transitions in animals. We emphasize that most of these hurdles also undermine our understanding of sex-asex transitions in other eukaryotes, although they may involve other specific issues (e.g. van Dijk 2009; Lee et al. 2010).

First, although traditional phylogenetic methods can be used to study the maternal origin of asexual lineages using mitochondrial markers, they can be misleading in many cases. Technically, the presence of nuclear mitochondrial pseudogenes (“numts”, Lopez et al. 1994) can result in incorrect inferences regarding the age of asexual ineages (Bi and Bogart 2010). More fundamentally, classical phylogenetic methods based on nuclear markers might work when asexuality arises spontaneously and when recombination is absent, but they can be very misleading otherwise. Phylogenetic trees cannot depict the potentially reticulated history of asexual lineages, when the origin of asexual lineages involve crosses and/or when recombination is present. A better approach consists in using the discordances between mitochondrial and nuclear markers to reveal the history of hybrid crosses (Schurko et al. 2009). Finally, with asexuality “by transmission”, the age of asexual lineages becomes an ambiguous concept. Indeed, different parts of the genome of these asexuals may have experienced asexuality for very different periods of time (Tucker et al. 2013).

The second hurdle to understanding sex-asex transitions is that recombination may persist in asexual lineages. While the absence of recombination in apomicts maintains heterozygosity levels across generations (except for mutation, gene conversion, and mitotic recombination events), the presence of recombination under automixis can result in a loss of heterozygosity (LOH) between generations. When the rate of LOH is heterogeneous across genomic regions along the chromosome (ranging from 0% to 100%, depending on the distance of the region from the centromere (Nougué et al. 2015*b*; Svendsen et al. 2015; Boyer et al. 2021)), different genomic regions will coalesce at different points in time, providing different phylogenetic signals. Hence, there are considerable uncertainties in the patterns of molecular variation to be expected within and among asexual genomes, as classic genetic distance metrics do not account for heterogeneity in LOH among markers. In addition, LOH could have different fitness consequences, depending on the mode of asexuality (Archetti 2004, 2010; Engelstädter 2017) or on the sex-determination system of the ancestral sexual species (Engelstädter 2008). For instance, LOH in ZW asexual females could produce low fitness ZZ and WW offspring, reducing the rate of transition to asexuality in ZW sexual species.

The third hurdle to understanding sex-asex transitions is that asexuality is often associated with polyploidy, at least in animals (Moritz et al. 1989; Dufresne and Hebert 1994; Otto and Whitton 2000), so that studying the origin of asexuality (e.g. allo- or autopolyploid origin) requires the use of specific genetic distance metrics that are defined across different ploidy levels (Clark & Jasienuk 2011). In addition, both the lack of dosage information (i.e. exact number of each allele) or the variation in nuclear DNA content among individuals from the same ploidy level can make the discrimination among ploidy levels difficult (e.g. Neiman et al. 2011). Finally, elucidating the role of polyploidy in sex-asex transitions is difficult. Most studies fail to reveal whether polyploidy is a cause, a consequence, or even just a correlate of asexuality in animals (Neiman et al. 2014).

The fourth hurdle to understanding sex-asex transitions is the potential occurrence of rare sex events in asexual taxa, in other circumstances than those of contagious asexuality, mentioned above. Meiosis might sometimes occur normally in asexual females (e.g. De Meester et al. 2004; Rey et al. 2013; Boyer et al. 2021). A haploid egg produced through regular meioses can be fertilized by a haploid sperm from a rare male of the same or a different asexual lineage or from a male from related sexual species. These rare events of cryptic sex may be difficult to detect in the field, especially if divergence between parents is low or if sampling is incomplete.

A fifth hurdle to understanding sex-asex transitions is the technical and methodological difficulty of identifying and exhaustively sampling the closest extant sexual species of asexual lineages, as they often have very different geographical distributions (Kearney 2005). In addition, the closest sexual populations might be extinct or might themselves result from the hybridization between divergent sexual populations, so that many different sex-asex transition scenarios need to be considered. Overall, except in species where asexuality is directly caused by endosymbionts, ruling out a potential hybrid origin of asexuality and demonstrating that asexuality arose spontaneously remains very difficult, due to this sampling challenge.

These biological and methodological challenges might exist for virtually all asexual taxa, and failing to address them consistenly and jointly might provide an incomplete picture of the origin and evolution of asexual lineages. Some of these issues have recently been addressed in several systems; for instance the description of automixis in Daphnia (Svendsen et al. 2015), the identification asexuality genes in various asexual arthropods (Sandrock and Vorburger 2011; Tucker et al. 2013; Yagound et al. 2020), or the occurrence of recombination in ancient asexual such as bdelloid rotifers (Simion et al. 2020).

In this study, we address all the hurdles that compromise our understanding sex-asex transitions in *Artemia parthenogenetica*, a group of asexual crustaceans that has been studied for decades and is emblematic of the multiple challenges encountered when studying the origin asexual species. This group includes diploid and polyploid asexual lineages that are found worldwide except on the American continent (Bowen et al. 1978; Browne 1992). The distribution of asexual lineages within the genus is exceptionally asymmetric, as they are all more closely related to Old-world sexual species: *A. sinica*, *A. tibetiana*, *A. urmiana* and an uncharacterized species from Kazakhstan, hereafter named *A.* sp kaz (Asem et al. 2016; see Table 1 for the nomenclature and abbreviations of the different taxa). The geographical distribution of both diploid and polyploid asexuals is much larger than that of these sexual species. *Artemia* resting stages can be dispersed by waterbirds (Sánchez et al. 2012) or humans (Rode et al. 2013) over large geographical distances.

**Table 1.** List of species names and abbreviations

| *Species name* | *abbreviation* | *mt abbreviation* |
| --- | --- | --- |
| *A. sinica* | *Asin* | *mt-sin* |
| *A. urmiana* | *Aurm* | *mt-urm* |
| *A. tibetiana* | *Atib* | *mt-tib* |
| *A. sp. Kazakhstan* | *Akaz* | *mt-kaz* |
| Diploid *A. parthenogenetica* (2n) with *Aurm*-type mitochondria | *Ap2n-urm* | *mt-2nu* |
| Diploid *A. parthenogenetica* (2n) with *Akaz*-type mitochondria | *Ap2n-kaz* | *mt-2nk* |
| Triploid *A. parthenogenetica* (3n) | *Ap3n* | *mt-3n* |
| Tetraploid *A. parthenogenetica* (4n) | *Ap4n* | *mt-4n* |
| Pentaploid *A. parthenogenetica* (5n) | *Ap5n* | *mt-5n* |

Although the origin of asexual *Artemia* lineages has been extensively studied since Bowen *et al*. (1978), previous studies have failed to jointly address the five hurdles to understanding sex-asex transitions mentioned above. In particular the paternal origins of asexual lineages have never been investigated so that neither a potential hybrid nor a contagious origin of asexuality have been tested. Fig. 1 presents a synthesis of the major conclusions of landmark papers. Briefly, asexuality in *Artemia* was first thought to have a single origin (Bowen and Sterling 1978; Abreu-Grobois and Beardmore 1982; Beardmore and Abreu-Grobois 1983; Abreu-Grobois 1987), such that the species was considered as an ‘ancient asexual scandal’ (Judson and Normark 1996). When both nuclear and mitochondrial markers and more sexual species (*Aurm*, *Atib* and *Asin*) were considered, the evidence pointed to multiple and much more recent origins of asexuality (Baxevanis et al. 2006; Muñoz et al. 2010; Maniatsi et al. 2011; Eimanifar et al. 2015; Asem et al. 2016). These phylogenetic investigations reported that *A. parthenogenetica* was likely to be a polyphyletic group (Baxevanis et al. 2006; Maniatsi et al. 2011). Based on mitochondrial data, *Ap2n* falls into two distinct maternal lineages, *Ap2n-kaz* and *Ap2n-urm,* whose mitochondrial haplotypes are closer to those of *Akaz* (mt-2n*k*) and *Aurm* (mt-2n*u*) respectively (Muñoz et al. 2010). A third *Ap2n* mitochondrial lineage has also been recently described (Maccari et al. 2013*a*). Based on mitochondrial and nuclear data, *Ap3n* are thought to maternally derive either from *Aurm* (Maniatsi et al. 2011) or from *Ap2n* (Asem et al. 2016). *Ap4n* and *Ap5n* are thought to have emerged successively: the former from an *Asin* female and the latter from an *Ap4n* female (Maniatsi et al. 2011; Asem et al. 2016).

**Figure 1.**
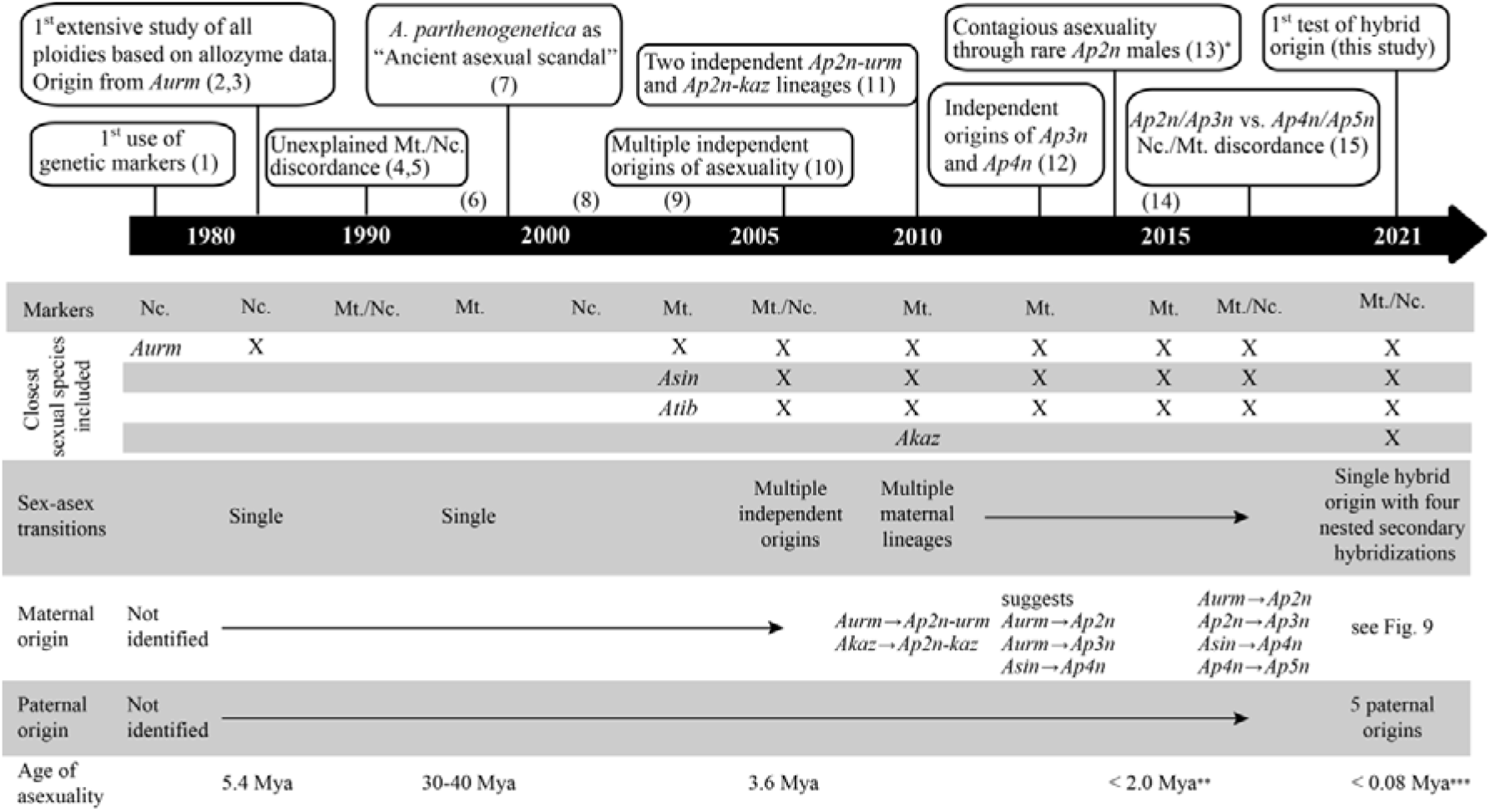
Time line of the major findings regarding the orign of diploid and polyploid asexual Artemia lineages. References : 1 : (Bowen and Sterling 1978); 2 : (Abreu-Grobois and Beardmore 1982) ; 3 : (Beardmore and Abreu-Grobois 1983); 4 : (Browne and Hoopes 1990); 5 : (Browne and Bowen 1991); 6: (Perez et al. 1994) ; 7 : (Judson and Normark 1996); 8 : (Nascetti et al. 2003) ; 9 : (Bossier et al. 2004) ; 10 : (Baxevanis et al. 2006) ; 11 : (Muñoz et al. 2010) ; 12 : (Maniatsi et al. 2011) ; 13 : (Maccari et al. 2013a) ; 14 : (Eimanifar et al. 2015); 15 : (Asem et al. 2016). Mya: millions year ago. *: F1 hybrids between Ap2n and Aurm observed by (1) but no contagion observed before (13). **: Based on Aurm/Ap2n-kaz divergence. ***: Age of the most recent common ancestor of all extent asexual lineages (asexuality gene(s) may be older).

The reproductive mode of *A. parthenogenetica* has also been intensely investigated for over a century. Cytogenetic studies in diploids have yielded contradictory results by reporting almost all known forms of automixis (Narbel-Hofstetter 1964; Nougué et al. 2015*b*), while most cytogenetic studies in polyploids have reported that they reproduce through apomixis (Barigozzi 1974). Recent genetic studies showed that *Ap2n* reproduces through a mechanism genetically equivalent to central fusion automixis (Nougué et al. 2015*b*; Boyer et al. 2021), which leads to loss of heterozygosity in centromere-distal chromosomal regions due to recombination (Svendsen et al. 2015). *A. parthenogenetica* has a ZW sex-determination system and produces males at low frequency (Stefani 1964). These rare males are thought to arise through rare recombination events between the Z and W chromosomes in *Ap2n* females, which results in a LOH at the sex locus and the production of ZZ males (Browne and Hoopes 1990; Abreu-Grobois and Beardmore 2001; Boyer et al. 2021). Last, for a long time, rare males have been thought to be useless and irrelevant to *Ap2n* reproduction (MacDonald and Browne 1987; Simon et al. 2003), but experiments showed they can transmit asexuality when crossed with *Aurm* and *Akaz* sexual females (Maccari et al. 2014; Boyer et al. 2021). Moreover, recent experiments have shown that some *Ap2n* females can engage very rarely in sex in the laboratory, with normal meiosis and recombination (Boyer et al. 2021). Hence contagious asexuality and/or sexual reproduction may occur at small rates within *Ap2n* asexuals *in natura*.

In addition, previous studies also suffered from a number of limitations. The first limitation is the lack of reliable nuclear markers available across asexual and sexual taxa. The most informative dataset (23 allozyme markers) dates back to the 80’s, but shows a limited resolution with few alleles per marker (Abreu-Grobois and Beardmore 1982). Similarly, nuclear sequences (ITS1, Na^+^/K^+^ ATPase, etc.) show limited diversity across asexual and sexual taxa (Baxevanis et al. 2006; Asem et al. 2016). Microsatellite markers described in Muñoz *et al.* (2008) present null alleles and fail to amplify in some sexual species (Maccari et al. 2013*b*). Mitochondrial data also present limitations due to the potential co-amplification of nuclear pseudogenes (Wang et al. 2008). Indeed, several studies have reported difficulties in amplifying COI sequences using universal primers or primers designed for the distantly related *A. franscicana* (Wang et al. 2008) and *A. salina* (Asem et al. 2016). Mitochondrial-nuclear discordance represents crucial evidence for hybridization scenarios, yet many studies did not combine nuclear and mitochondrial data (Fig. 1) and relied only on mitochondrial data for taxonomic identification (Sainz-Escudero et al. 2021). Furthermore, in many studies ploidy of all samples is not directly assessed or is based on existing literature regarding populations previously sampled in the same locality (e.g. Baxevanis et al. 2006). However, a same locality may host different (sexual and/or asexual) populations whose occurrence varies spatially or temporally (e.g. Agh et al. 2007). Importantly, ploidy cannot be assessed based on the number of alleles at each locus genetic markers as *Artemia* individuals from each ploidy level often exhibit only two alleles at each locus (Nougué et al. 2015*a*); whether this lack of variation is the result of LOH or of some other mechanisms has never been studied. Finally, no previous study has included all known sexual species and all ploidy levels. A failure to include the most closely related sexual species may result in asexuality appearing more ancient than it actually is (e.g. Perez et al. 1994). Similar biases due to limited sampling are frequent in studies of the age and origin of asexual taxa (Tucker et al. 2013).

In this paper, we investigate potential origins “by transmission” of diploid and polyploid asexual lineages, considering all new experimental information regarding the reproductive mode of *A. parthenogenetica* (Maccari et al. 2014; Nougué et al. 2015*b*; Boyer et al. 2021). We test the hypothesis that diploid and polyploid asexual lineages may represent a mixture of different “hybrids” resulting from several events of contagion and secondary backcrosses with different sexual species. We build a series of tailored population genetic methods to test whether asexual *Artemia* of various ploidies have a hybrid origin and we attempt to identify the corresponding parental species. We also investigate whether secondary crosses or contagion events can be identified. To this end, we conduct the first study that includes an exhaustive sampling of asexual lineages (*Ap2n-kaz*, *Ap2n-urm*, *Ap3n*, *Ap4n*, *Ap5n*) and major sexual relatives (*Aurm, Akaz*, *Atib*, *Asin*) and that combines nuclear and mitochondrial data with ploidy data (based on flow cytometry). Finally, we also test for the presence of cryptic sex in *Ap2n*. In the absence of sexual reproduction, we can indeed expect *Ap2n* individuals with different mitochondrial haplotypes to be characterized by different and specific nuclear backgrounds. In contrast, in the presence of cryptic sex, we expect a discordance between mitochondrial haplotypes and nuclear backgrounds (hereafter “mito-nuclear discordance”). We find that, after a single hybrid origin of one diploid asexual lineage, all other asexuals emerged through a series of four nested hybridization events involving several sexual species. Overall, this new approach changes our view of sex-asex transitions in *Artemia*. It may prove to be a valuable tool to investigate sex-asex transitions in other taxa, especially when classical phylogenetic approaches are not appropriate.

## Methods

### Samples

Based on the existing literature (Abreu-Grobois and Beardmore 1982; Muñoz et al. 2010; Maniatsi et al. 2011; Maccari et al. 2013*a*), we chose 37 populations from Eurasia and Africa, including both asexual strains (described as diploid, triploid, tetraploid and pentaploid) and the four closest sexual species (Fig. 2). We obtained samples from cyst bank collections and wild-collected adults (Fig. 2; Table S4). Cysts were hatched and individuals maintained following protocols described in (Rode et al. 2011). It has recently been shown that at least some *Ap2n* females can reproduce sexually in the laboratory at a rate of ~2‰ in the presence of males (Boyer et al. 2021). However, for simplicity (and because the capacity to undergo rare sexual reproduction *in natura* is unknown), we only categorized individuals are sexuals vs. asexuals and did not consider facultative asexuality. The reproductive mode of each population was verified based on the presence or absence of males among adults. When at least one male was present, we separated asexual from sexual females according to morphological characters (Maccari et al. 2013*b*). All five populations with at least one male consisted of a mixture of asexual females with females from different sexual species (*A. franciscana*: AIM, BOL, SAG, *A. sinica*: DON, or *A. salina*: BDP).

**Figure 2.**
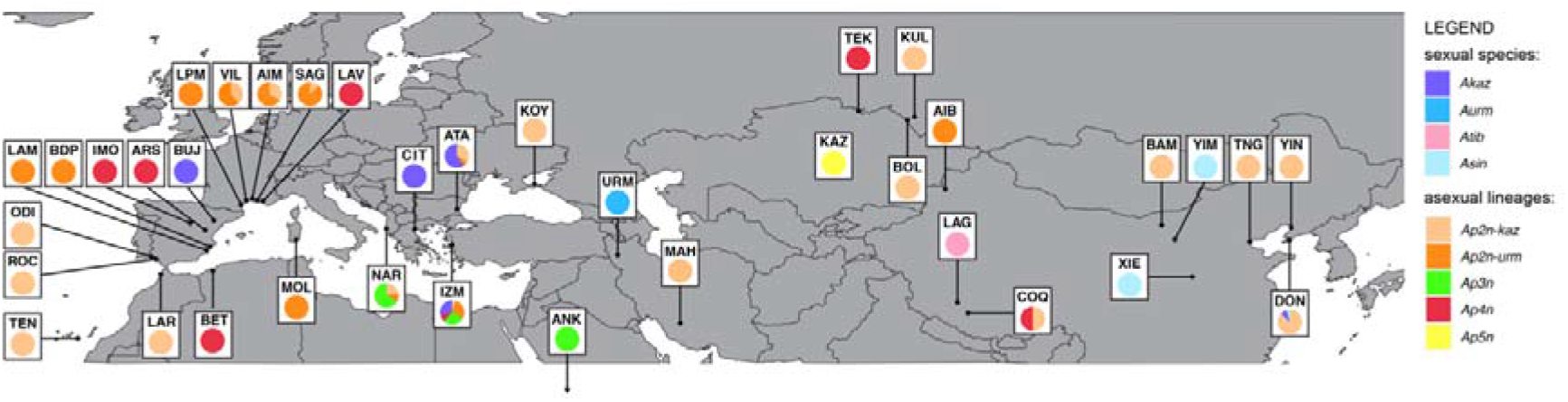
Geographic distribution of the sexual and asexual Artemia samples genotyped in this study. Each color corresponds to a mitogroup. AIB: Aibi Lake (China); AIM: Aigues-Mortes (France); ANK: Ankiembe (Madagascar); ARS: Arcos de la salinas (Spain); ATA1/ ATA2: Atanasovko Lake (Bulgaria); BAM: Unknown, Bameng area (China); BDP: Bras del port (Spain); BET: Bethioua (Algeria); BOL: Bolshoye Yarovoe (Russia); BUJ: Bujaraloz (Spain); CIT: Citros (Greece); COQ: Co Qen (China); DON: Dongjiagou (China); IMO: Imón (Spain); IZM: Izmir (Turkey); KAZ: Unknown location (Kazakhstan); KOY: Koyashskoye (Ukraine); KUL: Kulundinskoye (Russia); LAG: Lagkor Co (China); LAM: La Mata (Spain); LAR: Larache (Morocco); LAV: Lavalduc (France); LPM: La Palmes (France); MAH: Maharlu Lake (Iran); MOL: Molentargius (Italy); NAR: Narte (Albania); ODI: Odiel (Spain); ROC: N. S. Rocío (Spain); SAG: Salin De Giraud (France); TEK: Teke Lake (Kazakhstan); TEN: Tenefé (Spain); TNG: Tanggu (China); URM: Urmia Lake (Iran); VIL: Sète-Villeroy (France); XIE: Xiechi Lake (China); YIM: Unknown location, Yimeng area (China); YIN: Yingkou (China). Due to the scale of the map, the ANK Ap3n population is represented with an arrow pointing towards its location (Ankiembe, Madagascar). Mitochondrial data could not be obtained for KOY (Koyashskoye, Ukraine), which is represented based on the analysis of 17 Ap2n-kaz individuals from Maccari et al (2013a) that used the same cyst sample.

### Ploidy

We characterized the ploidy of putatively diploid and polyploid asexual females, as well as of males and females of each sexual species (147 individuals in total) using flow cytometry as described in (Nougué et al. 2015*b*) and sup. mat. 1. Data from (Nougué et al. 2015*b*) were added to the dataset, resulting in a total sample size of 206 individuals (Table S4). We tested for difference in genome size between asexual and sexual lineages using *t*-tests. All significant *P-values* remained significant after Bonferroni correction, so we present only the uncorrected values for simplicity.

### COI genotyping

A fragment of the mitochondrial Cytochrome c Oxidase Subunit I (COI) gene of 336 individuals (Table S4) was amplified using primers 1/2COI_Fol-F and 1/2COI_Fol-R following the protocol of Muñoz et al. (2010). Because these primers turned out to lack specificity and resulted in the amplification of a large number of numts (see below), we amplified the COI sequence of 23 additional individuals using the more specific primers Co1APAR-F(5’- TTTGGAGCTTGAGCAGGAAT-3’) and Co1APAR-R(5’- TGCGGGATCAAAGAAAGAAG-3’) (see sup. mat. 2). PCR products were purified and directly sequenced (i.e. with no cloning), using an ABI Prism Big Dye Terminator Kit (Applied Biosystems, Warrington, UK) on an ABI PRISM 3130xl sequencer. After removing sequences with indels and sequences from the same individual that were identical, we recovered 359 sequences that were split into two datasets: dataset1, which only included high quality sequences (198 sequences including at least 538 sites in the final alignment and without any ambiguous position), and dataset2, which included all other sequences (161 sequences that were either too short or included one or more ambiguous positions). COI phylogenetic inferences are often biased by the inclusion of numts (Buhay 2009) or chimeric sequences that result from the coamplification by PCR of mitochondrial sequences with contaminating sequences (either numts or mitochondrial sequences from other individuals; e.g. Dubey et al. 2009). We developed a new method to identify potential chimeric sequences based on a dataset composed of known numts sequences (see “chimera detection” section below). To build this dataset, we performed additional cloning and sequencing of PCR fragments amplified either from total DNA or from DNA enriched from mitochondrial DNA (sup. mat. 2, Table S4). Cloning allowed the recovery of 32 numt sequences without indels (which we identified as minority sequences among those obtained from the same individual through cloning, sup. mat. 2) and seven mitochondrial sequences (hereafter dataset3). We built a dataset that combined these 39 sequences (dataset3), our 198 sequences (dataset1) and 748 *Artemia* spp. high quality sequences from GenBank without any indel (sup. mat. 2). All analyses of the final dataset of 985 COI sequences were performed in R 3.6.3 (www.r-project.org).

### Analyses of mitochondrial data (1): pseudogene detection

We aligned sequences using MAFFT v7.427 (Katoh et al. 2002) as implemented within the package ips (v0.0.11; Heibl 2008) with default settings. To detect potential numts without indels, we translated sequences into amino acid sequences using the invertebrate mitochondrial DNA genetic code. To detect potential numts, we tested for changes in the polarity of amino acid residues (Kunz et al. 2019). For each sample, we estimated the absolute difference in polarity (i.e. PP1 in Cruciani et al. 2004) between derived and ancestral amino-acid sequences. Based on the observed distribution of this polarity difference across the 985 protein sequences (Fig. S1, sup. mat. 3), we set 0.1 as the threshold in polarity differences above which sequences were labeled as potential numts. This procedure allowed the reliable detection of 65% of the reference numts obtained by cloning (i.e. 21 of the 32 known numts). We also detected 26 additional sequences (including 25 sequences from GenBank) that were labelled as potential numts for the rest of the analyses.

### Analyses of mitochondrial data (2): haplotype reference set

Poorly aligned positions were removed using gblocks 0.91b as implemented within the package ips (v0.0.11; Heibl 2008) and collapsed into 230 unique haplotypes using the haplotype function of the package haplotypes (v1.1.2; Aktas 2020).

### Analyses of mitochondrial data (3): chimera detection

We designed a quantitative test to detect and exclude potential chimeric sequences. The principle of the method is to determine, for each focal sequence, how many mutations could be “explained” by assuming that this sequence represents a chimera. We compared each haplotype sequence to the remaining 229 sequences in the dataset to identify the most similar one. For each mutation differing between the focal sequence and the most similar one, we provide a score of one if this mutation (i.e. the same SNP) was found in another unrelated sequence of the dataset or zero otherwise. For each of the 230 haplotype sequences, we computed the sum of this score over the different SNPs of each haplotype. We considered that each sequence could have acquired a mutation which happened to be present in an unrelated sequence of the dataset (score =1), but that it could not have acquired two or more of these mutations (score >1). Hence, haplotype sequences with score equal or greater than two were considered as potentially chimeric (see sup. mat. 4 for details). We removed 80 haplotypes (corresponding to 101 samples) that appeared as potential chimeras resulting from the co-amplification of a mitochondrial sequence and either a numt or contamination from other mitochondrial sequences. Overall, we obtained 123 reference non-chimeric haplotypes (corresponding to 884 samples).

### Analyses of mitochondrial data (4): haplotype assignation

To study the phylogenetic relationship among major asexual and sexual taxa (*Ap2n-kaz*, *Ap2n-urm*, *Ap3n*, *Ap4n*, *Ap5n*, *Aurm*, *Akaz*, *Atib*, *Asin*), we built a haplotype network based on the 123 reference haplotypes and using the parsimnet function (95% probability of parsimony, Templeton et al. 1992) of the package haplotypes (v1.1.2; Aktas 2020). We found the same tree topology when building a maximum likelihood phylogeny with the phangorn package (v2.5.5 Schliep 2011).

In addition, we assigned the 61 *Ap2n* sequences in dataset2 to either *Ap2n-kaz*, *Ap2n-urm* or other sequences (i.e. numts or chimeras). To do so, we aligned them with the 123 reference haplotype sequences and with the 107 numt and chimeric sequences and estimated pairwise genetic distances as described above. For each sequence in dataset2, the mitochondrial haplotype was assigned based on the identity of the closest reference haplotype(s). Whenever a numt, a potential numt or a chimera was found among the closest sequences, the *Ap2n* mitochondrial haplotype was set as “unknown”. We could successfully assign to either *Ap2n-kaz* or *Ap2n-urm* 46 of the *Ap2n* sequences.

### Microsatellite genotyping

We genotyped 432 individuals with a panel of 12 microsatellite markers (see Muñoz et al. 2008; Nougué et al. 2015*a* for details regarding markers and amplification protocol). Data from Nougué *et al.* (2015*b*) were added to the dataset, resulting in 489 typed individuals (Table S4, sup. mat. 5). Standardization was achieved by adding DNA from the same individual onto the different plates. Genotype data at three microsatellite markers (Apdq01TAIL, Apdq02TAIL, Apdq03TAIL) were only used when investigating the relationship among *Ap2n*, but excluded when analyzing the full dataset due to the presence of null alleles in polyploids and sexuals (Maniatsi et al. 2011).

### Microsatellite data analyses (1): Lynch distance

The Lynch genetic distance is a genetic distance estimate based on a band sharing index, i.e., one minus the similarity index, equation 1 in Lynch (1990). It is an appropriate metric to broadly compare well-separated sexual and asexual groups with different ploidy levels, such as *Ap2n*, *Ap3n*, *Ap4n, Ap5n,* and the different sexual species (i.e. ignoring the occurrence of LOH in *Ap2n*). We computed the average pairwise nuclear distance between and within lineages using the Lynch genetic distance, as implemented in the *polysat* package (v1.7-4; Clark and Jasieniuk 2011) in R. To visualize distance data, we transformed individual pairwise distances into Principal Coordinate Axes using the ‘cmdscale’ function in the *stats* v3.6.3 package. To investigate the homogeneity of the reference sexual populations, we investigate whether they show a signal of admixture (Estoup et al. 2016). We examined the variation in allele sizes within individuals from each sexual taxon (*Aurm*, *Akaz*, *Atib*, *Asin*; sup. mat. 6).

### Microsatellite data analyses (2): genetic distance for automictic *Ap2n*

Because it does not account for the possibility of LOH, the Lynch genetic distance is of limited use regarding relationships within *Ap2n*. We therefore present a new genetic distance measure for automicts, accounting for the different possible paths between diploid genotypes (sup. mat. 6, Table S1). We assumed that mutations rate is the same across loci, but that LOH varies across loci proportionally to the average inbreeding coefficient (*Fis*), which can be independently estimated. This new measure weights events according to the relative magnitude of LOH and mutation events (with rates *r* and *μ* respectively). This genetic distance is a proxy for the time length of the path between individuals (or averaged across different possible paths, according to their relative probability of occurrence). For instance, with this new distance measure, two individuals with genotype AA and AB are weighed as distant if the locus has a strongly negative *Fis* (low LOH rate), since their difference likely results from a single A to B mutation. However, they are not weighed as distant if the locus has a strongly positive *Fis* (high LOH), as AA can result from an LOH event from an AB parent.

### Monophyly of *Ap2n-kaz* and *Ap2n-urm* clades

In the absence of sexual reproduction, we expect *Ap2n-kaz* and *Ap2n-urm* individuals to be characterized by different and specific nuclear backgrounds. In contrast, in the presence of sexual reproduction between *Ap2n-kaz* and *Ap2n-urm* individuals, we expect a discordance between mitochondrial haplotypes and nuclear backgrounds (hereafter “mito-nuclear discordance”). Using data from the 12 microsatellite makers, we used our new genetic distance metric to compute pairwise genetic distances among 127 *Ap2n* individuals with known mitochondrial haplotypes. Using a randomization test, we first investigated whether the genetic distance between *Ap2n-kaz* and *Ap2n-urm* lineages was significantly larger than that within *Ap2n-kaz* or *Ap2n-urm* lineages. As this test only considers average nuclear distances between and within lineages, the difference might be significant even if *Ap2n-kaz* and *Ap2n-urm* clade are not monophyletic (e.g. due to rare events of sexual reproduction between *Ap2n-kaz* and *Ap2n-urm* individuals). Using the pairwise distance matrix computed above, we first built a neighbor-joining (NJ) tree using the *nj* function from the ape package (v5.4-1; Paradis and Schliep 2019) and estimated branch length using non-negative least squares (nnls.tree function from the phangorn package, v2.5.5 Schliep 2011). We computed the 95% confidence interval of this branch length by resampling microsatellite markers to build 1000 bootstrap replicates. Finally, we built NJ trees separately for *Ap2n-kaz* and *Ap2n-urm* and assembled them into a single tree where each lineage is monophyletic. We again estimated the branch length of this tree using non-negative least square and then tested whether the estimated branch length was outside of the 95% confidence interval computed above.

### Evolutionary origin of *Ap2n-kaz* and *Ap2n-urm*

To investigate the evolutionary relationships between *Ap2n-kaz* and *Ap2n-urm* and the three sexual species (*Aurm*, *Akaz* and *Asin*), we considered different scenarios based on the Lynch genetic distance (Fig. 3). We considered two independent spontaneous origins within *Aurm* and *Akaz* (Fig. 3A), a single spontaneous origin followed by a hybridization event (Fig. 3B), two independent hybridization events (Fig. 3C), and one hybridization and one backross event (Fig. 3D). These four scenarios were considered with or without the presence of an unknown species, denoted *Aunk*, which is assumed to carry mt-*urm*. We did not consider the scenarios where *Aunk* carried mt-*kaz*. Indeed, nuclear data indicated that all *Ap2n* are much closer to *Akaz* than to *Aurm* (see Results), and it is therefore very likely that *Ap2n-kaz* inherited their mitochondria directly from *Akaz*. Because the outgroup *Asin* can have different positions (it can be closest to either *Akaz*, *Aurm* or *Aunk*), each scenario was evaluated assuming all three possible topologies. For each topology, we described each branch length by a parameter. The number of identifiable parameters is given for each scenario in Fig. 3. For scenarios in which *Ap2n* arose spontaneously, a branch length between *Ap2n* and the sexual species was included. When *Ap2n* arose from a hybrid cross, the genetic distance between them and a given sexual species was computed as the averages of the branch lengths between either parent and that sexual species. The model corresponding to each scenario was fitted using least square to the matrix of Lynch genetic distance among *Akaz*, *Aurm*, *Asin*, *Ap2n-kaz* and *Ap2n-urm*. To avoid any confounding effect due to potential cryptic sex between *Ap2n-kaz* and *Ap2n-urm*, we computed this matrix after removing 13 *Ap2n* individuals with discordant mito-nuclear data (see result section). We assumed that genetic distance within each species or within each of two *Ap2n* lineages was negligible compared to the genetic distance among species (i.e. we ignored divergence within each sexual or asexual lineage). Models were compared based on corrected Akaike's information criterion (AICc, Hurvich and Tsai 1989). We computed the difference(ΔAICc) between the AICc of a given model and that of the model with lowest AICc. Models with ΔAICc higher than two were considered as poorly supported (Burnham and Anderson 2002).

**Figure 3.**
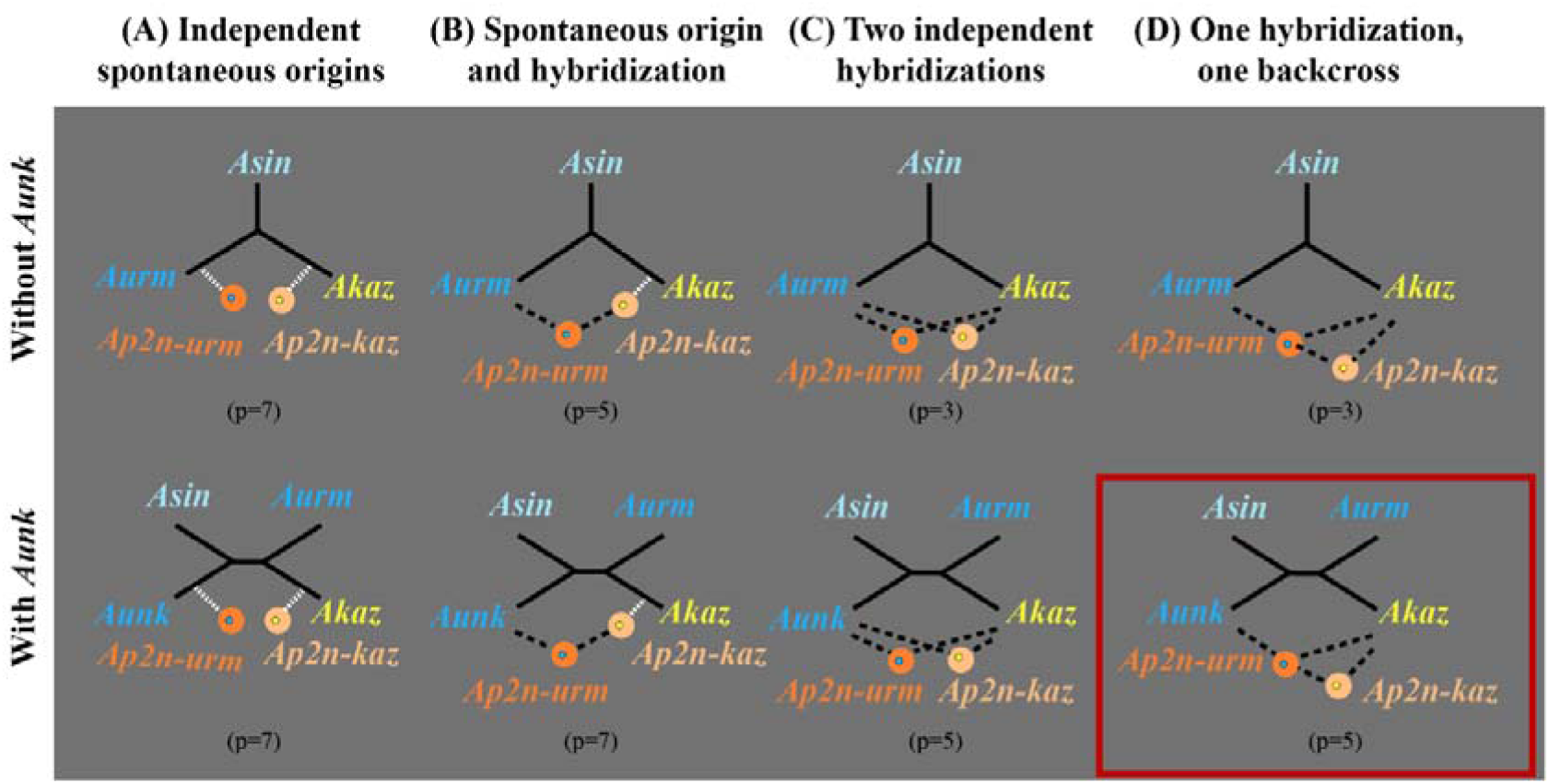
Scenarios for the origin of Ap2n-kaz and Ap2n-urm. The two lineages are respectively depicted by light and dark orange circles. The color of the dot within each circle represents the maternal lineage of the mitochondrion (blue: mt-2nu from an Aurm mother, yellow: mt-2nk from an Akaz mother). White dotted lines and black dashed lines indicate spontaneous and hybrid origins, respectively. In (A) and in the presence of Aunk, the nine branch lengths are not identifiable. The branch length leading to Aunk cannot be fitted and was therefore dropped. In (B), the scenarios illustrated assumes a spontaneous origin of Ap2n-kaz in Akaz followed by a cross with Aurm or Aunk. We also considered the reciprocal scenario (origin in Aurm or in Aunk and cross with Akaz). In (D), the scenarios illustrated shows an origin of Ap2n-urm through hybridization followed by a backcross with Akaz. The reciprocal scenario with a backcross with Aurm or Aunk was also considered. Only one topology (where Asin is closest to Aunk) is represented for all scenarios involving Aunk (second row). The two other topologies were also considered (Asin closest either to Akaz or Aurm, respectively). The best model is indicated by the red rectangle and involved one hybridization with an unknown species and backcross on Akaz (bottom right). The number of parameters fitted for each model (corresponding to the number of identifiable branch lengths) is given by the number between parentheses below each scenario.

### Evolutionary origin of *Ap3n*, *Ap4n* and *Ap5n*

We assumed the maternal origin of the ancestor of *Ap3n*, *Ap4n* and *Ap5n* lineages to be known based on mitochondrial data (see results below). For each ploidy level, we compared different scenarios involving different paternal origins (see sup. mat. 7 for details). For each ploidy level and each scenario, we simulated 10,000 synthetic hybrids using a custom script in R. For each hybrid, we first randomly sampled a mother with the observed mitochondrial haplotype in our dataset (i.e. *Ap2n-kaz* for *Ap3n*, *Asin* for *Ap4n* and *Ap4n* for *Ap5n*). Second, to draw a haploid genotype (representing the sperm genotype), we randomly sampled an individual for each paternal origin (i.e. *Akaz*, *Ap2n-kaz*, *Ap2n-urm*, *Aurm*, *Atib* or *Asin*) and randomly sampled one allele of this individual at each locus. We assumed that the 12 microsatellite loci were unlinked, so that probabilities of sampling alleles were independent across loci. We assumed that alternative scenarios involving fertilization by an unreduced sperm were less likely (e.g. origin of *Ap3n* through the fertilization of a reduced *Ap2n-kaz* egg by an unreduced sperm or origin of *Ap4n* through the fertilization of an unreduced *Asin* egg by an unreduced sperm). We then computed average Lynch genetic distance based on the 100 synthetic hybrids closest to *Ap3n*, *Ap4n* or *Ap5n* individuals in our dataset.

## Results

### Ploidy characterization

The results from the flow cytometry measurements are summarized in Fig. 4. The ploidy levels detected in each population were in good agreement with those found in previous cytological or genetic studies (Abreu-Grobois and Beardmore 1982; Muñoz et al. 2010; Maccari et al. 2013*a*), except for two *Ap5n* populations previously described as *Ap4n* (BUJ and CIT, Abatzopoulos et al. 1986; Amat et al. 1994; Maniatsi et al. 2011). Interestingly, the genome size of *Ap2n-kaz* was not significantly different from that of *Ap2n-urm* (4.74 pg SD=0.17 vs. 4.65 pg SD=0.20 respectively; t=−1.93, df=50.48, *P-value*=0.059). The *Ap2n-kaz* genome size was significantly lower than *Akaz* (4.92 pg SD=0.14; t=−3.21, df=14.74, *P-value*=0.006), and *Ap2n-urm* genome size was significantly higher than *Aurm* (4.22 pg SD=0.20; t=4.46, df=5.67, *P-value*=0.005). The genome size of *Ap3n* (7.13 pg SD=0.43) was consistent with their ploidy level and was not significantly different from 1.5 times that of *Ap2n-kaz* (t=−0.16, df=37.63, *P-value=*0.88). Although *Ap4n* harbor an *Asin* mitochondrion (Asem et al. 2016), their genome size (10.15 pg SD=0.34) was more than twice that of *Asin* (2 × 4.74=9.49 pg SD=0.24; t=4.44, df=6.38, *P-value*=0.004). In contrast, the size of the genome of *Ap5n* (12.22 pg SD=0.45) that also harbor an *Asin* mitochondria was not significantly different from the value 2.5 times that of *Asin* (t=2.15, df=4.57, *P-value*=0.09). These observations seem inconsistent with the scenario of an origin of *Ap4n* through an endoduplication in *Asin*. Interestingly, the genome size of *Atib* was 57%, 34% and 39% larger than that of *Aurm*, *Akaz* and *Asin* respectively, suggesting an increase in genome size in the lineage leading to *Atib*.

**Figure 4.**
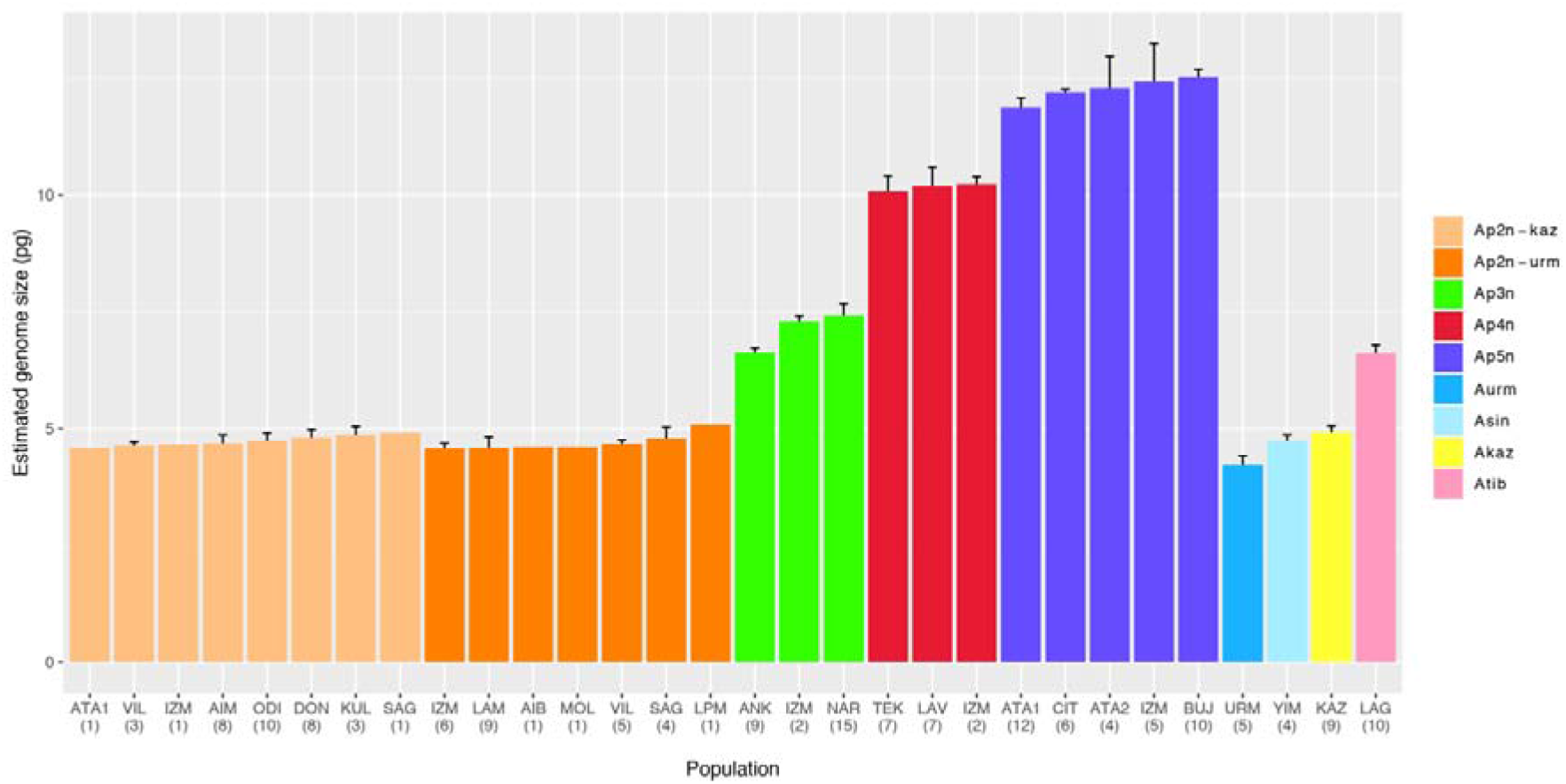
Estimated genome size of Artemia sexual and asexual lineages. Genome size in pg of diploid Ap2n-kaz and Ap2n-urm, triploid, tetraploid and pentaploid asexual lineages and related sexual species (A. urmiana, A. sinica, A. sp Kaz, and A. tibetiana). Mean ± SD C-values are shown. Abbreviations for population labels are provided as in Fig. 2. Number below population labels represent sample sizes. 37 Ap2n individuals had unknown mitochondrial haplotypes and are not represented.

### COI genotyping

*Ap2n*, *Ap3n*, *Ap4n* and *Ap5n* were found in 24, 3, 7 and 5 populations respectively (Table S4, Fig. 2). Among the 37 populations sampled, only five (ATA, IZM, NAR, COQ and DON) were composed of individuals with different ploidies. Similarly, among the 24 *Ap2n* populations, 13 comprised only individuals with *Ap2n-kaz* haplotypes, five comprised only individuals with *Ap2n-urm* haplotypes and six (AIM, ATA, VIL, SAG, IZM, NAR) comprised individuals with both *Ap2n-kaz* and *Ap2n-urm* haplotypes (Fig. 2). Consistent with previous studies, *Ap2n-kaz* is the clade with the largest geographic distribution (Fig. 2). When including all sequences found in GenBank, the distribution of *Ap2n-urm* is restricted to the Mediterranean area (Spain, France, Italy and Turkey), around the Black sea (Atanasovko Lake, Bulgaria; Oybuskoye Lake, Ukraine) and in Western China (Aibi Lake, Lagkor Co). Polyploids also have a large geographic distribution (Fig. 2).

Among the 950 sequences from dataset1 and from GenBank, we found 47 sequences that had a large change in the polarity of the amino-acid sequence and 88 sequences that included two or more mutations found in another unrelated sequence of the dataset. These sequences were likely to be numts and chimeras respectively. The remaining 850 COI sequences were collapsed into 123 unique haplotypes of diploid mt-2n*k*, diploid mt-2n*u*, triploid, tetraploid and pentaploid asexual lineages and related sexual species (Fig. 5). In line with Maccari et al. (2013*a*), we found three networks separated by more than 30 mutation steps. The fourth network described in their study corresponded to pseudogenes (GenBank accession: EF615587-8) that differ from other sequences by several non-synonymous mutations that changed polarity. Each asexual lineage was characterized by a majority haplotype found at high frequency in many populations and recently-derived satellite haplotypes found at a lower frequency in a few populations (Fig. 5), as previously observed in *Ap2n* lineages (Muñoz et al. 2010; Maccari et al. 2013*a*). This observation of low haplotypic diversity with a star-like shape is consistent with a recent range expansion of the different asexual lineages, which now have widespread geographical distributions. No haplotype was shared between asexual lineages and either of the sexual species. The majority haplotype of *Ap2n-kaz* differed by six mutations from the closest *Akaz* haplotype (but one haplotype of *Ap2n-kaz* differed from *Akaz* by just one mutation). The majority haplotype of *Ap2n-urm* differed by one mutation from the closest *Aurm* haplotype. The majority hapotype of *Ap3n* were identical to the closest *Ap2n-kaz* haplotype, and *Ap4n* and *Ap5n* had the same majority happlotype, which differed by a single mutation from the closest *Asin* haplotype (Fig 5). We found a 18% divergence between *Aurm* and *Asin* COI haplotypes from reference mitochondrial genomes used in Sainz-Escudero et al. (2021), which corresponds to a divergence time of 6.6 Mya (5.47-7.40 My) in their study. Assuming that COI substitution rate is constant, one mutation in Fig. 5 corresponds to a divergence of 0.019% and represents an approximate age of 0.068 Myr (0.056-0.076 Myr). Using divergence between *Atib* and *Asin* COI haplotypes and the divergence time estimate from Sainz-Escudero et al. (2021) provides a qualitatively similar age estimate of 0.062 Myr (0.051-0.079 Myr). Although these results rely on strong assumptions (accurate dating of the fossil used for calibration and constant molecular clock), they suggest that all extant asexual lineages potentially emerged more recently than previously thought (i.e. less than 80,000 years ago).

**Figure 5.**
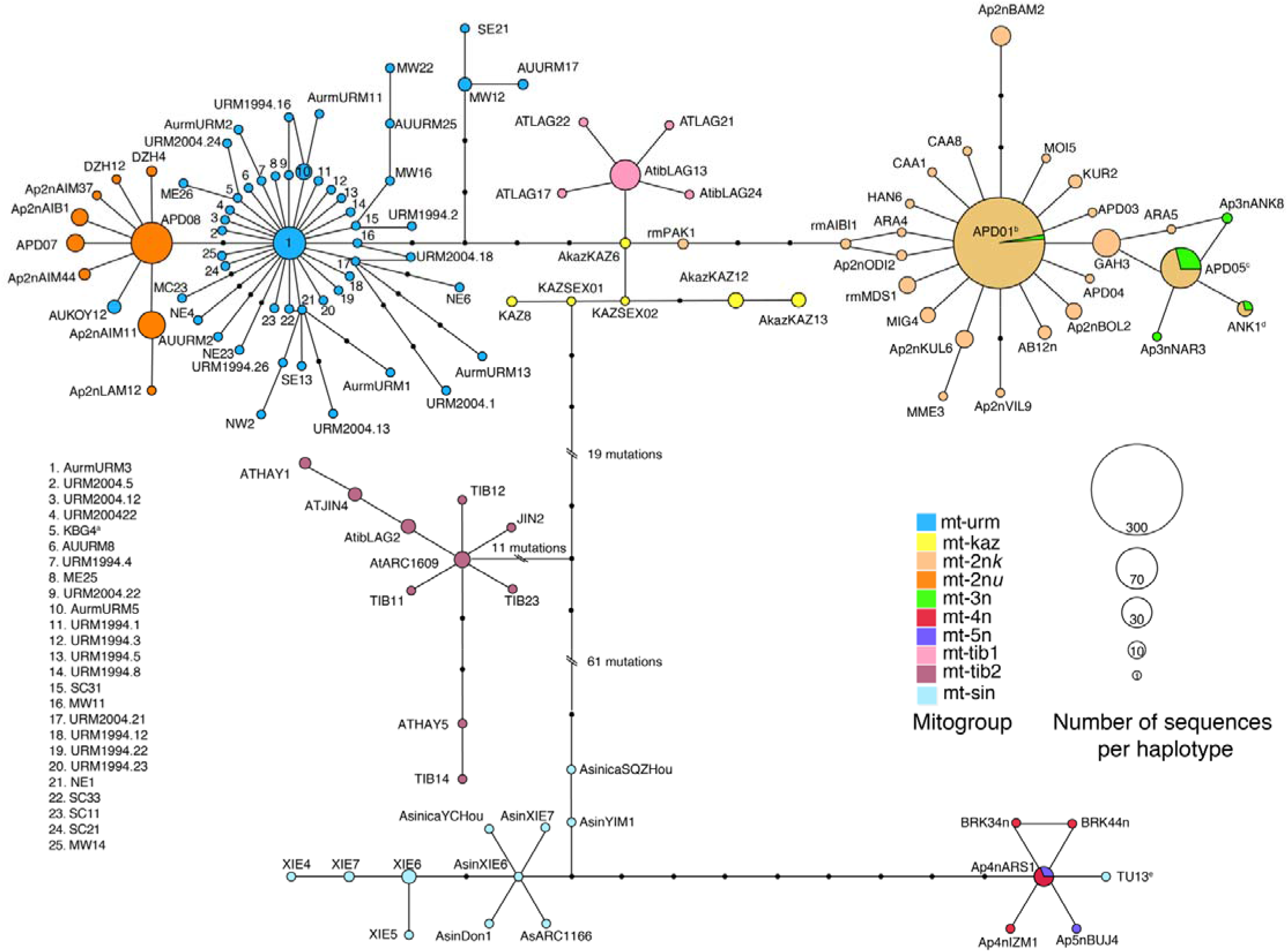
Statistical parsimony network of mitochondrial haplotypes from diploid mt-2nk, diploid mt-2nu, triploid mt-3n, tetraploid mt-4n and pentaploid mt-5n asexual lineages and related sexual species (mt-urm: A. urmiana, mt-sin: A. sinica, mt-kaz: A. sp Kaz, and mt-tib: A. tibetiana). Haplotype including samples with different ploidies are represented with pie charts. Circle diameter is proportional to the relative haplotype frequency among the 850 sequences that were neither numts nor chimeras. Connecting lines indicate single substitutions and small black circles represent putative missing haplotypes. Haplotype codes correspond to those reported in Table S5. ^a^The KBG4 sequence of a cyst from Kara-Bogaz-Gol (Turkmenistan) was molecularly assigned to Aurm by Eimanifar et al. 2014. ^b^Sequences from five Ap3n individuals (based on cytology; Asem et al. 2016) from Aibi Lake (China) had an APD01 Ap2n-kaz haplotype. ^c^Sequences from five Ap2n individuals (based on cytology; Asem et al. 2016) from Akkikol Lake (China) had an ANK1 Ap3n haplotype. ^d^Sequences from 52 individuals (based on morphology; Muñoz et al. 2010, Maccari et al. 2013a, Eimanifar et al. 2014, Eimanifar et al. 2015) had the same APD05 haplotype as Ap3n individuals in our dataset. ^e^The TU13 Asin sequences from Siberia was found to be chimeric between the Ap4nARS1 mitochondrial haplotype and an Ap5n numt, it was included to illustrate that Ap4n and Ap5n likely originated from Siberian Asin.

The first network includes haplogroups corresponding to *Ap2n-kaz*, *Ap2n-urm*, *Ap3n*, *Aurm*, *Akaz*, and *Atib*. This result confirms the existence of the two distinct major *Ap2n* lineages, *Ap2n-kaz* and *Ap2n-urm*. The third *Ap2n* lineage described in (Maccari et al. 2013*a*) possibly represents a chimera between two divergent PCR-amplified sequences in that study (Fig. S2). Consistent with Maccari et al. 2013a, the haplogroup mt-tib1 included most *Atib* sequences from Lagkor Co (also known as Gaize lake, Zheng and Sun 2013).

Triploid samples (*Ap3n*), whose assignation is based on flow cytometry and nuclear genotype, were found to be nested within *Ap2n-kaz* (Fig. 5). This suggests that triploids are maternally derived from this diploid lineage. *Ap3n* were characterized by a very low mitochondrial diversity (Table 2) despite their large geographical distribution (Madagascar, Turkey, Albania). When including sequences from other studies (retrieved from NCBI), we found triploids with sequences identical to those of the closely related *Ap2n-kaz* haplotypes and we observed diploids with sequences identical to those of *Ap3n* haplotypes (Fig. 5).

**Table 2.** Mitochondrial (Kimura-2-parameter on COI sequences) genetic distances between and within mitogroups of sexual species and asexual lineages. See Table S5 for the Accession Numbers of the 123 unique haplotype sequences.

|  | *Aurm* | *Ap2n-urm* | *Ap2n-kaz* | *Ap3n* | *Akaz* | *Atib (LAG)* | *Atib (others)\** | *Asin* | *Ap4n* | *Ap5n* |
| --- | --- | --- | --- | --- | --- | --- | --- | --- | --- | --- |
| *Aurm* | 0.007 | . | . | . | . | . | . | . | . | . |
| *Ap2n-urm* | 0.009 | 0.004 | . | . | . | . | . | . | . | . |
| *Ap2n-kaz* | 0.026 | 0.029 | 0.005 | . | . | . | . | . | . | . |
| *Ap3n* | 0.030 | 0.033 | 0.007 | 0.003 | . | . | . | . | . | . |
| *Akaz* | 0.018 | 0.021 | 0.016 | 0.021 | 0.005 | . | . | . | . | . |
| *Atib (LAG)* | 0.017 | 0.020 | 0.015 | 0.015 | 0.007 | 0.003 | . | . | . | . |
| *Atib (others)\** | 0.075 | 0.080 | 0.081 | 0.083 | 0.073 | 0.074 | 0.006 | . | . | . |
| *Asin* | 0.184 | 0.188 | 0.175 | 0.175 | 0.180 | 0.175 | 0.167 | 0.006 | . | . |
| *Ap4n* | 0.206 | 0.210 | 0.196 | 0.196 | 0.202 | 0.197 | 0.187 | 0.021 | 0.002 | . |
| *Ap5n* | 0.206 | 0.210 | 0.197 | 0.196 | 0.202 | 0.197 | 0.184 | 0.021 | 0.002 | 0.002 |

\* Distances based on *Atib* sequences from other populations than Lagkor Co (LAG) were computed separately.

Among sexuals, *Aurm* had a larger haplotypic diversity than *Akaz* and *Atib* from Lagkor Co, which is likely due to a larger sample size, which increases the likelihood of sampling rare alleles (Fig. 5, Table 2).

A second network consisted of one *Atib* sequence from Lagkor Co (AtibLAG2) and sequences from other sexual populations (Haiyan Lake, Jingyu Lake, Nima, Yangnapengco, Qi Xiang Cuo, etc.) from the Qinghai-Tibet plateau, that we refer to as mt-tib2 (Fig. 5). This haplogroup differed by more than 35 mutation steps from mt-tib1 and segregates at a frequency of ~3% in Lagkor Co (1 over 34 *Atib* sequences). No sequence from the mt-tib1 haplogroup was found in populations from Haiyan and Jingyu Lakes. Only a few sequences per population were available from GenBank, so making general conclusions is not possible.

The third network included sequences from *Ap4n*, *Ap5n* and *Asin*. We found that some *Ap4n* and *Ap5n* samples shared the same mitochondrial haplotype (Fig. 5). We found low haplotype diversity among *Ap4n* and *Ap5n* haplotypes (in contrast to Asem et al. 2016; Table 2). The diversity found in *Ap4n* and *Ap5n* by Asem *et al.* (2016) was due to 12 sequences, which were likely to be numts or chimeras according to our analysis. *Ap4n* and *Ap5n* were closest to a haplotype from an *Asin* sample from Lake Dus-Khol (aka Lake Svatikovo, East Siberia, Russia; Naganawa and Mura 2017), which was found to be chimeric between the *Ap4n* mitochondrial haplotype and an *Ap5n* numt (Table S5). This suggests that *Asin* from Lake Dus-Khol, *Ap4n* and *Ap5n* share the same mitochondrial haplotype and at least one numt. Hence, *Ap4n* might have originated in East Siberia from an *Asin* mother. Mitochondrial diversity levels in the *Asin* population from Xiechi Lake (AsinXIE and XIE haplotypes on Fig. 5) were similar to those observed in other sexual species.

### Microsatellite genotyping

Genetic distances within and among asexual and sexual lineages are represented in Fig. 6 and Table 3. The proportion of total variability explained by the first, second and third axes of the PCoA were to 28.6%, 25.9% and 19.2% respectively (Fig. 6). We observed a larger genetic diversity within both *Ap2n-kaz* and *Ap2n-urm*, than within *Ap3n*, *Ap4n,* or *Ap5n* polyploids. Sharing of multilocus genotypes among populations was rare except for some geographically close populations (*Ap2n*: VIL/AIM, YIN/DON, *Ap5n*: IZM/ATA/CIT). Consistent with mitochondrial data, *Ap2n-kaz* were more closely related with *Akaz* than with *Aurm*. Surprisingly, *Ap2n-urm* were also more closely related to *Akaz* than to *Aurm*. Compared to *Ap2n-kaz* and *Ap2n-urm*, *Ap3n* were more closely related to *Aurm*. Compared to *Ap2n-kaz*, *Ap2n-urm* and *Ap3n*, *Ap4n* and *Ap5n* were more closely related to *Asin*, consistent with mitochondrial data. Variation in microsatellite allele size was smaller in *Akaz* and *Asin* (YIM population) than in *Aurm* and *Atib*, which suggests that the later two populations might be admixed (Fig. S3, sup. mat. 6).

**Figure 6.**
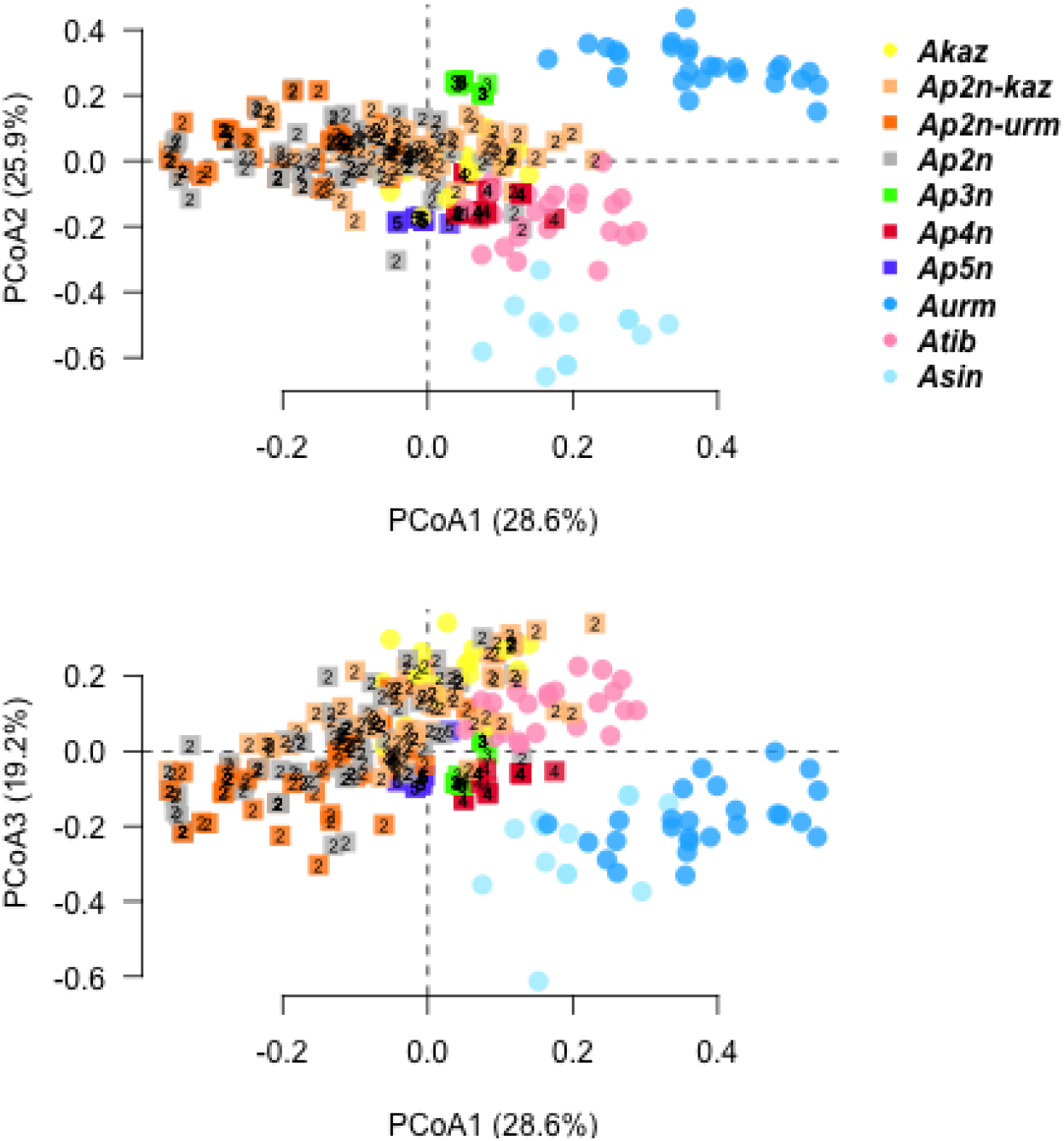
Principal coordinate analyses of asexual lineages and sexual species. The first three principal coordinate axes are shown, with the percentage of variation represented by each axis given between parentheses. Sexual and asexual taxa are represented with filled circles and filled squared, respectively. Numbers within filled squares represent the ploidy level. Ap2n individuals with unknown mitochondrial haplotypes are represented in grey.

**Table 3.** Nuclear genetic distances between and within mitogroups of sexual species and asexual lineages. Lynch distance was computed based on the 9 microsatellite markers from Nougué et al (2015a). The 113 Ap2n individuals with unknown mitochondrial haplotypes were excluded for the computation.

|  | *Aurm* | *Ap2n-urm* | *Ap2n-kaz* | *Ap3n* | *Akaz* | *Atib* | *Asin* | *Ap4n* | *Ap5n* |
| --- | --- | --- | --- | --- | --- | --- | --- | --- | --- |
| *Aurm* | 0.40 | . | . | . | . | . | . | . | . |
| *Ap2n-urm* | 0.71 | 0.30 | . | . | . | . | . | . | . |
| *Ap2n-kaz* | 0.67 | 0.35 | 0.30 | . | . | . | . | . | . |
| *Ap3n* | 0.52 | 0.42 | 0.41 | 0.06 | . | . | . | . | . |
| *Akaz* | 0.69 | 0.43 | 0.34 | 0.47 | 0.26 | . | . | . | . |
| *Atib* | 0.71 | 0.53 | 0.45 | 0.51 | 0.41 | 0.27 | . | . | . |
| *Asin* | 0.90 | 0.75 | 0.75 | 0.83 | 0.76 | 0.58 | 0.18 | . | . |
| *Ap4n* | 0.62 | 0.41 | 0.37 | 0.47 | 0.40 | 0.36 | 0.50 | 0.12 | . |
| *Ap5n* | 0.70 | 0.38 | 0.37 | 0.49 | 0.41 | 0.41 | 0.47 | 0.23 | 0.04 |

### Monophyly of *Ap2n* mitochondrial clades

The NJ tree based on our genetic distance for automicts, which accounts for recombination and null alleles, is represented in Fig. 7. Individuals that cluster together in this tree often had the same mitochondrial haplotype group (*Ap2n-kaz* or *Ap2n-urm*). In other words, nuclear genetic distance between pairs of individuals with mt-2n*k* and mt-2n*u* mitochondrial haplotypes was larger, on average, than that between pairs of individuals with the same mitochondrial haplotype group (Fig. 7; distance d(*Ap2n-kaz*-*Ap2n-urm*)=9.23; d(*Ap2n-urm*-*Ap2n-urm*)=8.51; d(*Ap2n-kaz*-*Ap2n-kaz*)=8.22). This association between mitochondrial haplotype and nuclear background was highly significant: among 10000 randomizations of the dataset, the estimated genetic distance between pairs of individuals with the same mitochondrial haplotype was never lower than observed genetic distances of 8.51 or 8.22 (i.e. *P<0.0001*). We identified 13 individuals with a potential mismatch between mitochondrial haplotype and nuclear background (arrows in Fig. 7). Importantly, the mitochondrial haplotypes of these individuals were neither unique nor shared with sexual species (*Aurm*, *Akaz*), but rather corresponded to major *Ap2n* haplotypes. Moreover, based on the bootstrap analysis, the total length of the NJ tree was not significantly lower than of a tree with forced monophyly of *Ap2n-urm* and *Ap2n-kaz* (i.e., without mito-nuclear discordance, *P*=0.295). Hence, we could not rule out the hypothesis that the mito-nuclear discordance patterns occurred by chance.

**Figure 7.**
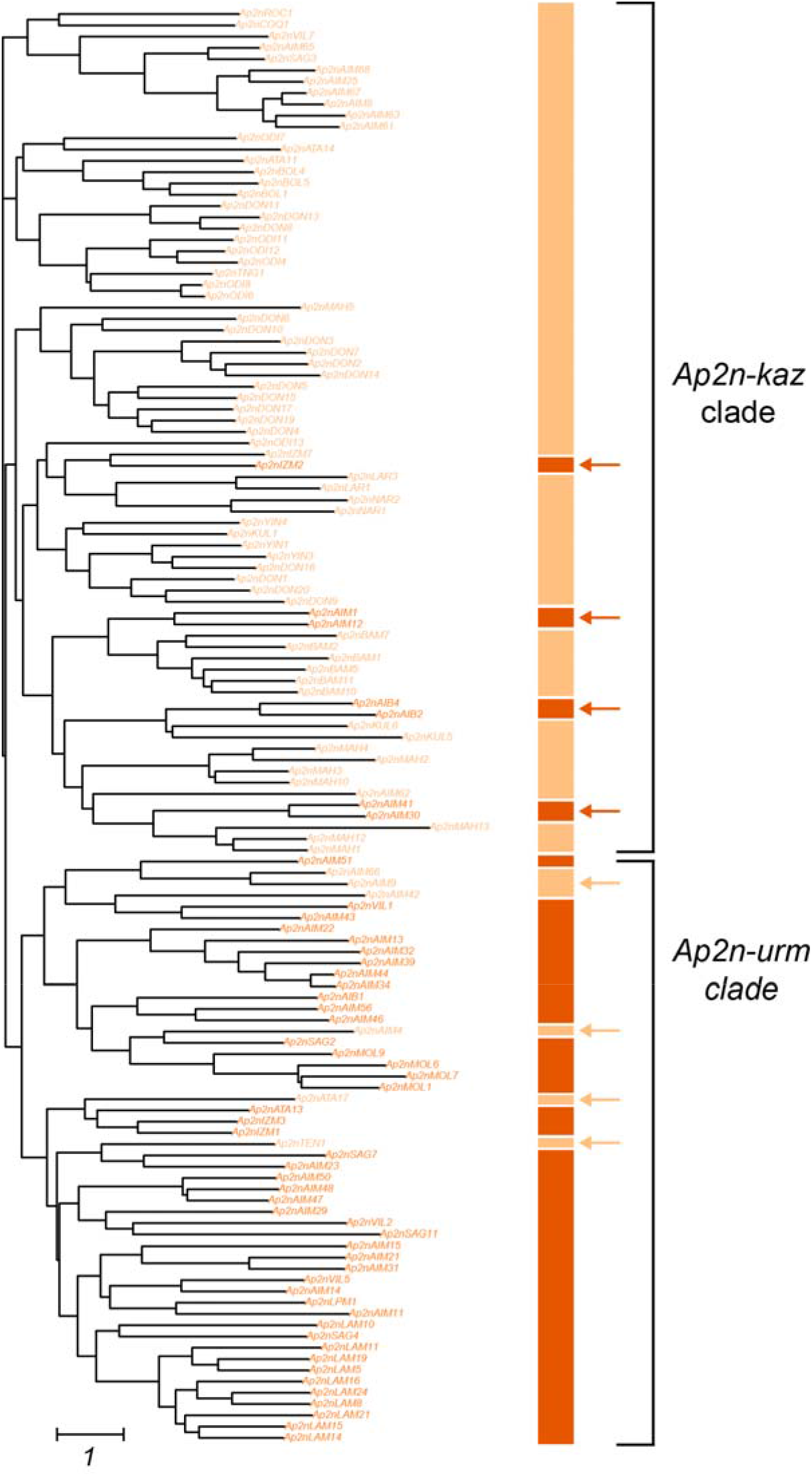
Neighbor-joining tree between pairs of Ap2n individuals based on our nuclear genetic distance for automicts. Individual labels are colored based on mitochondrial haplotypes. Square brackets indicate the two asexual clades Ap2n-urm and Ap2n-kaz based on the match between nuclear and mitochondrial data. Horizontal arrows indicate the few individuals or groups of individuals with mito-nuclear discordances. The association between mitochondrial haplotype and nuclear background is highly significant. Bootstrap values could not be computed due to a low number of markers (note this topology does not fit the data significantly better than the topology where Ap2n-urm and Ap2n-kaz form monophyletic groups, see Results for details).

### Evolutionary origin of *Ap2n-kaz* and *Ap2n-urm*

To compare the different scenarios for the evolutionary origin of *Ap2n-kaz* and *Ap2n-urm*, we used the average pairwise genetic distance between *Ap2n-kaz*, *Ap2n-urm*, *Akaz*, *Aurm* and *Asin* (Table 3). The scenario with the emergence of *Ap2n* lineages through one hybridization and one backcross event (Fig. 3D) had the highest support (Table S2). According to this scenario, *Ap2n-urm* first arose through a hybridization event between an *Akaz* male and an *Aunk* female, carrying a mt-*urm*. *Ap2n-kaz* arose through a backcross between an *Akaz* female and an *Ap2n-urm* rare male. All other scenarios could be ruled out (ΔAICc > 2.5, Table S2), including scenarios involving a spontaneous origin, two independent hybrid origins, or a different order of events (e.g. *Ap2n-kaz* being the F1 and *Ap2n-urm* the backcross).

### Evolutionary origin of *Ap3n*,*Ap4n* and *Ap5n*

For *Ap3n*, average genetic distances inferred from the scenario involving an *Aurm* paternal origin were significantly lower than other genetic distances (*P*<2e-16, Table S3, Fig. 8A). For *Ap4n*, average genetic distances inferred from the scenario involving an *Ap2n-kaz* paternal origin was slightly better than that with an *Ap2n-urm* paternal origin (*P*=0.029, Table S3, Fig. 8B). For *Ap5n*, average genetic distances inferred from the scenario involving an *Atib* paternal origin were significantly lower than genetic distances from other scenarios (*P*<0.0003, Table S3, Fig. 8C).

**Figure 8.**
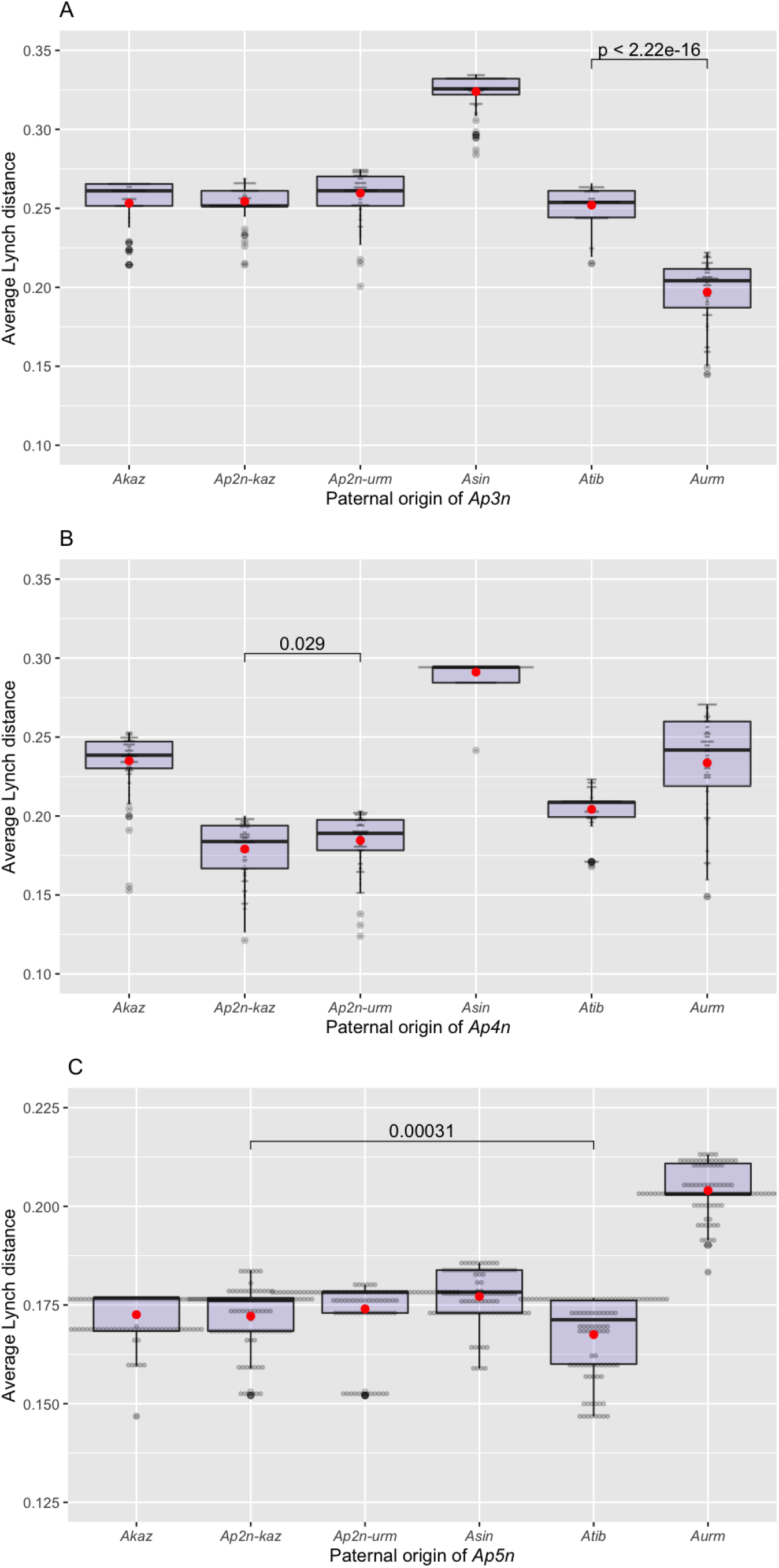
Comparison of the different scenarios for the paternal origin of (A) Ap3n, (B) Ap4n and (C) Ap5n. For each ploidy level and each paternal origin, we estimated Lynch genetic distance across 9 microsatellite loci between 10,000 simulated hybrids and real individuals of our dataset with the corresponding ploidy level (Ap3n: n=16, Ap4n: n=23, Ap5n: n=15). The horizontal line and the red dot respectively represent the median and the mean of the 100 synthetic hybrids with the smallest genetic distance. See main text for details.

## Discussion

### General implications for the study of the origin and evolution of asexuals

Our study proposes new ways to investigate the origin of asexual lineages. We try to address the five hurdles that frequently face such studies, using approaches that can likely be used in other asexual taxa than *A. parthenogenetica*. First, when considering potential hybrid or contagious origins of asexuality, our study showcases that the genealogy of asexual groups might include several origins and/or several nested hybridization events. Hence, the origin of asexuality gene(s) should be clearly distinguished from the origin of asexuals clades that carry these genes. Indeed, we find a single origin of asexuality gene(s) in *A. parthenogenetica*, followed by several nested hybridization with different sexual species. Second, heterogeneity in LOH within and across chromosomes should be carefully considered in automictic asexuals. In particular, with central fusion automicts, which are frequent, heterozygosity can be high close to the centromere, but decreases further away from the centromere. Thus, LOH creates a heterogeneity in coalescence times both among markers along each chromosome and between asexuality gene(s) and the rest of the genome. The methods developed in this study should improve genetic distance estimates between automictic asexual lineages by explicitly accounting for this heterogeneity. In contrast to a widespread hypothesis regarding the evolution of asexuality in ZW lineages (Engelstädter 2008), we show that this sex-determination system does not represent a major obstacle to the evolution of asexuality, provided that recombination is low enough (or quickly evolved to low levels; Boyer et al. 2021). In addition, in these systems, the production of rare ZZ males may allow for the production of new asexual lineages through contagious asexuality. Third, our results support the hypothesis that polyploidy is often a consequence rather than cause of asexuality (Neiman et al. 2014). In addition, our findings suggest that asexual groups with odd ploidy levels (3n, 5n) can reproduce through automixis. This implies that, despite the uneven number of chromosomes, pairing between some homologous chromosomes does occur, but that this pairing does not prevent the formation of an unreduced egg. Fourth, the occurence of rare sex can greatly impact the genomic evolution of asexual lineages by increasing their diversity. The genomic consequences of these events depend on whether they occur within an asexual lineage, between different asexual lineages or between asexual lineages and related sexual species. Rare sex events within an asexual lineage can only be detected when there is some genetic diversity within asexual populations. Importantly, selection might favor a particular asexual lineage within a population and may locally erase the genetic footprints of rare sex events, increasing the difficulty to detect them. Rare sex between different asexual lineages or between asexual lineages and related sexual species, can be detected through admixture tests or through cyto-nuclear discordances. The new method developed here to test for cyto-nuclear discordances assesses whether groups defined based on mitochondria are monophyletic at the nuclear level. This method should be widely applicable in other systems. Fifth, although our sampling was as exhaustive as possible, our results point to particular geographic locations towards which sampling efforts of sexual relatives should be directed in the future in order to refine the estimates of the age for the origin of asexuality in *Artemia* (see below).

### Origins of asexuality in *Artemia*: overview

The most parsimonious scenario to summarize our main results is presented in Fig. 9 and indicates that that both diploid and polyploid asexual *Artemia* harbor nuclear genomes that are admixed between the nuclear genomes of *Akaz* and one or several of the other sexual species. Additional sampling and more genetic data may reveal that some specific crosses are actually more complex and/or that other scenarios need to be statistically compared, e.g. scenarios involving serial backcrosses.

**Figure 9.**
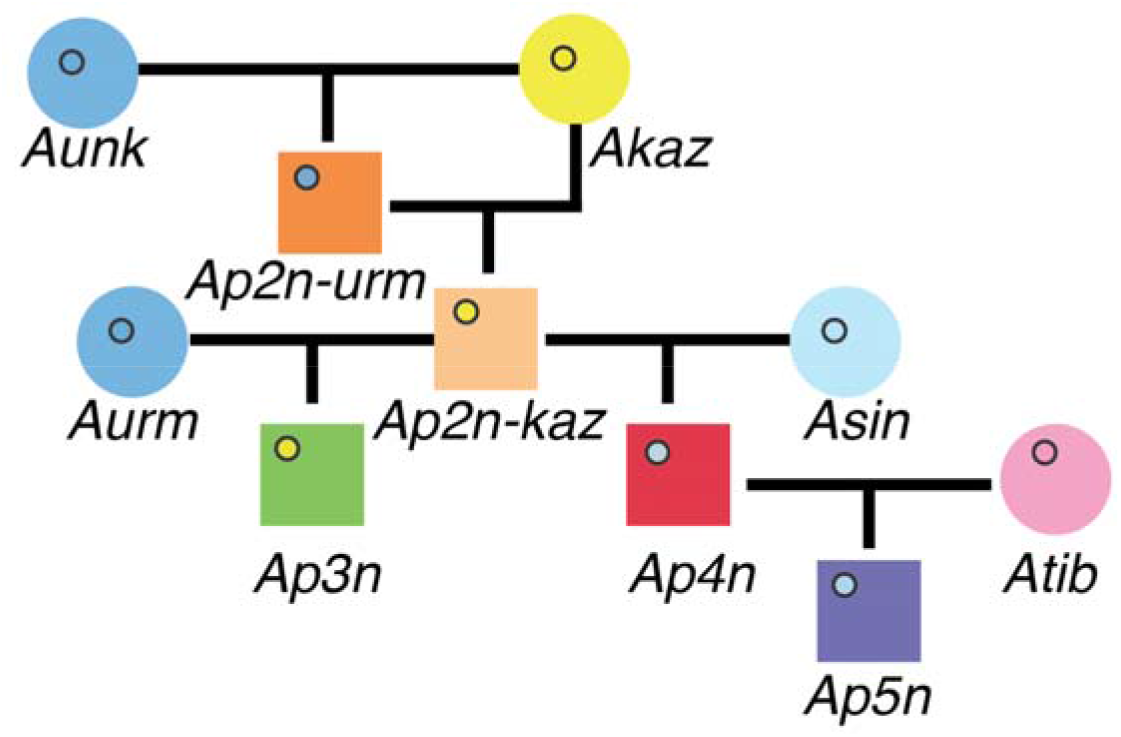
Most likely scenario for the origin of diploids (Ap2n-kaz and Ap2n-urm) and polyploid (Ap3n, Ap4n, Ap5n) asexual Artemia lineages. Briefly, Ap2n-urm likely arose through hybridization between a Aunk female from an unknown species or population (carrying a mitochondrion close to mt-urm) and an Akaz male (note that Ap2n-urm could also derive from a series of backcrosses with the parental species, and there is currently no data to sort this out without knowing this unknown species/population). Ap2n-kaz likely arose as a backcross between an Ap2n-urm rare male and an Akaz female. Ap3n likely arose through the fertilization of Ap2n-kaz female by an Aurm male. Ap4n likely arose through the fertilization of Asin female by an Ap2n-kaz male, followed by an endoduplication in this diploid offspring or one of its descendants. Finally, Ap5n likely resulted from the fertilization of an Ap4n female by an Atib male. Sexual and asexual taxa are represented by filled circles and filled squares, repectively. The filled circle within each symbol represents the mitochondrial haplotype, which shows the maternal origin of each asexual lineage.

The relatively low diversity in mitochondrial haplotypes suggests that contagious asexuality is present but rare in *Artemia* (or leads most of the time to unfit hybrids). Despite the relatively frequent occurrence of rare males in *Ap2n* (Maccari et al. 2013*b*), our data suggest that contagious asexuality via rare males has occurred twice, leading to the *Ap2n-kaz* and *Ap4n* lineages with no evidence for further successful events of contagious asexuality. Indeed, the diversity of mitochondrial haplotypes within each of the main asexual clades is limited and these haplotypes are not intermingled with haplotypes found in sexual lineages. Mutation (rather than backcrossing with unsampled sexual populations) thus seems the most parsimonious explanation for the observed haplotype diversity within each main lineage of asexual *Artemia.*

In addition, each mitochondrial haplotype is only a few mutational steps away from the mitochondria of the different sexual species (*Akaz*, *Aurm*, *Asin*), which suggests that all extant asexual lineages are more recent than previously thought (less than 80,000 years old compared to more than 3 and 0.84 Mya; Perez et al. 1994; Baxevanis et al. 2006; Eimanifar et al. 2015). This estimate represents the age of the most recent common ancestor of all these lineages (i.e. the oldest extant lineage *Ap2n-urm*, from which all other lineages likely arose through subsequent hybridization events). Importantly, the loci that cause asexuality may be much older than our rough estimate of the age of their common ancestor. Indeed, we cannot exclude that all original asexual lineages went extinct, leaving only recently derived ones. This issue can only be solved by comparing the divergence of the region(s) directly associated with asexuality with that of the rest of the genome (Tucker et al. 2013). We discuss the implications of these findings for *A. parthenogenetica* asexuality before discussing each of these events in details.

### Diploids and polyploids may all be automicts

Our findings strongly suggest that all asexual *Artemia* probably derive from a single original hybrid ancestor. The same asexuality genes are therefore very probably shared among all, diploid and polyploid *A. parthenogenetica*. We found that the apparent polyphyly of the group (Baxevanis et al. 2006) results from a history of nested crosses between asexuals and sexual relatives.

Automixis in *Ap2n* has been a major source of confusion throughout one century of cytological observations. Recent genetic data have clarified this debate, and support the conclusion that *Ap2n* reproduce by central fusion automixis (Nougué et al. 2015*b*; Boyer et al. 2021). In contrast most of the literature on polyploids is not controversial and claim that they are apomictic (Brauer 1894; Artom 1931; Barigozzi 1944, 1974). Reproduction of polyploids via an automictic process that would involved recombination has been ruled out based on three types of observations. First, polyploids do not seem to produce rare males (Goldschmidt 1952; Metalli and Ballardin 1970; Chang et al. 2017). Second, each ploidy level shows high heterozygosity, but little clonal diversity (Abreu-Grobois and Beardmore 1982; Zhang and King 1992; Maniatsi et al. 2011). Third, cytological observations have been claimed to refute automixis (due to the failure to observe meiosis, the authors inferred that sister chromatids separate through a mitosis; (Barigozzi 1974)).

Asexuality is likely to have the same genetic determinism in diploid and polyploid asexual lineages, cince they share a common asexual ancestor. If true, this means that the distinction between *Ap2n* automicts and polyploid apomicts may be erroneous. Beyond shared ancestry, several lines of evidence support that polyploids may well, in fact, have the same reproductive mode as diploids.

First, the cytological evidence is not as clear-cut as often reported. This confusion relies on different definitions of automixis by cytologists and geneticists. The former use the fusion of meiotic products as a criterion, while the latter use the genetic consequences of modified meiosis as a criterion (Asher 1970; Nougué et al. 2015*b*; Svendsen et al. 2015). Most cytologists did not consider the possibility that automixis could occur through the abortion of one of the two meiotic steps. In fact, this aborted meiosis has been described by Goldschmidt (1952) in *Ap3n* and *Ap5n* polyploids. She did not observe any fusion of meiotic products, but a brief synapsis and the production of a polar body. Furthermore, she observed that the number of elements (bivalent or univalent) drops during diakinesis and increases afterwards to reach univalent number at metaphase. These observations refute the occurrence of apomixis. They show that meiosis I is aborted at the end of prophase I (with bivalents being separated within the oocyte, but the resulting univalents not being distributed to different different daughter cells and instead re-aligned at the equatorial plate) and jumps directly to metaphase II. Note that in *Ap3n* and *Ap5n*, this brief pairing of homologues can easily occur despite the odd number of chromosomes (one chromosome simply stays unpaired during prophase I), as observed in other animal species (e.g. Christiansen and Reyer 2009). This meiosis modification corresponds to the reproductive mode of *Ap2n* (Nougué et al. 2015*b*; Boyer et al. 2021) and exactly matches “central fusion automixis” as defined by geneticists. According to Asher (1970), the suppression (or abortion) of meiosis I is genetically equivalent to automixis with central fusion as defined cytologically (i.e., where fusion actually occurs), while supression or abortion of meiosis II is cytologically equivalent to automixis with terminal fusion. Other authors refer to the suppression (or abortion) of one of the meiotic divisions as “meiotic apomixis” (e.g. Archetti 2010). However, this term refers both to the suppression of meiosis I and the suppression of meiosis II, whose outcome differ genetically when recombination is present. For this reason, we prefer the term “central fusion automixis” in the large, genetic sense, noting that it includes meiotic apomixis with supression of meiosis I.

Second, if polyploids are automicts, they may occasionally lose heterozygosity because of recombination, as observed in *Ap2n* lineages (Boyer et al. 2021). However, there are good reasons to expect that, in polyploids, homeologous chromosomes (i.e. pairs of chromosomes derived from the two parental species of allopolyploids; Glover et al. 2016) are more divergent than non-homeologous chromosmes (i.e. pairs of chromosomes derived from only one of the two parental species) and that they will pair much less frequently, which likely results in almost no LOH between homeologs. This also likely limits recombination-generated genetic variation in polyploids. In addition, it might also drastically reduce the rate of rare male production in polyploids. In ZZW triploids, Z chromosomes are likely to preferentially pair and recombine (leaving the W unpaired), before all chromosomes realign on the equatorial plate and meiosis II starts. Similarly, in ZZWW tetraploids or ZZZWW pentaploids, W non-homeologous chromosomes are likely to preferentially pair together (and even if Z and W homoeologous chromosomes would pair, two subsequent LOH would be require to produce ZZZZ or ZZZZZ males). Furthermore, as in all polyploids, there is probably a strong selection pressure to reduce the number of cross overs to avoid interlocking cross over events among different pairs of chromatids (Lenormand et al. 2016), which may reduce recombination in polyploids and reinforce this apparent apomictic-like reproduction. As we often observe only two alleles in polyploids, recombination rates between non-homeologous chromosomes might be small but greater than the mutation rate. Hence, the absence of rare males in polyploids is not necessarily an argument against central fusion automixis. Polyploids *A. parthenogenetica* may therefore not be apomicts. They may reproduce by central fusion automixis, but polyploidy and the absence of pairing between homoeologous chromosomes would make this reproductive mode genetically very close to apomixis. This interpretation is open to further tests, but it would explain why polyploids have apomictic-like reproduction, while being derived from automictic lineages as suggested by our study.

### Hybrid origin of *Ap2n-urm* and contagious origin of *Ap2n-kaz* via an *Ap2n-urm* rare male

Our best scenario is that *Ap2n-urm* likely arose through hybridization between a female from and *Aunk*, an unknown sexual species with a mitochondrial haplotype close to that of *Aurm.* It is quite likely that *Aunk* would be related to *Akaz* and *Aurm*, and therefore present in Crimea or in Central Asia, if not extinct. Interestingly, one sexual population from Crimea could be a good candidate. It is currently described as an *Aurm* population (Abatzopoulos et al. 2009; Maccari et al. 2013*a*), but, unexpectedly, its mitochondrial haplotype is closer to that of *Ap2n-urm* than to that of *Aurm* (1 versus 3 mutational steps, respectively, in the network on Fig. 5). Unfortunately, we could not obtain this sample for this study.

According to our best model, the second *Ap2n* group originated through a backcross between a rare male of this first lineage and an *Akaz* female. The genome size of *Ap2n-kaz* is consistent with this scenario. The backcross might have occurred almost immediately after the first occurrence of *Ap2n-urm*, as rare males may be produced at a higher rate in young asexual lineages (Boyer et al. 2021) and as both crosses rely on the presence of *Akaz*, which likely has a very limited geographical range, at least today. All *Ap2n* lineages seem to branch from these two major groups, and we did not find firm evidence of any further event of contagious asexuality. Such secondary contagion through rare males would indeed capture new mitochondrial haplotypes from sexual species, which would be easily detected. The rmPAK1 sequence in Fig. 4 from Maccari et al. (2013*b*) might be a rare candidate, but without nuclear data, we cannot further investigate this possibility. Overall, the low diversity among *Ap2n* haplotypes suggests that secondary contagion is probably very rare. The restricted geographical distributions of sexuals in Central Asia and around the Crimean peninsula and their spatial or temporal segregation from asexuals (Mura and Nagorskaya 2005; Shadrin and Anufriieva 2012) might indeed limit the chances of contagion.

Recent experimental evidence (Boyer et al. 2021) has shown that *Ap2n-kaz* may occasionally reproduce sexually. Reproduction between *Ap2n-kaz* and *Ap2n-urm* could perhaps explain some mito-nuclear discordance observed in our data (Fig 7). Alternatively, *Ap2n-kaz* and *Ap2n-urm* might form monophyletic clades and these discordances might be due to our limited number of microsatellite markers. Finally, sexual reproduction might preferentially occur within *Ap2n-kaz* and *Ap2n-urm*, as most populations are composed of individuals from a single clade (Fig. 2). Mitochondrial divergence is low within *Ap2n-kaz* and *Ap2n-urm*, so that we could not test and detect mito-nuclear discordance within each clade with our method. Additional mitochondrial and nuclear data will be needed to better test these different hypotheses.

### Origin by transmission of *Ap3n* via an *Ap2n-kaz* female

All *Ap3n* branch within the *Ap2n-kaz* lineage and show very limited genetic diversity at both the mitochondrial and nuclear levels, despite having a worldwide distribution, from Madagascar to the Mediterranean. This result is in line with that of Maniatsi et al (2011) that found only two different clones across 10 *Ap3n* populations. It contrasts with the large allozyme diversity reported within and among three *Ap3n* populations by Abreu-Grobois and Beardmore (1982). Our samples (NAR, IZM, ANK) belong to a single *Ap3n* lineage that likely arose through a cross between an *Ap2n-kaz* female and an *Aurm* male. When considering samples from previously published studies, we found several mitochondrial haplotypes that are shared between *Ap3n* and *Ap2n-kaz* (Fig. 5 and sup. mat. 9). This might have been caused by improper assignation of ploidy levels of these samples or from the independent origin of several, highly related, but distinct, *Ap3n* lineages. Further sampling, genotyping and careful check of ploidy levels should resolve the matter.

### Contagious origin of *Ap4n* via an *Ap2n-kaz* rare male

We found that *Ap4n* show very limited genetic diversity, consistent with previous studies (Abreu-Grobois and Beardmore 1982; Maniatsi et al. 2011). In the most likely scenario (Fig. 8), *Ap4n* resulted from a standard hybrid cross between reduced haploid gametes from an *Asin* mother and an *Ap2n-kaz* rare male (other scenarios involving unreduced *Asin* gametes are less likely as they require the combination of multiple rare events). Tetraploidy would have occurred through endoduplication in one of these hybrids (or within its descendants). This scenario is consistent with the observation that most microsatellite loci have only two alleles in those individuals. The genome size of *Ap4n* is 7% larger than that of *Asin* and *Ap2n-kaz* combined, which suggest an increase in genome size following polyploidization. Multiple origins of *Ap4n* is unlikely, since we would expect to observe the capture of several mitochondrial haplotypes provided the mitochondrial diversity observed in *Asin*. Historically, the hybridization event probably took place in East Siberia as *Ap4n* carry a mitochondrion that is only found in an *Asin* sample from this area (see sup. mat. 9 for a discussion of mitochondrial diversity within *Asin*). Finally, all the *Ap4n* lineages observed in the Mediterranean area would be a subsample of few successful lineages, which reached a worldwide distribution.

### Origin by transmission of *Ap5n* via an *Ap4n* female

Based on mitochondrial data (Fig. 5), *Ap5n* are derived from an *Ap4n* unreduced gamete fertilized by a male of unknown origin. The microsatellite data agree with this scenario as *Ap5n* genotype are very close to *Ap4n* and suggests an *Atib* paternal origin. However, based on flow cytometry data, the genome size of a hybrid resulting from the fertilization of an *Ap4n* unreduced egg by a reduced *Atib* sperm would be 10.0% larger than the observed genome size of *Ap5n.* This discrepancy may be due to secondary reduction in *Ap5n* genome size or to an incorrect paternal assignment. The second and third best candidates would be *Ap2n*-*kaz* and *Akaz* respectively and the genome size of hybrids resulting from these crosses would better match that of *Ap5n* (2.4% and 3.1% larger, respectively). Additional nuclear data will indicate whether *Atib* paternal origin is the correct inference.

### Implications for the study of the origin of asexual lineages

A more robust dating of the origin of asexuality in *Artemia* and establishing firm scenarios for the origin of the different asexual lineages requires the different sexual species to be more extensively sampled and characterized. *Ap2n-urm* is clearly more closely related to the sexual Crimean population from Lake Koyashskoe than to *Aurm* based on both mitochondrial (Fig. 3) and nuclear data (Abatzopoulos et al. 2009; Sainz-Escudero et al. 2021). This suggests that the Lake Koyashskoe population might be *Aunk*. Additional nuclear genetic data from this and other populations (e.g. Kara-Bogaz-Gol) is required to confirm this hypothesis. Similarly, mitochondrial and nuclear genetic data from sexual populations from the Altai region currently described as *Aurm* (Mura and Nagorskaya 2005; Shadrin and Anufriieva 2012) could help decipher whether one of these populations actually correspond to the *Akaz* population involved in the paternal origin of *Ap2n-urm* and/or the maternal origin of *Ap2n-kaz*. In addition, studying *Asin* populations from Siberia (which are very divergent from *Asin* populations from China; Fig. 5) should shed a light on the maternal origin of *Ap4n*. Finally, studying the diversity of *Atib* populations outside the reference population used to describe this species (Lagkor Co) could help assess the paternal origin of *Ap5n*. Indeed, we confirm that most individuals from Lagkor Co have mt-tib1 mitochondrial haplotypes very close to mt-*kaz* (Fig. 5), whereas all other sexual populations from the Qinghai-Tibet plateau assigned as *Atib* only carry the very divergent mt-tib2 haplotype (as previously found in Maccari et al. (2013*a*)). Our microsatellite data suggests that the Lagkor Co population might be admixed (Fig. S3). Hence, it is possible that the Lagkor Co population has been introgressed by a mitochondrial haplotype closely related to mt-*kaz* (Maccari et al. 2013*a*). Alternatively, all other populations may represent another undescribed sexual species that can hybridize with *Atib* individuals, resulting in a low frequency of the divergent mt-tib2 haplotype in Lagkor Co. Similarly, our microsatellite data suggest that *Aurm* might be admixed between two divergent populations (Fig. S3), which is consistent with the observation of two very divergent ITS1 haplotypes that segregate within *Aurm* (Eimanifar et al. 2014; Sainz-Escudero et al. 2021). Additional genetic data from populations from the Qinghai-Tibet plateau and from Crimea will be required to disentangle among different admixture scenarios and resolve potential mito-nuclear discordances in *Atib* and *Aurm*.

#### Taxonomic implications

The taxonomy of asexual lineages is often ambiguous, as clearly exemplified by our case study. Provided that multiple hybridization events can give rise to different asexual lineages, phylogenetically-defined monophyly is a poor criterion of the taxonomic description of asexual lineages. Asexuality also prevents the use of species delimitation criteria based on interfertility between different taxa. Facing these difficulties, it may be more biologically relevant to focus on the common origin of asexual lineages and their common reproductive mode. In our case study, it makes sense to collectively refer to *A. parthenogenetica* as a relevant taxonomic unit, as they probably inherited the same asexuality gene(s) from their common ancestor. The drawback of this taxonomic approach, in *Artemia* and other groups, may be that a single name hides the diversity of hybridization events that led to the different lineages. As a consequence, a minimal convenient way to designate these taxa would be to distinguish the major groups derived from these crosses, if they can be distinguished. For instance, in *A. parthenogenetica*, five major groups may summarize the major sex-asex transitions: *Ap2n-kaz*, *Ap2n-urm*, *Ap3n*, *Ap4n* and *Ap5n*. Finally, taxonomic issues, lineage history and age, may also be better resolved by directly identifying and studying the genomic regions associated with asexuality (Sandrock and Vorburger 2011; Tucker et al. 2013; Yagound et al. 2020). As our case study shows, sex-asex transitions are quite different from the idealized view of an apomictic mutant arising in a sexual species. Ultimately, a better characterization and understanding of sex-asex transitions represents a pivotal step to refine our theories for the long-term maintenance of sexual reproduction and the extant distribution of sexual and asexual taxa.

## Acknowledgements

We thank Christ Mahieu (Laboratory of Aquaculture & *Artemia* Reference Center), Paco Amat (Instituto de Acuicultura de Torre de la Sal) and Marta Sanchez (University of Sevilla) for providing samples and Emmanuel Douzery (ISEM, University of Montpellier) and Arnaud Estoup (CBGP, INRAE) for advice on phylogenetic and population genetic methods. We thank Marie-Pierre Dubois and the GEMEX molecular platform of CEFE laboratory, and the GENSEQ platform of the Centre Méditerranéen Environnement Biodiversité (LABEX CEMEB, Montpellier, France) for assistance with molecular laboratory work. We also thank Maria Orive and two anonymous reviewers for their useful comments.

