## Supplemental File for "The origin of asexual brine shrimps"

### Supplementary Materials

#### 1 Ploidy characterization

Following Nougué et al 2015, we estimated the genome size of each of 147 individuals using flow cytometry. Briefly, each individual was cut in halves using a razor blade and the first half was kept in 96% ethanol for subsequent COI and microsatellite genotyping. The second half was chopped in a small plastic petri dish, together with a 0.5-mm2 leaf of *Oryza sativa* ssp *japonica* cv Nipponbare in 1mL of modified LB01 nuclei extraction buffer (Nougué et al. 2015*b*). After nuclei homogenization and filtering through a 30-µm Partec filter, we added 5uL of RNAse (final concentration: 50µg/mL) and 40uL of propidium iodide (0.2 mg/mL). After vortexing for 5s at low speed, samples were incubated on ice for 10 minutes. We estimated the nuclear DNA content of each individual based on DNA-PI fluorescence emission at 600–640 nm, using a Partec PAII laser Flow Cytometer (Partec GmbH, Münster, Germany). Histograms were analysed using the Partec FloMax software, which determines peak position and coefficient of variation. We estimated genome size using the mean fluorescence intensity (FL, arbitrary units) of gated data corresponding to first and second fluorescence peaks as follows:

Total genome size = (Second peak FL / First Peak FL) x 0.91 pg,

where 0.91 pg corresponds to the diploid genome size of *Oryza sativa* ssp *japonica* (Uozuet al 1997). For six populations (*Aurm*, *Akaz*, AIM, ODI, BUJ, CIT), we also used the data from 57 individuals from (Nougué et al. 2015*b*).

#### 2 COI genotyping

For each of 336 individuals, we amplified a fragment of the mitochondrial Cytochrome c Oxidase Subunit I (COI) gene using the COI_Fol-F/COI_Fol-R primers (Muñoz et al. 2010). We replicated this step once and twice for respectively 15 and 1 individuals, resulting in 417 sequences. Provided the large number of numts amplifications (see below), we designed shorter and more specific primers: Co1APAR-F(5’-TTTGGAGCTTGAGCAGGAAT-3’) and Co1APAR-R(5’-TGCGGGATCAAAGAAAGAAG-3’). We genotyped 31 additional individuals with these primers: Ap3nANK1, Ap3nANK7, Ap2nBDP1, Ap2nAIM2, Ap2nAIM1, Ap2nAIM5, Ap2nATA11, Ap2nATA13, Ap2nATA14, Ap2nATA15, Ap2nATA16, Ap2nBDP2, Ap2nLAR1, Ap2nLAR3, Ap2nNAR1, Ap2nNAR3, Ap2nNAR4, Ap2nROC1, Ap2nROC2, Ap2nROC3, Ap2nROC4, Ap2nROC5, Ap2nTNG1, Ap2nTEN1, Ap2nTEN2, Ap2nTEN3, Ap2nTEN4, Ap2nTEN4, Ap2nTEN5, Ap2nYIN2, Ap2nYIN3. We used 25-50ng of DNA template, 2.5µl of 10X PCR buffer with 1.5mM of MgCl2, 1 unit of Taq polymerase, 10nmol of dNTPs, 15pmol of each primer. A fragment of 606 base pairs (bp) was amplified by PCR under the following conditions: one cycle of denaturation at 94°C for 3min, and 40 cycles of denaturation at 94°C for 1min, annealing at 53°C for 1min, and extension at 72°C for 2min, with a final extension at 72°C for 15min. PCR products were sequenced from both strands sequenced by Eurofins MWG Operon (Germany). Electrophoregrams were checked by eye using CodonCode Aligner v. 3.5 (CodonCode Corporation, Dedham, MA). We removed 53 sequences with indels. For three and one individuals with two and three identical replicate sequences, we build a consensus sequence. The resulting dataset of 359 sequences was split into two datasets (dataset1: 198 sequences including at least 538 sites in the final alignment and without any ambiguous position, and dataset2 including 161 sequences of low or intermediate quality).

To investigate the potential coamplification by PCR of nuclear pseudogenes with their mitochondrial counterpart, we used two different approaches. We first amplified, cloned and sequenced PCR fragments for 6 individuals among the 400 mentioned above (2 *Ap4n*: Ap4nIZM1, Ap4nTEK3 and 4 *Ap5n*: Ap5nBUJ2, Ap5nCIT6, Ap5nDON2, Ap5nIZM2). For each ligation, 5–10 positive clones were identified by PCR analysis of the plasmid DNA, depending on the success of the cloning. Successful amplification was confirmed by agarose gel electrophoresis, and PCR products were sequenced as mentioned previously. Overall, we recover 26 unique sequences, which can include both numt and mitochondrial sequences. Second, we selected one individual either from different sexual species (*Atib* or *Aurm*) or asexual lineages (*Ap2n-urm*, *Ap2n-kaz*, *Ap3n*, *Ap5n*). To increase the final proportion of mitochondrial DNA within total DNA, we extracted mitochondrial DNA using the mitochondrial DNA isolation kit ab65321 (Abcam, Cambridge, UK) following the manufacturer’s protocol. We then amplified, cloned and sequenced the PCR fragments as above. Assuming no heteroplasmy, the majority sequences for each individual was considered the mitochondrial sequence, while minority sequences were considered as numts. Hence, we recovered six mitochondrial and nine numt sequences. Importantly, the *Ap5n* mitochondrial sequence only matched two (Ap4nIZM1 and Ap5nDON2) of the 26 sequences obtained through the first cloning approach. Provided that *Ap4n* and *Ap5n* are very close based on nuclear data (see main text), we considered these two sequences to be mitochondrial sequences while the other 22 sequences were considered as numts. We removed one mitochondrial sequence which was identical to one sequence of dataset1 obtained from the same individual and one numt sequence with ambiguous positions, resulting in a dataset of 7 mitochondrial sequences and 32 numt sequences (hereafter dataset3).

We also retrieved 804 additional sequences from GenBank (Table S5), most of which have been included in previous studies regarding the origin of asexual lineages (Muñoz et al. 2010; Maccari et al. 2013*b*, 2013*a*; Eimanifar et al. 2014, 2015; Asem et al. 2016). We did not include the 18 haplotypes described in Maniatsi et al. (2011) as an undisclosed number of these shorter sequences were obtained from pools of cysts (i.e. sequences could represent chimeras due to the co-amplification of the mitochondrial sequences of different genotypes). We removed seven sequences with indels and 49 low quality sequences (with ambiguous positions or with less than 538 sites in the final alignment), resulting in a dataset with 748 high quality sequences. The latter was combined with dataset1 and dataset3 into a large dataset of 985 high quality sequences.

#### 3 Numt detection based on polarity changes

To detect potential numts, we tested for drastic changes in the polarity of amino acid residues (Kunz et al. 2019).The reading frame of the alignement of the 985 COI nucleotide sequences was found based on the absence of stop codons in the protein sequences. For each polymorphic position in the COI protein alignement, we considered the majority amino-acid as the ancestral one and the minority amino acid as the derived one. We computed the standardized polarity difference between ancestral and derived amino acids as the difference in their PP1 (principal property 1; Cruciani et al. 2004), as provided in the R package *Peptides* (Osorio et al. 2015). For each of the 985 COI protein sequences, we computed the sum of their standardized polarity differences of their derived amino-acids. Based on the distribution of this sum across sequences, we defined 0.1 as the threshold above which sequences were considered as potential numts (Fig. S1).

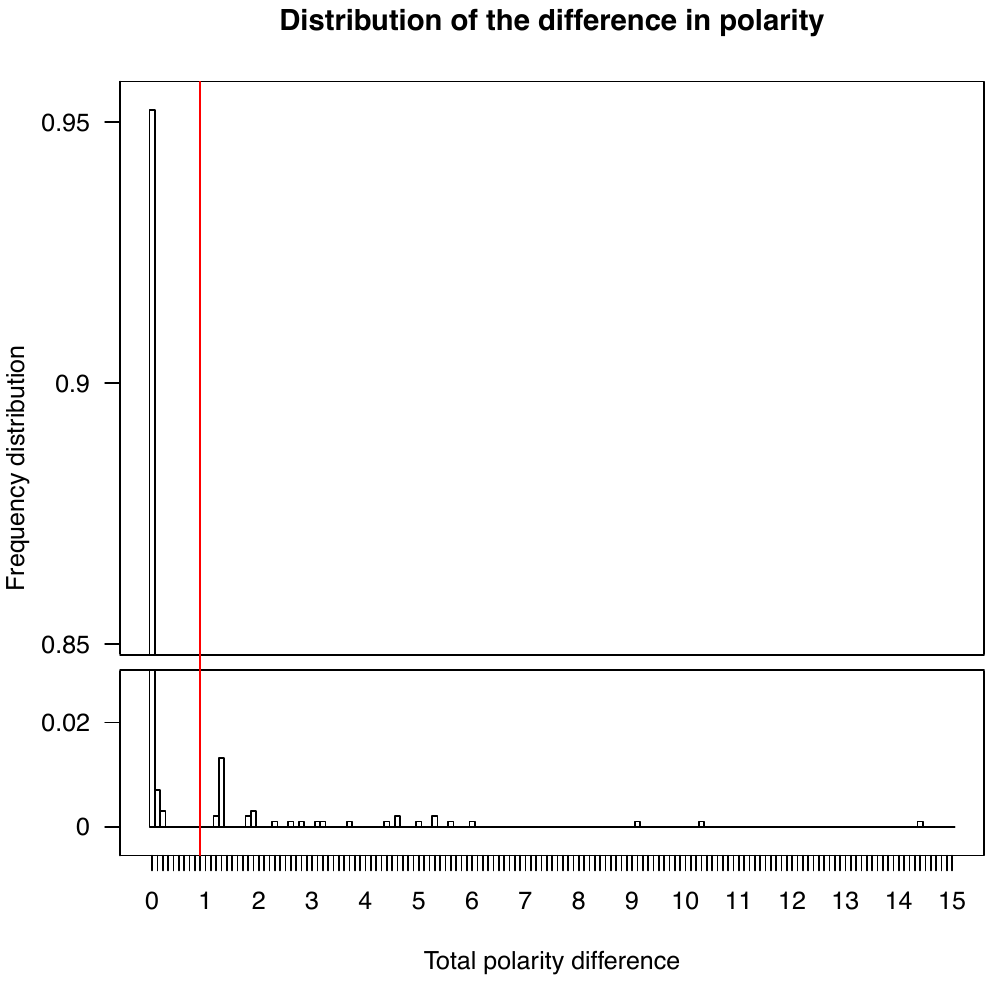

**Fig. S1. Distribution of the sum of polarity differences of each of the 985 COI protein sequences in our dataset.** The red vertical red bar represents as threshold of a polarity difference of 0.1 above which sequences were considered as numts. The horizontal axis is not to scale in order to visually represent low-frequency sequences with extremely high polarity differences.

#### 4 Automatic chimera detection

We developed a new test to automatically detect chimeras. First, we collapsed the 985 sequences into a reference set of 230 unique haplotypes. Second, we compared each of those 230 focal sequences to all pairs of remaining sequences in the dataset. For each pair, we count the number of mutations in the focal sequence which are not found in the pair. We note dB this baseline distance (expressed in number of mutations). Third, for each focal sequence, we selected the best pair, i.e. the one with the lowest dB (or several best pairs if they all have the same lowest dB). This best pair (or these best pairs) represents the best candidates for the two ‘true’ sequences leading to a chimera. For each best pair, we computed the number of mutations to each of these two ‘true’ sequences, noted d1 and d2. Finally, we computed a chimera score dchim as the minimum of d1-dB and d2-dB. We applied a cutoff of dchim=2 to identify likely chimeras. This means that we consider that a sequence is a chimera when it harbors a least two mutations compared to its closest sequence that also appear in another unrelated sequence of the dataset. Intuitively, one sees that a specific mutation at a given position may occur in two unrelated sequences, but there is a very low chance that this rare event occurs twice or more. Let’s consider an example using “rmKUJ1” (Maccari et al. 2013*a*) as a focal sequence. It has a dB score of zero with the pair of sequences “rmPAK1” and “ATHAY5” in the reference set. However, it is distant to both of them with d1 =5 and d2 = 34, respectively. Hence, rmPAK1 has a chimera score of 5. In other words, rmPAK1 is the closest sequence to rmKUJ1, differing by five mutations. However, those five mutations are all found together in another unrelated sequence in the dataset (ATHAY5, Fig. S1). Hence, it is much more likely that rmKUJ1results from the coamplification of rmPAK1 and ATHAY5. The unlikely alternative woud be that five specific mutations occurred independently in ATHAY5 at the exact same positions.

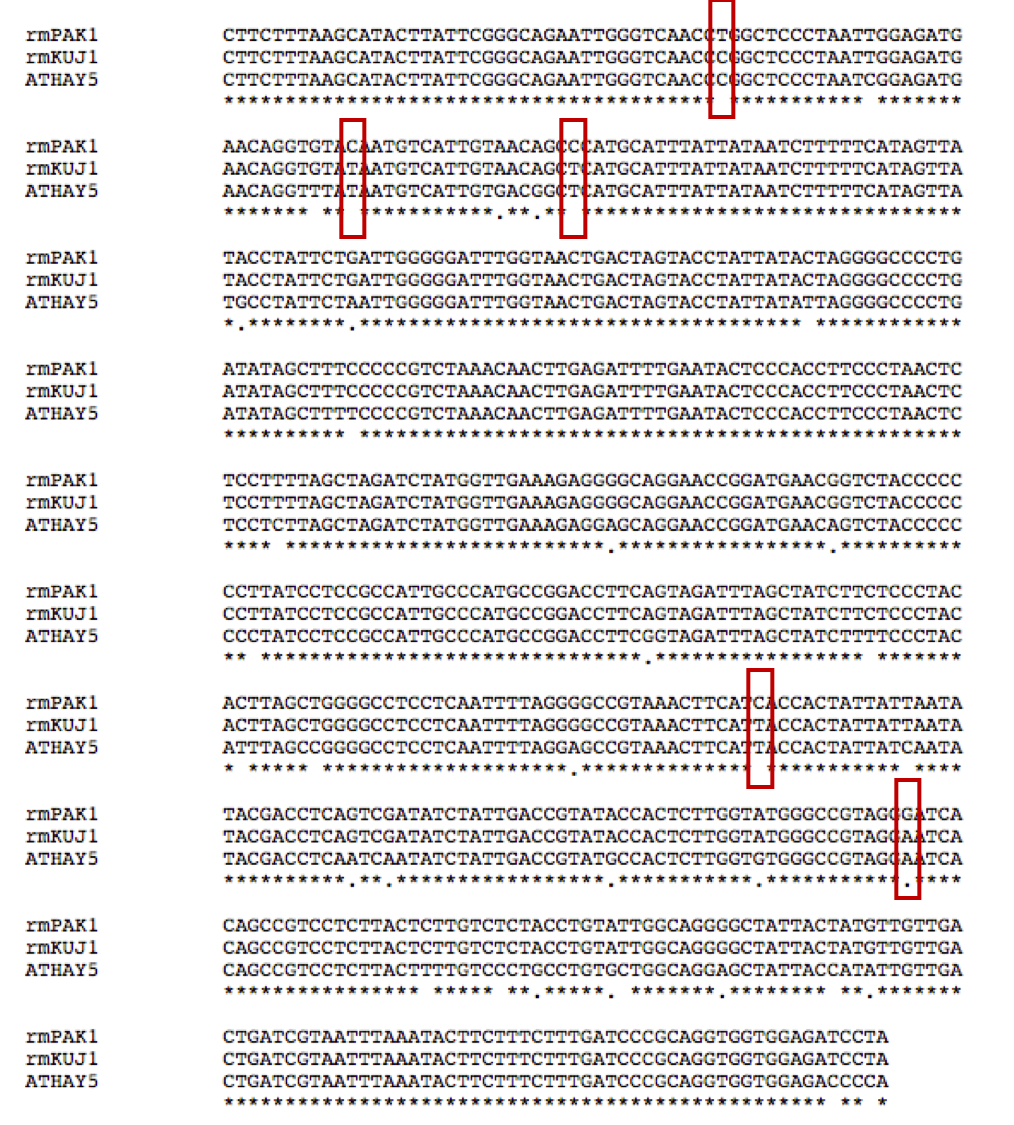

**Figure S2. Alignment of the three sequences rmKUJ1, rmPAK1 and ATHAY5.** Red rectangles indicate the five mutations that differ between the two *Ap2n* sequences rmKUJ1 and rmPAK1. These five mutations are also found in an unrelated *Atib* sequence, ATHAY5.

#### 5 Microsatellite genotyping

For each of the 432 individuals, genomic DNA was extracted using Sigma Extraction Solution (Sigma-Aldrich, St Louis, MO, USA). We use the panel of 12 microsatellite markers described in (Muñoz et al. 2008; Nougué et al. 2015*a*). After PCR amplification, we mixed between 10 and 15 ng of the final product with 15 µl of HI-DI TM formamide (Applied Biosystems) and 0.2 µl of GeneScan-500 LIZ size standard. Fluorescent dye-labeled alleles were detected on an ABI PRISM 3130xL DNA Analyzer (Applied Biosystems) at the LabEx CeMEB sequencing platform (Montpellier, France) and analyzed using GeneMapper v. 3.7 (Soft Genetics). For six populations (*Aurm*, *Akaz*, AIM, ODI, BUJ, CIT), we also used the data from 57 individuals from Table 2 in Nougué *et al.* (Nougué et al. 2015*b*).

#### 6 Detection of admixture in sexuals

Assuming a standard GSM mutation model for microsatellites, a sexual population that is admixed between two sexual source populations is expected to have a larger variance in microsatellite allele size than its two parental populations (provided heterozygosity is roughly the same across populations; Estoup et al. 2016). We examined the proportion of heterozygous loci and the variation in the average difference in allele sizes within individuals from each sexual species taxon (*Aurm*, *Akaz*, *Atib*, *Asin*). For each individual, we computed the proportion of loci that were heterozygous. We also computed the number of mutational steps between the two alleles of each individual at each microsatellite marker (e.g. for a dinucleotide microsatellite, a difference of two base pairs represents one mutational step), which we averaged across the 12 microsatellite markers for each individual separately. The distributions of proportion of heterozygous loci and of the average difference in allele sizes within individuals are represented in Fig. S3. Heterozygosity is lower in *Atib* and *Asin* than in *Akaz* and *Aurm* (Fig. S3A), which is likely due to the presence of null alleles. In contrast, the variance in average difference in allele sizes within individuals is smaller in *Akaz* and *Asin* (YIM population) than the variance observed in *Aurm* and *Atib* (Fig. S3B). This suggests that the latter two populations might be admixed.

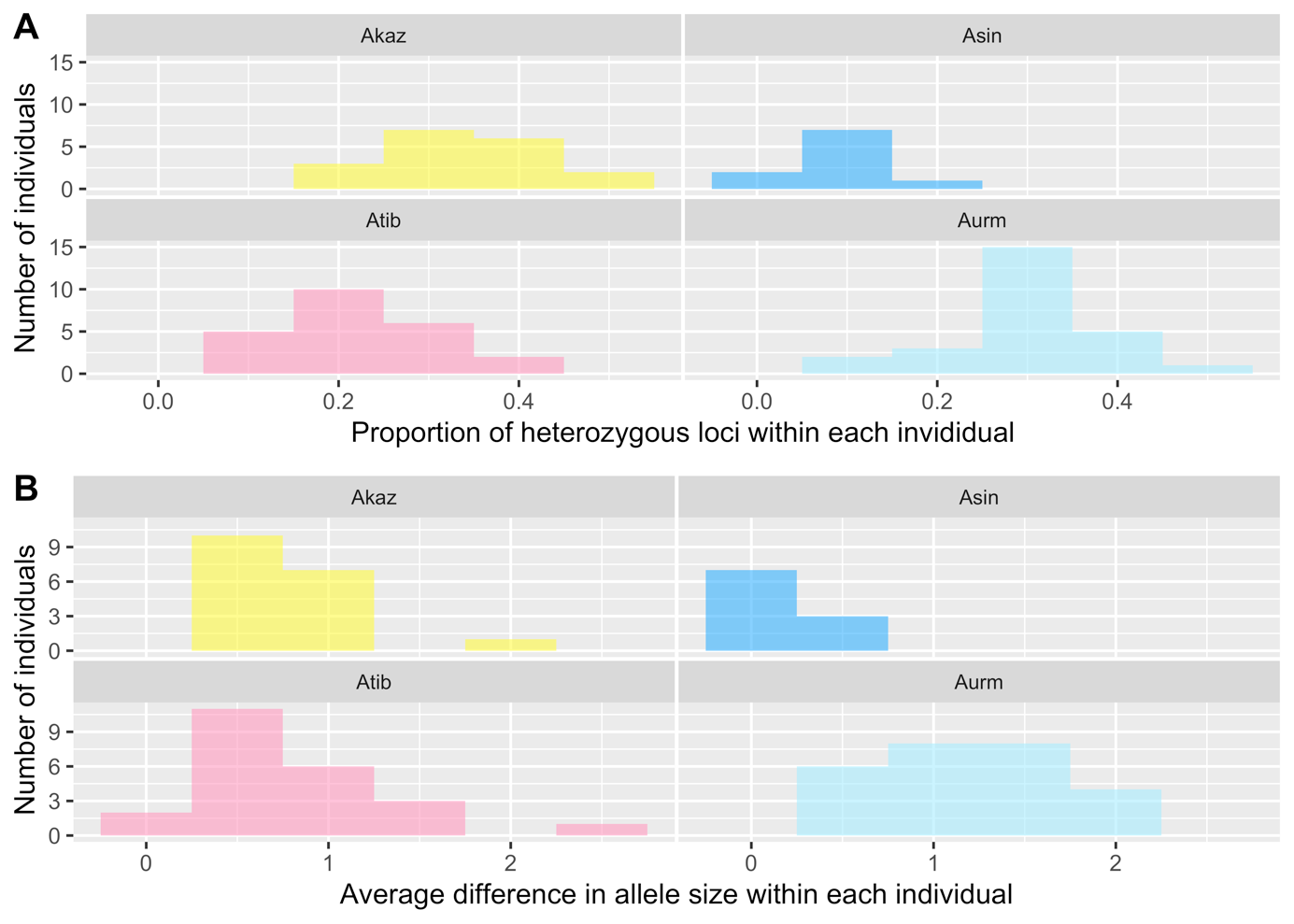

**Figure S3. Distribution of (A) the proportion of heterozygous loci and (B) the average difference in allele size (average number of mutational steps) within individuals from each sexual species.** For *Asin*, only the YIM population was considered.

#### 7 Genetic distance in *Ap2n* automicts

We developed a new genetic distance measure for automicts that accounts for the different possible events producing genotypic variation (mutation at rate *µ* and LOH at rate *r*). Two diploid genotypes are connected by different paths that correspond to different events. We develop a measure that is a proxy for the time length of the path (or time length averaged across different possible paths, according to their relative probability of occurrence). When one event is required along a path, say a mutation, the average time length of the path is simply the inverse of the rate of the occurrence of the event (e.g. 1/µ). When two successive events are required, say mutation and LOH, the time length is the sum of the time to the first plus time to the second event (e.g. 1/µ + 1/*r*). When two events are competing (say mutation or LOH, whichever comes first), the average time length of the path is the inverse of the sum of the rate of occurrence of the competing events (e.g. 1/(*µ*+*r*)). Table S1 below list all possible cases and the corresponding path length.

Because of the possible occurrence of null alleles, several paths need to be considered, reflecting the different combination of ‘true’ genotypes being compared. For instance, individuals scored as homozygous for A allele may be either “AA” or “A∅” (where ∅ denotes a null allele). The latter is expected to be more frequent if the locus has a higher frequency of null allele. However, the frequency of null alleles is difficult to estimate in automictic populations, as the traditional estimation method based on Hardy-Weinberg assumption (Chapuis and Estoup 2007) is likely to be violated.

To address this issue, and for each locus independently, we compute *Q*, the probability that an individual with observed allele A has genotype A∅ (rather than AA). To compute *Q*, we assume that there is a single type of null allele (as in methods used for sexuals), and we use two independent observations: the observed frequency *R* of ∅∅ individuals at this locus, and the excess or deficit of heteroyzotes *F_is_* at this locus (noted *F* in equations below), estimated from another population dataset (Nougué et al. 2015*b*). The frequency of null allele at this locus, *p*, can be directly obtained, knowing *R* and *F,* by solving the equation:

$R=p^{2}+p\left( 1-p \right)F$.

(1)

*Q* is then simply expressed as the proportion of A∅ among A∅ + AA individuals:

$$Q=\frac{2p\left( 1-p \right)(1-F)}{{(1-p)}^{2}+p\left( 1-p \right)F+2p\left( 1-p \right)(1-F)} ,$$

(2)

which is directly estimated knowing *p* and *F*. This estimate can be improved by considering the uncertainty in the estimation of *R* and *p*. Noting *k* the observed number of ∅∅ individuals scored, and *n* the total number of individuals scored, we can average $Q$ weighting Eq. (2) by the likelihood function corresponding to these (*n*, *k*) observations:

$$Q=\int_{0}^{1} \left( n+1 \right) B\left( n,R;k \right)Q^{*} dR ,$$

(3)

where $B\left( n,R;k \right)$ is the probability mass function of the binomial distribution *B*(*n*, *R*) for *k* success among *n* trials, and where $Q^{*}$ is the expression of $Q$ as provided by Eq (2), but where *p* is expressed as the solution of Eq (1). The (*n*+1) factor normalizes the integral and is equal to $1/\int_{0}^{1} B\left( n,R;k \right) dR$. In practice, for each locus, we estimated $Q$ using the average *F_is_* from two European *Ap2n* populations (Nougué et al. 2015*b*), while accounting for the uncertainty associated with the estimation of *R* in our data (*n* = 299 *Ap2n* individuals sampled).

#### 8 Evolutionary origin of *Ap3n*, *Ap4n* and *Ap5n*

We compare different scenarios regarding the paternal origin of *Ap3n*, *Ap4n* and *Ap5n* ancestors. Based on mitochondrial data, we assumed that *Ap3n* originated from the fertilization of an unreduced *Ap2n-kaz* egg by the haploid sperm of a male from either *Akaz*, *Ap2n-kaz*, *Ap2n-urm*, *Aurm*, *Atib* or *Asin*. To form the genotype of *Ap3n* synthetic hybrids, we ramdomly sampled an *Ap2n-kaz* individual and a synthetic haploid sperm from a potential father from each potential species.

Based on mitochondrial data, we assumed that *Ap4n* originated from the fertilization of a reduced *Asin* egg by the haploid sperm of a male from either *Akaz*, *Ap2n-kaz*, *Ap2n-urm*, *Aurm*, *Atib* or *Asin*, followed by an endoduplication. To form the genotype of *Ap4n* synthetic hybrids, we ramdomly sampled an *Asin* individual as a potential mother and sampled a random synthetic haploid ovum. We then randomly sampled an individual from each potential paternal species as a potential father and sampled a random synthetic haploid sperm.

Based on mitochondrial data, we assumed that *Ap5n* originated from the fertilization of an unreduced *Ap4n* egg by a haploid sperm of male from either *Akaz*, *Ap2n-kaz*, *Ap2n-urm*, *Aurm*, *Atib* or *Asin*. To form the genotype of *Ap5n* synthetic hybrids, we ramdomly sampled an *Ap4n* individual and a synthetic haploid sperm from a potential father from each potential species.

To randomly sample haploid sperm (ovum) genotypes in each simulation, we used the genotype of the randomly sampled father (mother) at each of the 12 microstatellite markers. For each marker and when the individual was heterozygous, we randomly sampled one of the two alleles. When the individual was homozygous, we sampled either the observed allele or a null allele. Let~~’s~~ *p* and *p_∅_* be the frequencies of an allele A and a null allele at a given locus. Assuming the existence of a single null allele, apparent homozygotes [AA] can either be real homozygotes with frequency $p^{2}$ or A∅ heterozygotes with frequency ${2 p p}_{\emptyset}$. Double null homozygous have frequency $p_{\emptyset}^{2}$. Assuming Hardy-Weinberg equilibrium, *p* can be computed as $\sqrt{{freq}_{[AA]}+p_{\emptyset}^{2}}-p_{\emptyset}$.

**9 Comparisons with previous studies**

Relationship between *Ap3n* and *Ap2n-kaz*

Some of our *Ap3n* haplotypes were identical to an APD05 haplotype found in supposedly diploid individuals from Narte (Albania) (Muñoz et al. 2010; Maccari et al. 2013*a*) and in uncharacterized individuals from Izmir (Turkey; CAM-2 Eimanifar et al. 2014, 2015). As we used the same cyst samples as these studies, these samples are likely to be triploids. The population from Narte has been described as *Ap2n*, but we found that it is a mixture of both *Ap2n* and *Ap3n*. Other individuals with an APD05 haplotype, and described as *Ap2n,* originated from different locations: Egypt (Port Saied, Bourg El-Arab, El Max/Alexandria Saline), Tibet (Lagkor Co), Turkmenistan (Kara-Bogaz-Gol) and Iran (Mighan Salt Lake, Lake Incheh, Lake Urmia and nearby laggoons). We did not study those populations. These samples could correspond to incorrect ploidy assignation or belong to the diploid lineage from which *Ap3n* originated. In addition, five individuals from China (Aqqikkol Lake) described as diploid (Asem et al. 2016) had a derived *Ap3n* haplotype (ANK1). Again, this suggests incorrect ploidy assignation or alternatively that the same mutation happened independently in both diploid and triploid individuals carrying the same APD05 haplotype. Finally, five individuals from Aibi Lake described as *Ap3n* (Asem et al. 2016) had the widespread APD01 *Ap2n-kaz*. This observation could again be due to an incorrect ploidy assignation or to the existence of a second clade of *Ap3n*. All these uncertainties have to be carefully checked. If the ploidy of these samples proved to be correctly assigned, we can conclude that *Ap3n* evolved multiple times independely from closely related *Ap2n-kaz* ancestors.

Maternal origin of *Ap4n* from an *Asin* population from East Siberia

We label this mitochondrial haplotype as being part of the mt-*sin* group, following current description (Naganawa and Mura 2017), but further investigations may be necessary. Indeed, those *Asin* from East Siberia might be morphologically close to *Asin* from China, but they show substantial divergence at the mitochondrial level (at least 11 mutation steps from the closest mt-*sin*, Fig. 5). *Asin* range is large, encompassing East Siberia, Mongolia and China (Litvinenko et al. 2009; Shadrin and Anufriieva 2012; Naganawa and Mura 2017) and we probably have only a limited representation of *Asin* diversity in the samples currently available. This will be resolved by additional sampling and genotyping.

*Ap5n* populations from Citros and Bujaraloz

We found the populations from Citros (Greece) and Bujaraloz (Spain) to be *Ap5n*, based on flow cytometry and nuclear data. We did not find any *Ap4n* individuals in these samples. In contrast, previous studies have found that these populations included only *Ap4n* individuals (Abatzopoulos et al. 1986; Amat et al. 1994; Maniatsi et al. 2011). These results could be due to incorrect ploidy assignation or to a mixture in these samples of *Ap4n* and *Ap5n* cysts with different survival rates.

*Species status of* Artemia frameshifta *and* A. murae

Individuals from Mongolia have recently been proposed to belong to two new species: *A. murae* and *A. frameshifta* (Naganawa and Mura 2017). This description was mainly based on the proximity of the COI sequences of these two species to *Asin* and *Aurm* respectively. We found that these two sequences are likely to be numts. The occurrence of pseudogenes when using universal COI markers is extremely frequent in *Artemia* spp., amounting to around 10% of COI sequences in our samples. Consequently, phylogenetic inferences based on these markers is unreliable and cannot be used to describe new species. The species *Artemia murae* appears closely related to *Asin* and further morpholocial and genetic evidence is required to make it a distinct species. Similarly, further morphological and genetic evidence is required to know whether *A. frameshifta* is different from other described *Artemia* species.

**Table S1. New genetic distance for *Ap2n* automicts.** Expected genetic distance between two individuals with genotypes corresponding the nine combinations of microsatellite genotypes potentially scored at one locus, as a function of the probability of recombination (r) and mutation ($\mu$). Q represents the fraction of null hererozygotes (A∅) among observed homozygotes (AA+ A∅)

| Scored genotypes | Expected genetic distance | Actual genotypes | Probability occurrence of actual genotypes | Genetic distance between genotypes |
| --- | --- | --- | --- | --- |
| AA / AA | $2Q(1-Q)/\mu$ | AA / AA | ${(1-Q)}^{2}$ | 0 |
|  |  | AA / A∅ | $Q(1-Q)$ | $1/\mu$ |
|  |  | A∅ / AA | $Q(1-Q)$ | $1/\mu$ |
|  |  | A∅ / A∅ | $Q^{2}$ | $0$ |
| AA / AB | $\left( 1-Q \right)/{(\mu+r)}+Q/\mu$ | AA / AB | $1-Q$ | $1/(\mu+r)$ |
|  |  | A∅ / AB | $Q$ | $1/\mu$ |
| AA / BB | ${\left( 1-Q \right)^{2}}/\varphi+ {2Q\left( 1-Q \right)}/\rho+{Q^{2}}/\mu$ | AA / BB | ${(1-Q)}^{2}$ | $1/\varphi$ (**) |
|  |  | A∅ / BB | $Q(1-Q)$ | $1/\rho$ (*) |
|  |  | AA / B∅ | $Q(1-Q)$ | $1/\rho$ (*) |
|  |  | A∅ / B∅ | $Q^{2}$ | $1/\mu$ |
| AB / AB | $0$ | AB / AB | $1$ | $0$ |
| AA / BC | $\left( 1-Q \right)/\rho+{2Q}/\mu$ | AA / BC | $1-Q$ | $1/\rho$ (*) |
|  |  | A∅ / BC | $Q$ | $2/\mu$ |
| AB / AC | $1/\mu$ | AB / AC | $1$ | $1/\mu$ |
| AB / CD | $2/\mu$ | AB / CD | $1$ | $2/\mu$ |
| AA / ∅∅ | $\left( 1-Q \right)/\varphi+Q/{(\mu+r)}$ | AA / ∅∅ | $1-Q$ | $1/\varphi$ (**) |
|  |  | A∅ / ∅∅ | $Q$ | $1/(\mu+r)$ |
| ∅∅ / ∅∅ | $0$ | ∅∅ / ∅∅ | $1$ | $0$ |

* $\rho={\mu\left( \mu+r \right)}/{(r+2\mu)}$ corresponds to the inverse of $1/\mu+1/(\mu+r)$, i.e. one mutation followed by one mutation or a LOH.

$**\varphi=r/2+\rho$ corresponds to the same events as above or, alternatively two LOH, whichever comes first.

**Table S2. Results of the AICc model selection for the different scenarios for the origin of *Ap2n-kaz* and *Ap2n-urm*.** See Fig. 3 and Mathematica notebook on Dryad for data and script details.

| **Model** | **Scenario** | **Aunk** | **Comment** | **Df** | **AIC** | **AICc** | **ΔAICc** |
| --- | --- | --- | --- | --- | --- | --- | --- |
| 18 | One hybridization, one backcross | Present | *Asin* closest to *Aunk* | 5 | -55.23 | -40.23 | 0.00 |
| 17 | One hybridization, one backcross | Present | *Asin* closest to *Akaz* | 5 | -52.72 | -37.72 | 2.51 |
| 16 | One hybridization, one backcross | Present | *Asin* closest to *Aurm* | 5 | -52.69 | -37.69 | 2.54 |
| 20 | One hybridization, one backcross | Present | *Asin* closest to *Akaz* | 5 | -41.43 | -26.43 | 13.80 |
| 21 | One hybridization, one backcross | Present | *Asin* closest to *Aunk* | 5 | -41.43 | -26.43 | 13.80 |
| 5 | One spontaneous origin, one hybridization | Absent | Secondary cross with *Akaz* | 5 | -40.00 | -25.00 | 15.23 |
| 14 | One hybridization, one backcross | Absent | Secondary backcross with *Akaz* | 3 | -22.77 | -18.77 | 21.46 |
| 11 | Two independent hybridizations | Present | *Asin* closest to *Aurm* | 5 | -33.32 | -18.32 | 21.91 |
| 12 | Two independent hybridizations | Present | *Asin* closest to *Akaz* | 5 | -32.99 | -17.99 | 22.24 |
| 13 | Two independent hybridizations | Present | *Asin* closest to *Aunk* | 5 | -32.99 | -17.99 | 22.24 |
| 19 | One hybridization, one backcross | Present | *Asin* closest to *Aurm* | 5 | -31.67 | -16.67 | 23.56 |
| 10 | Two independent hybridizations | Absent | - | 3 | -15.82 | -11.82 | 28.41 |
| 15 | One hybridization, one backcross | Absent | Secondary backcross with *Aurm* | 3 | -10.56 | -6.56 | 33.66 |
| 6 | One spontaneous origin, one hybridization | Absent | Secondary cross with *Aurm* | 5 | -16.07 | -1.07 | 39.16 |
| 7 | One spontaneous origin, one hybridization | Present | *Asin* closest to *Aurm* | 7 | -53.14 | 2.86 | 43.09 |
| 2 | Two independent spontaneous origins | Present | *Asin* closest to *Aurm* | 7 | -49.71 | 6.29 | 46.52 |
| 9 | One spontaneous origin, one hybridization | Present | *Asin* closest to *Aunk* | 7 | -46.46 | 9.54 | 49.77 |
| 8 | One spontaneous origin, one hybridization | Present | *Asin* closest to *Akaz* | 7 | -46.46 | 9.54 | 49.77 |
| 1 | Two independent spontaneous origins | Absent | - | 7 | -31.15 | 24.85 | 65.08 |
| 3 | Two independent spontaneous origins | Present | *Asin* closest to *Akaz* | 7 | -31.15 | 24.85 | 65.08 |
| 4 | Two independent spontaneous origins | Present | *Asin* closest to *Aunk* | 7 | -31.15 | 24.85 | 65.08 |

**Table S3. Comparison of the mean genetic distance between the 100 synthetic hybrids closest to observed genotypes of individuals from each ploidy.** Only the results of a Welsh’s t-test that compares the genetic distance obtained for the paternal species with the shortest genetic distance (i.e. *Aurm*, *Ap2n-kaz* and *Atib* for *Ap3n, Ap4n* and *Ap5n* respectively) and the genetic distance of other potential paternal species are reported.

|  | **Species** | **Mean genetic distance** | **SD genetic distance** | **df** | **t-test** | ***P-value*** |
| --- | --- | --- | --- | --- | --- | --- |
| *Ap3n* | *Akaz* | 0.253 | 0.015 | 184.09 | 22.81 | <2e-16 |
|  | *Ap2n-kaz* | 0.254 | 0.01 | 147.45 | -25.98 | <2e-16 |
|  | *Ap2n-urm* | 0.26 | 0.014 | 178.37 | -25.95 | <2e-16 |
|  | *Asin* | 0.324 | 0.012 | 165.26 | -54.8 | <2e-16 |
|  | *Atib* | 0.252 | 0.011 | 154.06 | -24.5 | <2e-16 |
|  | *Aurm* | 0.197 | 0.02 | - | - | NA |
| *Ap4n* | *Akaz* | 0.235 | 0.018 | 197.96 | -22.01 | <2e-16 |
|  | *Ap2n-kaz* | 0.179 | 0.018 | - | - | NA |
|  | *Ap2n-urm* | 0.185 | 0.017 | 196.96 | -2.21 | 0.0286 |
|  | *Asin* | 0.291 | 0.007 | 124.94 | -58.19 | <2e-16 |
|  | *Atib* | 0.204 | 0.014 | 186.17 | -11 | <2e-16 |
|  | *Aurm* | 0.234 | 0.032 | 156.04 | -14.81 | <2e-16 |
| *Ap5n* | *Akaz* | 0.173 | 0.006 | 154.95 | -4.37 | 2.27E-05 |
|  | *Ap2n-kaz* | 0.172 | 0.008 | 185.42 | -3.67 | 0.000314 |
|  | *Ap2n-urm* | 0.174 | 0.008 | 191.15 | -4.96 | 1.56E-06 |
|  | *Asin* | 0.177 | 0.007 | 170.6 | -7.99 | 1.90E-13 |
|  | *Atib* | 0.167 | 0.01 | - | - | - |
|  | *Aurm* | 0.204 | 0.006 | 164.06 | -30.75 | < 2e-16 |

**Table S4. List of the different *Artemia* populations studied**. Samples were characterized using flow cytometry, microsatellite or COI genotyping. The three methods were used on a reference set of 155 individuals (sample size: 72 *Ap2n*, 17 *Ap3n*, 16 *Ap4n*, 25 *Ap5n*, 9 *A*. sp Kaz, 3 *A. sinica*, 10 *A. tibetiana*, 1 *A. urmiana*). ARC: Artemia Reference Center, Ghent, Belgium. IATS col: Cyst collection from Instituto de Acuicultura de Torre de la Sal, CSIC, Castellon, Spain (https://www.gbif.es/en/coleccion/banco-de-quistes-de-artemia-instituto-de-acuicultura-de-torre-de-la-sal/). CEFE col: Cyst collection from Centre d’Écologie Fonctionnelle et Évolutive, Montpellier, France. *Afra*: *Artemia franciscana*, *Asal*: *Artemia salina*.

| Species | Population, Country | Population abbreviation | Sample size genotyping | Genome size estimation | Sample size for COI genotyping | Sample size used for cloning (total/mt DNA) | Reference number | Coordinates | Collection year | Other species |
| --- | --- | --- | --- | --- | --- | --- | --- | --- | --- | --- |
| *Aurm* | Lake Urmia, Iran* | URM | 26 | 5 | 12 | 0/1 | ARC1542 | 37.33, 45.67 | 2002 |  |
| *Akaz* | Unknown, Kazakhstan* | KAZ | 18 | 9 | 16 | 0/0 | ARC1039 | 51.16, 71.47 | 1987 |  |
| *Atib* | Lagkor Co, China | LAG | 23 | 10 | 23 | 0/1 | ARC1347 | 32.05, 84.22 | 1997 |  |
| *Asin* | Unknown, Yimeng area, China | YIM | 10 | 4 | 3 | 0/0 | ARC1188 | 39.17, 108.92 | 1991 |  |
|  | Dongjiagou, China | DON | 1 | 0 | 0 | 0/0 | ARC1216 | 39.12, 122.04 | 1991 | Ap2n/4n/5n |
|  | Xiechi Lake, China | XIE | 2 | 0 | 8 | 0/0 | ARC1218 | 34.99, 111.02 | 1991 |  |
| *Ap2n* | Aibi Lake, China | AIB | 7 | 1 | 5 | 0/0 | ARC1236 | 44.88, 82.9 | 1992 |  |
|  | Aigues-Mortes, France* | AIM | 75 | 12 | 70 | 0/1 | Pers. coll.** | 43.52, 4.18 | 2011 | Afra |
|  | Atanasovko Lake, Bulgaria | ATA1 | 5 | 2 | 7 | 0/0 | IATS col. | 42.57, 27.47 | 2005 | Ap5n |
|  | Unknown, Bameng area, China | BAM | 16 | 0 | 14 | 0/0 | ARC1317 | 40.77, 107.45 | 1995 |  |
|  | Bras del port, Spain | BDP | 0 | 0 | 2 | 0/0 | IATS col. | 38.19, -0.61 | 2004 | Asal |
|  | Bolshoye Yarovoe, Russia | BOL | 11 | 0 | 6 | 0/0 | ARC1706 | 52.85, 78.62 | 2007 | Afra |
|  | Co Qen, China | COQ | 11 | 0 | 1 | 0/0 | ARC1526, ARC1612 | 31.07, 85.4 | 2001 | Ap4n |
|  | Dongjiagou, China | DON | 21 | 8 | 21 | 0/0 | ARC1216 | 39.12, 122.04 | 1991 | Asin, Ap5n |
|  | Izmir, Turkey | IZM | 7 | 7 | 7 | 0/0 | ARC1512 | 38.65, 26.88 | 2001 | Ap3n/4n/5n |
|  | Koyashskoye, Ukraine | KOY | 15 | 5 | 0 | 0/0 | IATS col. | 45.04, 36.2 | 2007 | Aurm |
|  | Kulundinskoye, Russia | KUL | 7 | 8 | 7 | 0/0 | ARC 1700 | 53.01, 79.51 | 2007 |  |
|  | La Mata, Spain | LAM | 1 | 10 | 27 | 0/0 | IATS col. | 38.04, -0.7 | 1988 |  |
|  | Larache, Morocco | LAR | 4 | 0 | 4 | 0/0 | IATS col. | 35.2, -6.12 | 2005 |  |
|  | La Palmes, France | LPM | 1 | 1 | 1 | 0/0 | Pers. coll. | 42.98, 3.02 | 2002 |  |
|  | Maharlu Lake, Iran | MAH | 15 | 0 | 14 | 0/0 | ARC 1327 | 29.47, 52.77 | 1997 |  |
|  | Molentargius, Italy | MOL | 18 | 10 | 9 | 0/1 | IATS col. | 39.23, 9.21 | 2004 |  |
|  | Narte, Albania | NAR | 5 | 1 | 5 | 0/0 | IATS col. | 40.5, 19.45 | 2006 | Ap3n |
|  | Odiel, Spain* | ODI | 15 | 14 | 14 | 0/0 | Pers. coll.** | 37.26, -6.99 | 2011 |  |
|  | N. S. Rocío, Spain | ROC | 4 | 0 | 5 | 0/0 | IATS col. | 36.86, -6.34 | 2001 |  |
|  | Salin De Giraud, France | SAG | 13 | 8 | 14 | 0/0 | Pers. coll.** | 43.42, 4.63 | 2011 | Afra |
|  | Tenefé, Spain | TEN | 4 | 0 | 5 | 0/0 | IATS col. | 27.81, -15.42 | 2005 |  |
|  | Tanggu, China | TNG | 1 | 0 | 1 | 0/0 | IATS col. | 38.93, 117.62 | 1989 |  |
|  | Sète-Villeroy, France | VIL | 12 | 12 | 12 | 0/0 | Pers. coll.** | 43.38, 3.62 | 2012 | Afra |
|  | Yingkou, China | YIN | 4 | 0 | 4 | 0/0 | IATS col. | 40.55, 122.32 | 1989 |  |
| *Ap3n* | Ankiembe, Madagascar | ANK | 11 | 9 | 9 | 0/1 | IATS col. | -23.39, 43.7 | Unkown |  |
|  | Izmir, Turkey | IZM | 4 | 2 | 4 | 0/0 | ARC1512 | 38.65, 26.88 | 2001 | Ap2n/4n/5n |
|  | Narte, Albania | NAR | 11 | 15 | 6 | 0/0 | IATS col. | 40.5, 19.45 | 2006 | Ap2n |
| *Ap4n* | Arcos de la salinas, Spain | ARS | 4 | 0 | 4 | 0/0 | IATS col. | 39.99, -1.06 | Unkown |  |
|  | Bethioua, Algeria | BET | 4 | 0 | 3 | 0/0 | IATS col. | 35.71, -0.28 | 2009 |  |
|  | Co Qen, China*** | COQ | 3 | 0 | 1 | 0/0 | ARC1526, ARC1612 | 31.07, 85.4 | 2001 | Ap2n |
|  | Imón, Spain | IMO | 4 | 0 | 4 | 0/0 | IATS col. | 41.16, -2.73 | Unkown |  |
|  | Izmir, Turkey | IZM | 2 | 2 | 2 | 1/0 | ARC1512 | 38.65, 26.88 | 2001 | Ap2n/3n/5n |
|  | Lavalduc, France | LAV | 7 | 7 | 6 | 0/0 | CEFE col. | 43.47, 4.97 | 1978 | Afra |
|  | Teke Lake, Kazakhstan | TEK | 9 | 7 | 7 | 1/0 | ARC 1363 | 53.82, 72.97 | 1997 |  |
| *Ap5n* | Atanasovko Lake, Bulgaria | ATA1 | 13 | 12 | 5 | 0/0 | IATS col. | 42.57, 27.47 | 2005 | Ap2n |
|  | Atanasovko Lake, Bulgaria | ATA2 | 6 | 4 | 4 | 0/0 | ARC 1220 | 42.57, 27.47 | 1991 |  |
|  | Bujaraloz, Spain* | BUJ | 14 | 10 | 14 | 1/0 | IATS col. | 41.42, -0.19 | Unkown |  |
|  | Citros, Greece* | CIT | 10 | 6 | 6 | 1/1 | IATS col. | 40.35, 22.64 | Unkown |  |
|  | Dongjiagou, China*** | DON | 2 | 0 | 2 | 1/0 | ARC1216 | 39.12, 122.04 | 1991 | Asin, Ap2n |
|  | Izmir, Turkey | IZM | 5 | 5 | 5 | 1/0 | ARC1512 | 38.65, 26.88 | 2001 | Ap2n/3n/4n |

*Genome size and microsatellite genotype data from Nougué *et al.* 2015b

** Wild-collected individuals

*** The genome size of these samples could not be measured; their ploidy was assigned based on their genetic proximity to *Ap4n* and *Ap5n* samples respectively.

**Table S5. List of the 1196 sequences collapsed into the 123 COI haplotypes whose network is represented in Fig. 5** (after removal of putative COI-numts and chimeras, except that from *Asin* from East Siberia). 399 sequences (hereafter “CEFE sequences”) were obtained for this study either directly (dataset1 and dataset2) or following a cloning step (dataset3). 797 additional sequences were obtained from the GenBank database (“GenBank” sequences) whose species and ploidy correspond to their original assignment. We retained respectively 237 CEFE and 748 GenBank putative COI sequences for further analyses (i.e. sequences with any ambiguous site and including less that 538 sites within the final alignment were removed). After excluding the 32 known numts, we found among the remaining 953 putative COI sequences 3 CEFE and 100 GenBank sequences that were potential numts or chimeras (assigned based on polarity changes in the amino acid sequence or on an unusually high number of mutations shared with another unrelated sequence of the dataset). Only sequences of high quality (dataset1 and dataset3) were deposited into GenBank.

| **AccNum** | **Species** | **Ploidy** | **Population** | **Name** | **Origin** | **Cloning** | **Proportion of ambiguous nucleotides** | **Length of the sequence** | **Dataset** | **Polarity Change** | **Potential Numt?** | **Chimera Score** | **Numt or Chimera?** | **Haplogroup** |
| --- | --- | --- | --- | --- | --- | --- | --- | --- | --- | --- | --- | --- | --- | --- |
| NA | Akaz | 2n | KAZ | AkazKAZ9 | This study | No | 0.01 | 613 | dataset2 | NA | NA | NA | 0 | Akaz |
| NA | Akaz | 2n | KAZ | AkazKAZ1 | This study | No | 0.01 | 615 | dataset2 | NA | NA | NA | 0 | Akaz |
| NA | Akaz | 2n | KAZ | AkazKAZ2 | This study | No | 0.01 | 617 | dataset2 | NA | NA | NA | 0 | Akaz |
| NA | Akaz | 2n | KAZ | AkazKAZ3 | This study | No | 0.00 | 625 | dataset2 | NA | NA | NA | 0 | Akaz |
| NA | Akaz | 2n | KAZ | AkazKAZ4 | This study | No | 0.00 | 625 | dataset2 | NA | NA | NA | 0 | Akaz |
| NA | Akaz | 2n | KAZ | AkazKAZ5 | This study | No | 0.02 | 618 | dataset2 | NA | NA | NA | 0 | Akaz |
| NA | Akaz | 2n | KAZ | AkazKAZ10 | This study | No | 0.02 | 590 | dataset2 | NA | NA | NA | 0 | Akaz |
| NA | Akaz | 2n | KAZ | AkazKAZ11 | This study | No | 0.05 | 466 | dataset2 | NA | NA | NA | 0 | Akaz |
| MT791641 | Akaz | 2n | KAZ | AkazKAZ12 | This study | No | 0.00 | 627 | dataset1 | 0 | 0 | 1 | 0 | Akaz |
| MT791642 | Akaz | 2n | KAZ | AkazKAZ13 | This study | No | 0.00 | 627 | dataset1 | 0 | 0 | 1 | 0 | Akaz |
| NA | Akaz | 2n | KAZ | AkazKAZ14 | This study | No | 0.01 | 627 | dataset2 | NA | NA | NA | 0 | Akaz |
| NA | Akaz | 2n | KAZ | AkazKAZ15 | This study | No | 0.02 | 589 | dataset2 | NA | NA | NA | 0 | Akaz |
| MT791643 | Akaz | 2n | KAZ | AkazKAZ16 | This study | No | 0.00 | 627 | dataset1 | 0 | 0 | NA | 0 | Akaz |
| NA | Akaz | 2n | KAZ | AkazKAZ7 | This study | No | 0.00 | 627 | dataset2 | NA | NA | NA | 0 | Akaz |
| NA | Akaz | 2n | KAZ | AkazKAZ8 | This study | No | 0.02 | 495 | dataset2 | NA | NA | NA | 0 | Akaz |
| MT791644 | Akaz | 2n | KAZ | AkazKAZ6 | This study | No | 0.00 | 627 | dataset1 | 0 | 0 | 1 | 0 | Akaz |
| DQ119653 | Akaz | 2n | NA | AspKazHou | GenBank | No | 0.00 | 658 | dataset6 | NA | NA | NA | 1 | Akaz |
| GU591385 | Akaz | 2n | NA | KAZSEX01 | GenBank | No | 0.00 | 658 | dataset5 | 0 | 0 | 1 | 0 | Akaz |
| GU591386 | Akaz | 2n | NA | KAZSEX02 | GenBank | No | 0.00 | 658 | dataset5 | 0 | 0 | 1 | 0 | Akaz |
| GU591387 | Akaz | 2n | NA | KAZSEX03 | GenBank | No | 0.00 | 658 | dataset5 | 0 | 0 | NA | 0 | Akaz |
| GU591388 | Akaz | 2n | NA | KAZSEX04 | GenBank | No | 0.00 | 658 | dataset5 | 0 | 0 | NA | 0 | Akaz |
| GU591389 | Akaz | 2n | NA | KAZSEX05 | GenBank | No | 0.00 | 658 | dataset5 | 0 | 0 | NA | 0 | Akaz |
| KF707671 | Akaz | 2n | NA | KAZ3 | GenBank | No | 0.00 | 614 | dataset5 | 0 | 0 | NA | 0 | Akaz |
| KF707672 | Akaz | 2n | NA | KAZ7 | GenBank | No | 0.00 | 614 | dataset5 | 0 | 0 | NA | 0 | Akaz |
| KF707673 | Akaz | 2n | NA | KAZ10 | GenBank | No | 0.00 | 614 | dataset5 | 0 | 0 | NA | 0 | Akaz |
| KF707674 | Akaz | 2n | NA | KAZ9 | GenBank | No | 0.00 | 614 | dataset5 | 0 | 0 | NA | 0 | Akaz |
| KF707675 | Akaz | 2n | NA | KAZ8 | GenBank | No | 0.00 | 614 | dataset5 | 0 | 0 | 1 | 0 | Akaz |
| KF707676 | Akaz | 2n | NA | KAZ6 | GenBank | No | 0.00 | 614 | dataset5 | 0 | 0 | NA | 0 | Akaz |
| KF707677 | Akaz | 2n | NA | KAZ5 | GenBank | No | 0.00 | 614 | dataset5 | 0 | 0 | NA | 0 | Akaz |
| KF707678 | Akaz | 2n | NA | KAZ4 | GenBank | No | 0.00 | 614 | dataset5 | 0 | 0 | NA | 0 | Akaz |
| KF707679 | Akaz | 2n | NA | KAZ2 | GenBank | No | 0.00 | 614 | dataset5 | 0 | 0 | NA | 0 | Akaz |
| KF707680 | Akaz | 2n | NA | KAZ1 | GenBank | No | 0.00 | 614 | dataset5 | 0 | 0 | NA | 0 | Akaz |
| MT791694 | Ap2n | 2n | BAM | Ap2nBAM1 | This study | No | 0.00 | 629 | dataset1 | 0 | 0 | NA | 0 | Ap2n-kaz |
| MT791695 | Ap2n | 2n | BAM | Ap2nBAM2 | This study | No | 0.00 | 633 | dataset1 | 0 | 0 | 1 | 0 | Ap2n-kaz |
| MT791696 | Ap2n | 2n | BAM | Ap2nBAM3 | This study | No | 0.00 | 647 | dataset1 | 0 | 0 | NA | 0 | Ap2n-kaz |
| MT791697 | Ap2n | 2n | BAM | Ap2nBAM4 | This study | No | 0.00 | 629 | dataset1 | 0 | 0 | NA | 0 | Ap2n-kaz |
| MT791698 | Ap2n | 2n | BAM | Ap2nBAM5 | This study | No | 0.00 | 640 | dataset1 | 0 | 0 | NA | 0 | Ap2n-kaz |
| MT791699 | Ap2n | 2n | BAM | Ap2nBAM6 | This study | No | 0.00 | 630 | dataset1 | 0 | 0 | NA | 0 | Ap2n-kaz |
| MT791700 | Ap2n | 2n | BAM | Ap2nBAM7 | This study | No | 0.00 | 640 | dataset1 | 0 | 0 | NA | 0 | Ap2n-kaz |
| MT791701 | Ap2n | 2n | BAM | Ap2nBAM10 | This study | No | 0.00 | 640 | dataset1 | 0 | 0 | NA | 0 | Ap2n-kaz |
| MT791702 | Ap2n | 2n | BAM | Ap2nBAM11 | This study | No | 0.00 | 640 | dataset1 | 0 | 0 | NA | 0 | Ap2n-kaz |
| MT791703 | Ap2n | 2n | BAM | Ap2nBAM13 | This study | No | 0.00 | 620 | dataset1 | 0 | 0 | NA | 0 | Ap2n-kaz |
| NA | Ap2n | 2n | AIM | Ap2nAIM10 | This study | No | 0.00 | 635 | dataset1 | 2.33 | 1 | 3 | 1 | Ap2n |
| MT791664 | Ap2n | 2n | AIM | Ap2nAIM11 | This study | No | 0.00 | 700 | dataset1 | 0.06 | 0 | 1 | 0 | Ap2n-urm |
| MT791665 | Ap2n | 2n | AIM | Ap2nAIM12 | This study | No | 0.00 | 700 | dataset1 | 0 | 0 | NA | 0 | Ap2n-urm |
| MT791666 | Ap2n | 2n | AIM | Ap2nAIM13 | This study | No | 0.00 | 699 | dataset1 | 0 | 0 | NA | 0 | Ap2n-urm |
| MT791667 | Ap2n | 2n | AIM | Ap2nAIM14 | This study | No | 0.00 | 688 | dataset1 | 0 | 0 | NA | 0 | Ap2n-urm |
| NA | Ap2n | 2n | AIM | Ap2nAIM15 | This study | No | 0.01 | 713 | dataset2 | NA | NA | NA | 0 | Ap2n-urm |
| NA | Ap2n | 2n | AIM | Ap2nAIM18 | This study | No | 0.02 | 713 | dataset2 | NA | NA | NA | 1 | Ap2n |
| MT791647 | Ap2n | 2n | AIM | Ap2nAIM19 | This study | No | 0.00 | 686 | dataset1 | 0 | 0 | NA | 0 | Ap2n-kaz |
| MT791704 | Ap2n | 2n | BAM | Ap2nBAM14 | This study | No | 0.00 | 688 | dataset1 | 0 | 0 | NA | 0 | Ap2n-kaz |
| MT791754 | Ap2n | 2n | LPM | Ap2nLPM1 | This study | No | 0.00 | 687 | dataset1 | 0.06 | 0 | NA | 0 | Ap2n-urm |
| MT791729 | Ap2n | 2n | LAM | Ap2nLAM4 | This study | No | 0.00 | 686 | dataset1 | 0.06 | 0 | NA | 0 | Ap2n-urm |
| MT791730 | Ap2n | 2n | LAM | Ap2nLAM5 | This study | No | 0.00 | 685 | dataset1 | 0.06 | 0 | NA | 0 | Ap2n-urm |
| MT791731 | Ap2n | 2n | LAM | Ap2nLAM6 | This study | No | 0.00 | 685 | dataset1 | 0.06 | 0 | NA | 0 | Ap2n-urm |
| MT791732 | Ap2n | 2n | LAM | Ap2nLAM7 | This study | No | 0.00 | 686 | dataset1 | 0.06 | 0 | NA | 0 | Ap2n-urm |
| MT791733 | Ap2n | 2n | LAM | Ap2nLAM8 | This study | No | 0.00 | 686 | dataset1 | 0.06 | 0 | NA | 0 | Ap2n-urm |
| MT791734 | Ap2n | 2n | LAM | Ap2nLAM9 | This study | No | 0.00 | 686 | dataset1 | 0.06 | 0 | NA | 0 | Ap2n-urm |
| MT791735 | Ap2n | 2n | LAM | Ap2nLAM10 | This study | No | 0.00 | 688 | dataset1 | 0.06 | 0 | NA | 0 | Ap2n-urm |
| MT791769 | Ap2n | 2n | MOL | Ap2nMOL1 | This study | No | 0.00 | 616 | dataset1 | 0 | 0 | NA | 0 | Ap2n-urm |
| NA | Ap2n | 2n | MOL | Ap2nMOL4 | This study | No | 0.00 | 568 | dataset1 | 0 | 0 | 1 | 1 | Ap2n |
| MT791770 | Ap2n | 2n | MOL | Ap2nMOL5 | This study | No | 0.00 | 688 | dataset1 | 0 | 0 | NA | 0 | Ap2n-urm |
| MT791771 | Ap2n | 2n | MOL | Ap2nMOL6 | This study | No | 0.00 | 654 | dataset1 | 0 | 0 | NA | 0 | Ap2n-urm |
| NA | Ap2n | 2n | MOL | Ap2nMOL7 | This study | No | 0.00 | 688 | dataset2 | NA | NA | NA | 0 | Ap2n-urm |
| NA | Ap2n | 2n | AIM | Ap2nAIM3 | This study | No | 0.00 | 687 | dataset2 | NA | NA | NA | 1 | Ap2n |
| MT791736 | Ap2n | 2n | LAM | Ap2nLAM11 | This study | No | 0.00 | 639 | dataset1 | 0.06 | 0 | NA | 0 | Ap2n-urm |
| MT791737 | Ap2n | 2n | LAM | Ap2nLAM12 | This study | No | 0.00 | 651 | dataset1 | 0.06 | 0 | 0 | 0 | Ap2n-urm |
| MT791738 | Ap2n | 2n | LAM | Ap2nLAM13 | This study | No | 0.00 | 684 | dataset1 | 0.06 | 0 | NA | 0 | Ap2n-urm |
| MT791648 | Ap2n | 2n | AIM | Ap2nAIM4 | This study | No | 0.00 | 580 | dataset1 | 0 | 0 | NA | 0 | Ap2n-kaz |
| NA | Ap2n | 2n | MOL | Ap2nMOL3 | This study | No | 0.00 | 617 | dataset2 | NA | NA | NA | 0 | Ap2n-urm |
| MT791668 | Ap2n | 2n | AIM | Ap2nAIM21 | This study | No | 0.00 | 629 | dataset1 | 0 | 0 | NA | 0 | Ap2n-urm |
| MT791669 | Ap2n | 2n | AIM | Ap2nAIM22 | This study | No | 0.00 | 615 | dataset1 | 0 | 0 | NA | 0 | Ap2n-urm |
| MT791670 | Ap2n | 2n | AIM | Ap2nAIM23 | This study | No | 0.00 | 596 | dataset1 | 0 | 0 | NA | 0 | Ap2n-urm |
| NA | Ap2n | 2n | AIM | Ap2nAIM24 | This study | No | 0.01 | 468 | dataset2 | NA | NA | NA | 0 | Ap2n-urm |
| MT791649 | Ap2n | 2n | AIM | Ap2nAIM25 | This study | No | 0.00 | 657 | dataset1 | 0 | 0 | NA | 0 | Ap2n-kaz |
| MT791671 | Ap2n | 2n | AIM | Ap2nAIM26 | This study | No | 0.00 | 690 | dataset1 | 0 | 0 | NA | 0 | Ap2n-urm |
| MT791672 | Ap2n | 2n | AIM | Ap2nAIM29 | This study | No | 0.00 | 687 | dataset1 | 0.06 | 0 | NA | 0 | Ap2n-urm |
| MT791673 | Ap2n | 2n | AIM | Ap2nAIM30 | This study | No | 0.00 | 588 | dataset1 | 0 | 0 | NA | 0 | Ap2n-urm |
| MT791674 | Ap2n | 2n | AIM | Ap2nAIM31 | This study | No | 0.00 | 562 | dataset1 | 0 | 0 | NA | 0 | Ap2n-urm |
| MT791739 | Ap2n | 2n | LAM | Ap2nLAM14 | This study | No | 0.00 | 688 | dataset1 | 0.06 | 0 | NA | 0 | Ap2n-urm |
| MT791675 | Ap2n | 2n | AIM | Ap2nAIM32 | This study | No | 0.00 | 657 | dataset1 | 0 | 0 | NA | 0 | Ap2n-urm |
| MT791740 | Ap2n | 2n | LAM | Ap2nLAM15 | This study | No | 0.00 | 686 | dataset1 | 0.06 | 0 | NA | 0 | Ap2n-urm |
| MT791741 | Ap2n | 2n | LAM | Ap2nLAM16 | This study | No | 0.00 | 686 | dataset1 | 0.06 | 0 | NA | 0 | Ap2n-urm |
| NA | Ap2n | 2n | AIM | Ap2nAIM33 | This study | No | 0.01 | 511 | dataset2 | NA | NA | NA | 1 | Ap2n |
| MT791742 | Ap2n | 2n | LAM | Ap2nLAM17 | This study | No | 0.00 | 688 | dataset1 | 0.06 | 0 | NA | 0 | Ap2n-urm |
| MT791743 | Ap2n | 2n | LAM | Ap2nLAM18 | This study | No | 0.00 | 635 | dataset1 | 0.06 | 0 | NA | 0 | Ap2n-urm |
| MT791676 | Ap2n | 2n | AIM | Ap2nAIM34 | This study | No | 0.00 | 657 | dataset1 | 0 | 0 | NA | 0 | Ap2n-urm |
| MT791744 | Ap2n | 2n | LAM | Ap2nLAM19 | This study | No | 0.00 | 655 | dataset1 | 0.06 | 0 | NA | 0 | Ap2n-urm |
| MT791677 | Ap2n | 2n | AIM | Ap2nAIM35 | This study | No | 0.00 | 688 | dataset1 | 0 | 0 | NA | 0 | Ap2n-urm |
| MT791745 | Ap2n | 2n | LAM | Ap2nLAM20 | This study | No | 0.00 | 655 | dataset1 | 0.06 | 0 | NA | 0 | Ap2n-urm |
| MT791678 | Ap2n | 2n | AIM | Ap2nAIM37 | This study | No | 0.00 | 654 | dataset1 | 0 | 0 | 1 | 0 | Ap2n-urm |
| MT791746 | Ap2n | 2n | LAM | Ap2nLAM21 | This study | No | 0.00 | 651 | dataset1 | 0.06 | 0 | NA | 0 | Ap2n-urm |
| MT791747 | Ap2n | 2n | LAM | Ap2nLAM22 | This study | No | 0.00 | 596 | dataset1 | 0.06 | 0 | NA | 0 | Ap2n-urm |
| MT791679 | Ap2n | 2n | AIM | Ap2nAIM39 | This study | No | 0.00 | 688 | dataset1 | 0 | 0 | NA | 0 | Ap2n-urm |
| MT791748 | Ap2n | 2n | LAM | Ap2nLAM23 | This study | No | 0.00 | 688 | dataset1 | 0.06 | 0 | NA | 0 | Ap2n-urm |
| MT791680 | Ap2n | 2n | AIM | Ap2nAIM40 | This study | No | 0.00 | 655 | dataset1 | 0 | 0 | NA | 0 | Ap2n-urm |
| MT791681 | Ap2n | 2n | AIM | Ap2nAIM41 | This study | No | 0.00 | 543 | dataset1 | 0 | 0 | NA | 0 | Ap2n-urm |
| MT791749 | Ap2n | 2n | LAM | Ap2nLAM24 | This study | No | 0.00 | 688 | dataset1 | 0.06 | 0 | NA | 0 | Ap2n-urm |
| NA | Ap2n | 2n | AIM | Ap2nAIM42 | This study | No | 0.01 | 522 | dataset2 | NA | NA | NA | 0 | Ap2n-kaz |
| MT791750 | Ap2n | 2n | LAM | Ap2nLAM25 | This study | No | 0.00 | 647 | dataset1 | 0.06 | 0 | NA | 0 | Ap2n-urm |
| MT791682 | Ap2n | 2n | AIM | Ap2nAIM43 | This study | No | 0.00 | 688 | dataset1 | 0 | 0 | NA | 0 | Ap2n-urm |
| MT791751 | Ap2n | 2n | LAM | Ap2nLAM26 | This study | No | 0.00 | 623 | dataset1 | 0.06 | 0 | NA | 0 | Ap2n-urm |
| MT791752 | Ap2n | 2n | LAM | Ap2nLAM27 | This study | No | 0.00 | 591 | dataset1 | 0.06 | 0 | NA | 0 | Ap2n-urm |
| MT791683 | Ap2n | 2n | AIM | Ap2nAIM44 | This study | No | 0.00 | 654 | dataset1 | 0 | 0 | 1 | 0 | Ap2n-urm |
| MT791684 | Ap2n | 2n | AIM | Ap2nAIM45 | This study | No | 0.00 | 685 | dataset1 | 0 | 0 | NA | 0 | Ap2n-urm |
| MT791685 | Ap2n | 2n | AIM | Ap2nAIM46 | This study | No | 0.00 | 654 | dataset1 | 0 | 0 | NA | 0 | Ap2n-urm |
| MT791686 | Ap2n | 2n | AIM | Ap2nAIM47 | This study | No | 0.00 | 688 | dataset1 | 0 | 0 | NA | 0 | Ap2n-urm |
| MT791687 | Ap2n | 2n | AIM | Ap2nAIM48 | This study | No | 0.00 | 688 | dataset1 | 0 | 0 | NA | 0 | Ap2n-urm |
| MT791688 | Ap2n | 2n | AIM | Ap2nAIM49 | This study | No | 0.00 | 559 | dataset1 | 0 | 0 | NA | 0 | Ap2n-urm |
| MT791689 | Ap2n | 2n | AIM | Ap2nAIM50 | This study | No | 0.00 | 655 | dataset1 | 0 | 0 | NA | 0 | Ap2n-urm |
| MT791690 | Ap2n | 2n | AIM | Ap2nAIM51 | This study | No | 0.00 | 686 | dataset1 | 0 | 0 | NA | 0 | Ap2n-urm |
| MT791691 | Ap2n | 2n | AIM | Ap2nAIM54 | This study | No | 0.00 | 686 | dataset1 | 0 | 0 | NA | 0 | Ap2n-urm |
| MT791692 | Ap2n | 2n | AIM | Ap2nAIM55 | This study | No | 0.00 | 688 | dataset1 | 0 | 0 | NA | 0 | Ap2n-urm |
| MT791693 | Ap2n | 2n | AIM | Ap2nAIM56 | This study | No | 0.00 | 671 | dataset1 | 0 | 0 | NA | 0 | Ap2n-urm |
| NA | Ap2n | 2n | AIM | Ap2nAIM1 | This study | No | 0.00 | 521 | dataset2 | NA | NA | NA | 0 | Ap2n-urm |
| NA | Ap2n | 2n | ATA1 | Ap2nATA11 | This study | No | 0.00 | 521 | dataset2 | NA | NA | NA | 0 | Ap2n-kaz |
| NA | Ap2n | 2n | BDP | Ap2nBDP1 | This study | No | 0.00 | 521 | dataset2 | NA | NA | NA | 0 | Ap2n-urm |
| NA | Ap2n | 2n | AIM | Ap2nAIM2 | This study | No | 0.00 | 521 | dataset2 | NA | NA | NA | 0 | Ap2n-urm |
| NA | Ap2n | 2n | AIM | Ap2nAIM5 | This study | No | 0.00 | 521 | dataset2 | NA | NA | NA | 0 | Ap2n-urm |
| NA | Ap2n | 2n | ATA1 | Ap2nATA13 | This study | No | 0.00 | 521 | dataset2 | NA | NA | NA | 0 | Ap2n-urm |
| NA | Ap2n | 2n | ATA1 | Ap2nATA14 | This study | No | 0.00 | 521 | dataset2 | NA | NA | NA | 0 | Ap2n-kaz |
| NA | Ap2n | 2n | ATA1 | Ap2nATA15 | This study | No | 0.00 | 521 | dataset2 | NA | NA | NA | 0 | Ap2n-kaz |
| NA | Ap2n | 2n | ATA1 | Ap2nATA16 | This study | No | 0.00 | 521 | dataset2 | NA | NA | NA | 0 | Ap2n-kaz |
| NA | Ap2n | 2n | LAR | Ap2nLAR1 | This study | No | 0.00 | 521 | dataset2 | NA | NA | NA | 0 | Ap2n-kaz |
| NA | Ap2n | 2n | LAR | Ap2nLAR3 | This study | No | 0.00 | 522 | dataset2 | NA | NA | NA | 0 | Ap2n-kaz |
| NA | Ap2n | 2n | NAR | Ap2nNAR1 | This study | No | 0.00 | 521 | dataset2 | NA | NA | NA | 0 | Ap2n-kaz |
| NA | Ap2n | 2n | NAR | Ap2nNAR4 | This study | No | 0.00 | 521 | dataset2 | NA | NA | NA | 0 | Ap2n-urm |
| NA | Ap2n | 2n | ROC | Ap2nROC1 | This study | No | 0.00 | 521 | dataset2 | NA | NA | NA | 0 | Ap2n-kaz |
| NA | Ap2n | 2n | ROC | Ap2nROC5 | This study | No | 0.00 | 520 | dataset2 | NA | NA | NA | 0 | Ap2n-kaz |
| NA | Ap2n | 2n | TNG | Ap2nTNG1 | This study | No | 0.00 | 522 | dataset2 | NA | NA | NA | 0 | Ap2n-kaz |
| NA | Ap2n | 2n | TEN | Ap2nTEN1 | This study | No | 0.00 | 522 | dataset2 | NA | NA | NA | 0 | Ap2n-kaz |
| NA | Ap2n | 2n | TEN | Ap2nTEN4 | This study | No | 0.00 | 521 | dataset2 | NA | NA | NA | 0 | Ap2n-kaz |
| NA | Ap2n | 2n | TEN | Ap2nTEN5 | This study | No | 0.00 | 522 | dataset2 | NA | NA | NA | 0 | Ap2n-kaz |
| NA | Ap2n | 2n | YIN | Ap2nYIN2 | This study | No | 0.00 | 511 | dataset2 | NA | NA | NA | 0 | Ap2n-kaz |
| NA | Ap2n | 2n | YIN | Ap2nYIN3 | This study | No | 0.00 | 521 | dataset2 | NA | NA | NA | 0 | Ap2n-kaz |
| NA | Ap2n | 2n | AIB | Ap2nAIB4 | This study | No | 0.00 | 627 | dataset2 | NA | NA | NA | 0 | Ap2n-urm |
| NA | Ap2n | 2n | AIB | Ap2nAIB5 | This study | No | 0.01 | 627 | dataset2 | NA | NA | NA | 1 | Ap2n |
| NA | Ap2n | 2n | IZM | Ap2nIZM1 | This study | No | 0.00 | 627 | dataset2 | NA | NA | NA | 0 | Ap2n-urm |
| NA | Ap2n | 2n | IZM | Ap2nIZM3 | This study | No | 0.00 | 627 | dataset2 | NA | NA | NA | 0 | Ap2n-urm |
| NA | Ap2n | 2n | IZM | Ap2nIZM4 | This study | No | 0.00 | 627 | dataset2 | NA | NA | NA | 0 | Ap2n-urm |
| NA | Ap2n | 2n | IZM | Ap2nIZM5 | This study | No | 0.00 | 623 | dataset2 | NA | NA | NA | 0 | Ap2n-urm |
| NA | Ap2n | 2n | IZM | Ap2nIZM6 | This study | No | 0.00 | 623 | dataset2 | NA | NA | NA | 0 | Ap2n-urm |
| NA | Ap2n | 2n | IZM | Ap2nIZM2 | This study | No | 0.00 | 625 | dataset2 | NA | NA | NA | 0 | Ap2n-urm |
| NA | Ap2n | 2n | IZM | Ap2nIZM7 | This study | No | 0.00 | 627 | dataset2 | NA | NA | NA | 0 | Ap2n-kaz |
| NA | Ap2n | 2n | KUL | Ap2nKUL1 | This study | No | 0.01 | 625 | dataset2 | NA | NA | NA | 0 | Ap2n-kaz |
| MT791645 | Ap2n | 2n | AIB | Ap2nAIB1 | This study | No | 0.00 | 685 | dataset1 | 0 | 0 | 1 | 0 | Ap2n-urm |
| MT791646 | Ap2n | 2n | AIB | Ap2nAIB2 | This study | No | 0.00 | 685 | dataset1 | 0 | 0 | NA | 0 | Ap2n-urm |
| MT791710 | Ap2n | 2n | DON | Ap2nDON1 | This study | No | 0.00 | 687 | dataset1 | 0 | 0 | NA | 0 | Ap2n-kaz |
| MT791711 | Ap2n | 2n | DON | Ap2nDON2 | This study | No | 0.00 | 688 | dataset1 | 0 | 0 | NA | 0 | Ap2n-kaz |
| MT791712 | Ap2n | 2n | DON | Ap2nDON3 | This study | No | 0.00 | 688 | dataset1 | 0 | 0 | NA | 0 | Ap2n-kaz |
| MT791713 | Ap2n | 2n | DON | Ap2nDON4 | This study | No | 0.00 | 687 | dataset1 | 0 | 0 | NA | 0 | Ap2n-kaz |
| MT791714 | Ap2n | 2n | DON | Ap2nDON5 | This study | No | 0.00 | 688 | dataset1 | 0 | 0 | NA | 0 | Ap2n-kaz |
| MT791715 | Ap2n | 2n | DON | Ap2nDON6 | This study | No | 0.00 | 687 | dataset1 | 0 | 0 | NA | 0 | Ap2n-kaz |
| MT791716 | Ap2n | 2n | DON | Ap2nDON8 | This study | No | 0.00 | 685 | dataset1 | 0 | 0 | NA | 0 | Ap2n-kaz |
| NA | Ap2n | 2n | DON | Ap2nDON9 | This study | No | 0.00 | 685 | dataset2 | NA | NA | NA | 0 | Ap2n-kaz |
| MT791717 | Ap2n | 2n | DON | Ap2nDON10 | This study | No | 0.00 | 687 | dataset1 | 0 | 0 | NA | 0 | Ap2n-kaz |
| MT791718 | Ap2n | 2n | DON | Ap2nDON11 | This study | No | 0.00 | 688 | dataset1 | 0 | 0 | NA | 0 | Ap2n-kaz |
| NA | Ap2n | 2n | DON | Ap2nDON12 | This study | No | 0.00 | 687 | dataset2 | NA | NA | NA | 1 | Ap2n |
| MT791719 | Ap2n | 2n | DON | Ap2nDON14 | This study | No | 0.00 | 687 | dataset1 | 0 | 0 | NA | 0 | Ap2n-kaz |
| MT791720 | Ap2n | 2n | DON | Ap2nDON15 | This study | No | 0.00 | 689 | dataset1 | 0 | 0 | NA | 0 | Ap2n-kaz |
| MT791784 | Ap2n | 2n | SAG | Ap2nSAG2 | This study | No | 0.00 | 687 | dataset1 | 0 | 0 | NA | 0 | Ap2n-urm |
| MT791783 | Ap2n | 2n | SAG | Ap2nSAG3 | This study | No | 0.00 | 686 | dataset1 | 0 | 0 | NA | 0 | Ap2n-kaz |
| NA | Ap2n | 2n | SAG | Ap2nSAG4 | This study | No | 0.00 | 686 | dataset2 | NA | NA | NA | 0 | Ap2n-urm |
| NA | Ap2n | 2n | LAR | Ap2nLAR2 | This study | No | 0.00 | 684 | dataset2 | NA | NA | NA | 1 | Ap2n |
| MT791753 | Ap2n | 2n | LAR | Ap2nLAR4 | This study | No | 0.00 | 686 | dataset1 | 0 | 0 | NA | 0 | Ap2n-kaz |
| MT791755 | Ap2n | 2n | MAH | Ap2nMAH1 | This study | No | 0.00 | 684 | dataset1 | 0 | 0 | NA | 0 | Ap2n-kaz |
| MT791756 | Ap2n | 2n | MAH | Ap2nMAH2 | This study | No | 0.00 | 686 | dataset1 | 0 | 0 | NA | 0 | Ap2n-kaz |
| MT791757 | Ap2n | 2n | MAH | Ap2nMAH3 | This study | No | 0.00 | 686 | dataset1 | 0 | 0 | NA | 0 | Ap2n-kaz |
| MT791758 | Ap2n | 2n | MAH | Ap2nMAH4 | This study | No | 0.00 | 685 | dataset1 | 0 | 0 | NA | 0 | Ap2n-kaz |
| MT791759 | Ap2n | 2n | MAH | Ap2nMAH5 | This study | No | 0.00 | 685 | dataset1 | 0 | 0 | NA | 0 | Ap2n-kaz |
| MT791760 | Ap2n | 2n | MAH | Ap2nMAH6 | This study | No | 0.00 | 685 | dataset1 | 0 | 0 | NA | 0 | Ap2n-kaz |
| MT791761 | Ap2n | 2n | MAH | Ap2nMAH8 | This study | No | 0.00 | 685 | dataset1 | 0 | 0 | NA | 0 | Ap2n-kaz |
| MT791762 | Ap2n | 2n | MAH | Ap2nMAH9 | This study | No | 0.00 | 686 | dataset1 | 0 | 0 | NA | 0 | Ap2n-kaz |
| MT791763 | Ap2n | 2n | MAH | Ap2nMAH10 | This study | No | 0.00 | 687 | dataset1 | 0 | 0 | NA | 0 | Ap2n-kaz |
| MT791764 | Ap2n | 2n | MAH | Ap2nMAH11 | This study | No | 0.00 | 684 | dataset1 | 0 | 0 | NA | 0 | Ap2n-kaz |
| MT791765 | Ap2n | 2n | MAH | Ap2nMAH12 | This study | No | 0.00 | 684 | dataset1 | 0 | 0 | NA | 0 | Ap2n-kaz |
| MT791766 | Ap2n | 2n | MAH | Ap2nMAH13 | This study | No | 0.00 | 686 | dataset1 | 0 | 0 | NA | 0 | Ap2n-kaz |
| MT791767 | Ap2n | 2n | MAH | Ap2nMAH14 | This study | No | 0.00 | 686 | dataset1 | 0 | 0 | NA | 0 | Ap2n-kaz |
| MT791768 | Ap2n | 2n | MAH | Ap2nMAH7 | This study | No | 0.00 | 686 | dataset1 | 0 | 0 | NA | 0 | Ap2n-kaz |
| MT791773 | Ap2n | 2n | NAR | Ap2nNAR2 | This study | No | 0.00 | 684 | dataset1 | 0 | 0 | NA | 0 | Ap2n-kaz |
| MT791774 | Ap2n | 2n | ODI | Ap2nODI11 | This study | No | 0.00 | 685 | dataset1 | 0 | 0 | NA | 0 | Ap2n-kaz |
| MT791775 | Ap2n | 2n | ODI | Ap2nODI12 | This study | No | 0.00 | 684 | dataset1 | 0 | 0 | NA | 0 | Ap2n-kaz |
| MT791776 | Ap2n | 2n | ODI | Ap2nODI13 | This study | No | 0.00 | 685 | dataset1 | 0 | 0 | NA | 0 | Ap2n-kaz |
| MT791777 | Ap2n | 2n | ODI | Ap2nODI14 | This study | No | 0.00 | 685 | dataset1 | 0 | 0 | NA | 0 | Ap2n-kaz |
| MT791705 | Ap2n | 2n | BOL | Ap2nBOL2 | This study | No | 0.00 | 686 | dataset1 | 0 | 0 | 1 | 0 | Ap2n-kaz |
| MT791706 | Ap2n | 2n | BOL | Ap2nBOL3 | This study | No | 0.00 | 686 | dataset1 | 0 | 0 | NA | 0 | Ap2n-kaz |
| MT791707 | Ap2n | 2n | BOL | Ap2nBOL4 | This study | No | 0.00 | 688 | dataset1 | 0 | 0 | NA | 0 | Ap2n-kaz |
| MT791708 | Ap2n | 2n | BOL | Ap2nBOL5 | This study | No | 0.00 | 685 | dataset1 | 0 | 0 | NA | 0 | Ap2n-kaz |
| NA | Ap2n | 2n | BOL | Ap2nBOL1 | This study | No | 0.01 | 523 | dataset2 | NA | NA | NA | 0 | Ap2n-kaz |
| NA | Ap2n | 2n | BOL | Ap2nBOL6 | This study | No | 0.00 | 669 | dataset2 | NA | NA | NA | 1 | Ap2n |
| MT791650 | Ap2n | 2n | AIM | Ap2nAIM58 | This study | No | 0.00 | 686 | dataset1 | 0 | 0 | NA | 0 | Ap2n-kaz |
| MT791651 | Ap2n | 2n | AIM | Ap2nAIM59 | This study | No | 0.00 | 685 | dataset1 | 0 | 0 | NA | 0 | Ap2n-kaz |
| MT791652 | Ap2n | 2n | AIM | Ap2nAIM60 | This study | No | 0.00 | 685 | dataset1 | 0 | 0 | NA | 0 | Ap2n-kaz |
| MT791653 | Ap2n | 2n | AIM | Ap2nAIM61 | This study | No | 0.00 | 685 | dataset1 | 0 | 0 | NA | 0 | Ap2n-kaz |
| MT791654 | Ap2n | 2n | AIM | Ap2nAIM62 | This study | No | 0.00 | 685 | dataset1 | 0 | 0 | NA | 0 | Ap2n-kaz |
| MT791655 | Ap2n | 2n | AIM | Ap2nAIM63 | This study | No | 0.00 | 686 | dataset1 | 0 | 0 | NA | 0 | Ap2n-kaz |
| MT791656 | Ap2n | 2n | AIM | Ap2nAIM64 | This study | No | 0.00 | 685 | dataset1 | 0 | 0 | NA | 0 | Ap2n-kaz |
| MT791657 | Ap2n | 2n | AIM | Ap2nAIM65 | This study | No | 0.00 | 685 | dataset1 | 0 | 0 | NA | 0 | Ap2n-kaz |
| MT791658 | Ap2n | 2n | AIM | Ap2nAIM66 | This study | No | 0.00 | 685 | dataset1 | 0 | 0 | NA | 0 | Ap2n-kaz |
| MT791659 | Ap2n | 2n | AIM | Ap2nAIM67 | This study | No | 0.00 | 686 | dataset1 | 0 | 0 | NA | 0 | Ap2n-kaz |
| MT791660 | Ap2n | 2n | AIM | Ap2nAIM68 | This study | No | 0.00 | 685 | dataset1 | 0 | 0 | NA | 0 | Ap2n-kaz |
| NA | Ap2n | 2n | AIM | Ap2nAIM69 | This study | No | 0.00 | 686 | dataset2 | NA | NA | NA | 1 | Ap2n |
| MT791785 | Ap2n | 2n | SAG | Ap2nSAG7 | This study | No | 0.00 | 681 | dataset1 | 0 | 0 | NA | 0 | Ap2n-urm |
| MT791786 | Ap2n | 2n | SAG | Ap2nSAG8 | This study | No | 0.00 | 664 | dataset1 | 0 | 0 | NA | 0 | Ap2n-urm |
| NA | Ap2n | 2n | SAG | Ap2nSAG11 | This study | No | 0.00 | 686 | dataset2 | NA | NA | NA | 0 | Ap2n-urm |
| MT791787 | Ap2n | 2n | SAG | Ap2nSAG13 | This study | No | 0.00 | 686 | dataset1 | 0 | 0 | NA | 0 | Ap2n-urm |
| MT791788 | Ap2n | 2n | SAG | Ap2nSAG14 | This study | No | 0.00 | 686 | dataset1 | 0 | 0 | NA | 0 | Ap2n-urm |
| MT791795 | Ap2n | 2n | YIN | Ap2nYIN1 | This study | No | 0.00 | 686 | dataset1 | 0 | 0 | NA | 0 | Ap2n-kaz |
| MT791796 | Ap2n | 2n | YIN | Ap2nYIN4 | This study | No | 0.00 | 683 | dataset1 | 0 | 0 | NA | 0 | Ap2n-kaz |
| NA | Ap2n | 2n | ATA1 | Ap2nATA12 | This study | No | 0.01 | 630 | dataset2 | NA | NA | NA | 1 | Ap2n |
| MT791721 | Ap2n | 2n | DON | Ap2nDON16 | This study | No | 0.00 | 615 | dataset1 | 0 | 0 | NA | 0 | Ap2n-kaz |
| MT791722 | Ap2n | 2n | DON | Ap2nDON18 | This study | No | 0.00 | 633 | dataset1 | 0 | 0 | NA | 0 | Ap2n-kaz |
| MT791723 | Ap2n | 2n | DON | Ap2nDON20 | This study | No | 0.00 | 634 | dataset1 | 0 | 0 | NA | 0 | Ap2n-kaz |
| NA | Ap2n | 2n | DON | Ap2nDON7 | This study | No | 0.00 | 618 | dataset2 | NA | NA | NA | 0 | Ap2n-kaz |
| NA | Ap2n | 2n | NAR | Ap2nNAR3 | This study | No | 0.00 | 634 | dataset2 | NA | NA | NA | 1 | Ap2n |
| MT791778 | Ap2n | 2n | ODI | Ap2nODI2 | This study | No | 0.00 | 632 | dataset1 | 0 | 0 | 1 | 0 | Ap2n-kaz |
| MT791779 | Ap2n | 2n | ODI | Ap2nODI4 | This study | No | 0.00 | 642 | dataset1 | 0 | 0 | NA | 0 | Ap2n-kaz |
| MT791780 | Ap2n | 2n | ODI | Ap2nODI6 | This study | No | 0.00 | 632 | dataset1 | 0 | 0 | NA | 0 | Ap2n-kaz |
| MT791781 | Ap2n | 2n | ODI | Ap2nODI8 | This study | No | 0.00 | 632 | dataset1 | 0 | 0 | NA | 0 | Ap2n-kaz |
| MT791661 | Ap2n | 2n | AIM | Ap2nAIM7 | This study | No | 0.00 | 633 | dataset1 | 0 | 0 | NA | 0 | Ap2n-kaz |
| MT791662 | Ap2n | 2n | AIM | Ap2nAIM9 | This study | No | 0.00 | 633 | dataset1 | 0 | 0 | NA | 0 | Ap2n-kaz |
| NA | Ap2n | 2n | VIL | Ap2nVIL10 | This study | No | 0.00 | 633 | dataset2 | NA | NA | NA | 1 | Ap2n |
| NA | Ap2n | 2n | VIL | Ap2nVIL12 | This study | No | 0.00 | 642 | dataset2 | NA | NA | NA | 0 | Ap2n-urm |
| NA | Ap2n | 2n | VIL | Ap2nVIL3 | This study | No | 0.00 | 629 | dataset2 | NA | NA | NA | 0 | Ap2n-urm |
| MT791792 | Ap2n | 2n | VIL | Ap2nVIL5 | This study | No | 0.00 | 649 | dataset1 | 0 | 0 | NA | 0 | Ap2n-urm |
| MT791789 | Ap2n | 2n | VIL | Ap2nVIL8 | This study | No | 0.00 | 633 | dataset1 | 0 | 0 | NA | 0 | Ap2n-kaz |
| NA | Ap2n | 2n | ATA1 | Ap2nATA17 | This study | No | 0.01 | 643 | dataset2 | NA | NA | NA | 0 | Ap2n-kaz |
| MT791724 | Ap2n | 2n | DON | Ap2nDON17 | This study | No | 0.00 | 634 | dataset1 | 0 | 0 | NA | 0 | Ap2n-kaz |
| NA | Ap2n | 2n | DON | Ap2nDON19 | This study | No | 0.00 | 621 | dataset2 | NA | NA | NA | 0 | Ap2n-kaz |
| MT791725 | Ap2n | 2n | DON | Ap2nDON21 | This study | No | 0.00 | 643 | dataset1 | 0 | 0 | NA | 0 | Ap2n-kaz |
| MT791726 | Ap2n | 2n | DON | Ap2nDON13 | This study | No | 0.00 | 643 | dataset1 | 0 | 0 | NA | 0 | Ap2n-kaz |
| MT791709 | Ap2n | 2n | COQ | Ap2nCOQ1 | This study | No | 0.00 | 637 | dataset1 | 0 | 0 | NA | 0 | Ap2n-kaz |
| MT791727 | Ap2n | 2n | KUL | Ap2nKUL5 | This study | No | 0.00 | 644 | dataset1 | 0 | 0 | NA | 0 | Ap2n-kaz |
| MT791728 | Ap2n | 2n | KUL | Ap2nKUL6 | This study | No | 0.00 | 636 | dataset1 | 0 | 0 | 1 | 0 | Ap2n-kaz |
| NA | Ap2n | 2n | MOL | Ap2nMOL9 | This study | No | 0.00 | 621 | dataset2 | NA | NA | NA | 0 | Ap2n-urm |
| NA | Ap2n | 2n | ODI | Ap2nODI7 | This study | No | 0.00 | 633 | dataset2 | NA | NA | NA | 0 | Ap2n-kaz |
| MT791782 | Ap2n | 2n | ODI | Ap2nODI9 | This study | No | 0.00 | 633 | dataset1 | 0 | 0 | NA | 0 | Ap2n-kaz |
| NA | Ap2n | 2n | AIM | Ap2nAIM6 | This study | No | 0.00 | 633 | dataset2 | NA | NA | NA | 1 | Ap2n |
| MT791663 | Ap2n | 2n | AIM | Ap2nAIM8 | This study | No | 0.00 | 633 | dataset1 | 0 | 0 | NA | 0 | Ap2n-kaz |
| MT791793 | Ap2n | 2n | VIL | Ap2nVIL1 | This study | No | 0.00 | 633 | dataset1 | 0 | 0 | NA | 0 | Ap2n-urm |
| NA | Ap2n | 2n | VIL | Ap2nVIL11 | This study | No | 0.00 | 633 | dataset2 | NA | NA | NA | 1 | Ap2n |
| MT791794 | Ap2n | 2n | VIL | Ap2nVIL2 | This study | No | 0.00 | 633 | dataset1 | 0 | 0 | NA | 0 | Ap2n-urm |
| NA | Ap2n | 2n | VIL | Ap2nVIL4 | This study | No | 0.00 | 633 | dataset2 | NA | NA | NA | 1 | Ap2n |
| MT791790 | Ap2n | 2n | VIL | Ap2nVIL7 | This study | No | 0.00 | 638 | dataset1 | 0 | 0 | NA | 0 | Ap2n-kaz |
| MT791791 | Ap2n | 2n | VIL | Ap2nVIL9 | This study | No | 0.00 | 633 | dataset1 | 0.09 | 0 | 1 | 0 | Ap2n-kaz |
| NA | Ap2n | 2n | VIL | Ap2nVIL6 | This study | No | 0.00 | 304 | dataset2 | NA | NA | NA | 1 | Ap2n |
| MT792814 | Ap2n | 2n | AIM | numt1Ap2nAIM7 | This study | Yes | 0.00 | 572 | dataset3 | 1.88 | 1 | 1 | 1 | Ap2n |
| NA | Ap2n | 2n | AIM | numt2Ap2nAIM7 | This study | Yes | 0.00 | 572 | dataset4 | NA | NA | NA | 0 | Ap2n-kaz |
| MT792815 | Ap2n | 2n | MOL | numt2Ap2nMOL10 | This study | Yes | 0.00 | 587 | dataset3 | 1.36 | 1 | 0 | 1 | Ap2n |
| MT792816 | Ap2n | 2n | MOL | numt1Ap2nMOL10 | This study | Yes | 0.00 | 590 | dataset3 | 1.4 | 1 | 1 | 1 | Ap2n |
| MT791772 | Ap2n | 2n | MOL | Ap2nMOL10 | This study | Yes | 0.00 | 590 | dataset3 | 0 | 0 | NA | 0 | Ap2n-urm |
| DQ426824 | Ap2n | 2n | NA | APD01 | GenBank | No | 0.00 | 658 | dataset5 | 0 | 0 | 1 | 0 | Ap2n-kaz |
| DQ426825 | Ap2n | 2n | NA | APD02 | GenBank | No | 0.00 | 658 | dataset5 | 0 | 0 | NA | 0 | Ap2n-kaz |
| DQ426826 | Ap2n | 2n | NA | APD03 | GenBank | No | 0.00 | 658 | dataset5 | 0 | 0 | 1 | 0 | Ap2n-kaz |
| GU591380 | Ap2n | 2n | NA | APD04 | GenBank | No | 0.00 | 658 | dataset5 | 0 | 0 | 1 | 0 | Ap2n-kaz |
| GU591381 | Ap2n | 2n | NA | APD05 | GenBank | No | 0.00 | 658 | dataset5 | 0 | 0 | 1 | 0 | Ap2n-kaz |
| GU591382 | Ap2n | 2n | NA | APD06 | GenBank | No | 0.00 | 658 | dataset5 | 0.18 | 1 | 1 | 1 | Ap2n |
| GU591383 | Ap2n | 2n | NA | APD07 | GenBank | No | 0.00 | 658 | dataset5 | 0.06 | 0 | 1 | 0 | Ap2n-urm |
| GU591384 | Ap2n | 2n | NA | APD08 | GenBank | No | 0.00 | 658 | dataset5 | 0 | 0 | 1 | 0 | Ap2n-urm |
| MG572052 | Ap2n | 2n | NA | Tanggu1990 | GenBank | No | 0.00 | 1406 | dataset6 | NA | NA | NA | 0 | Ap2n-kaz |
| MG572053 | Ap2n | 2n | NA | Aibi1991 | GenBank | No | 0.00 | 1407 | dataset6 | NA | NA | NA | 0 | Ap2n-kaz |
| MG572054 | Ap2n | 2n | NA | Aibi2011 | GenBank | No | 0.00 | 1418 | dataset6 | NA | NA | NA | 1 | Ap2n |
| MG572055 | Ap2n | 2n | NA | Aibi2016 | GenBank | No | 0.00 | 1419 | dataset6 | NA | NA | NA | 1 | Ap2n |
| MG572056 | Ap2n | 2n | NA | Gahai1991 | GenBank | No | 0.00 | 1406 | dataset6 | NA | NA | NA | 0 | Ap2n-kaz |
| MG572057 | Ap2n | 2n | NA | Borli2015 | GenBank | No | 0.00 | 1412 | dataset6 | NA | NA | NA | 1 | Ap2n |
| MG572058 | Ap2n | 2n | NA | Setine2015 | GenBank | No | 0.00 | 1346 | dataset6 | NA | NA | NA | 0 | Ap2n-kaz |
| MG572059 | Ap2n | 2n | NA | Aral2017 | GenBank | No | 0.00 | 1351 | dataset6 | NA | NA | NA | 0 | Ap2n-kaz |
| MG572060 | Ap2n | 2n | NA | Yarovoe2015 | GenBank | No | 0.00 | 1406 | dataset6 | NA | NA | NA | 0 | Ap2n-kaz |
| MG572061 | Ap2n | 2n | NA | Kunlundinskoe2014 | GenBank | No | 0.00 | 1406 | dataset6 | NA | NA | NA | 0 | Ap2n-kaz |
| MG572062 | Ap2n | 2n | NA | Medvejie2011 | GenBank | No | 0.00 | 1406 | dataset6 | NA | NA | NA | 0 | Ap2n-kaz |
| MG572063 | Ap2n | 2n | NA | Medvejie2015 | GenBank | No | 0.00 | 1406 | dataset6 | NA | NA | NA | 0 | Ap2n-kaz |
| MG572064 | Ap2n | 2n | NA | Ebeity2011 | GenBank | No | 0.00 | 1331 | dataset6 | NA | NA | NA | 0 | Ap2n-kaz |
| MG572065 | Ap2n | 2n | NA | Hangu1990 | GenBank | No | 0.00 | 1418 | dataset6 | NA | NA | NA | 1 | Ap2n |
| MG572066 | Ap2n | 2n | NA | Gaize1998 | GenBank | No | 0.00 | 1406 | dataset6 | NA | NA | NA | 0 | Ap2n-urm |
| KF707700 | Ap2n | 2n | NA | KOY1 | GenBank | No | 0.00 | 614 | dataset5 | 0 | 0 | NA | 0 | Ap2n-kaz |
| KF707701 | Ap2n | 2n | NA | KOY6 | GenBank | No | 0.00 | 614 | dataset5 | 0 | 0 | NA | 0 | Ap2n-kaz |
| KF707702 | Ap2n | 2n | NA | KOY10 | GenBank | No | 0.00 | 614 | dataset5 | 0 | 0 | NA | 0 | Ap2n-kaz |
| KF707703 | Ap2n | 2n | NA | KOY9 | GenBank | No | 0.00 | 614 | dataset5 | 0 | 0 | NA | 0 | Ap2n-kaz |
| KF707704 | Ap2n | 2n | NA | KOY8 | GenBank | No | 0.00 | 614 | dataset5 | 0 | 0 | NA | 0 | Ap2n-kaz |
| KF707705 | Ap2n | 2n | NA | KOY7 | GenBank | No | 0.00 | 614 | dataset5 | 0 | 0 | NA | 0 | Ap2n-kaz |
| KF707706 | Ap2n | 2n | NA | KOY5 | GenBank | No | 0.00 | 614 | dataset5 | 0 | 0 | NA | 0 | Ap2n-kaz |
| KF707707 | Ap2n | 2n | NA | KOY4 | GenBank | No | 0.00 | 614 | dataset5 | 0 | 0 | NA | 0 | Ap2n-kaz |
| KF707708 | Ap2n | 2n | NA | KOY3 | GenBank | No | 0.00 | 614 | dataset5 | 0 | 0 | NA | 0 | Ap2n-kaz |
| KF707709 | Ap2n | 2n | NA | KOY2 | GenBank | No | 0.00 | 614 | dataset5 | 0 | 0 | NA | 0 | Ap2n-kaz |
| KF707710 | Ap2n | 2n | NA | URM8 | GenBank | No | 0.00 | 614 | dataset5 | 0 | 0 | NA | 0 | Ap2n-kaz |
| KF707711 | Ap2n | 2n | NA | URM15 | GenBank | No | 0.00 | 614 | dataset5 | 0 | 0 | NA | 0 | Ap2n-kaz |
| KF707712 | Ap2n | 2n | NA | URM14 | GenBank | No | 0.00 | 614 | dataset5 | 0 | 0 | NA | 0 | Ap2n-kaz |
| KF707713 | Ap2n | 2n | NA | URM13 | GenBank | No | 0.00 | 614 | dataset5 | 0 | 0 | NA | 0 | Ap2n-kaz |
| KF707714 | Ap2n | 2n | NA | URM11 | GenBank | No | 0.00 | 614 | dataset5 | 0 | 0 | NA | 0 | Ap2n-kaz |
| KF707715 | Ap2n | 2n | NA | URM10 | GenBank | No | 0.00 | 614 | dataset5 | 0 | 0 | NA | 0 | Ap2n-kaz |
| KF707716 | Ap2n | 2n | NA | URM7 | GenBank | No | 0.00 | 614 | dataset5 | 0 | 0 | NA | 0 | Ap2n-kaz |
| KF707717 | Ap2n | 2n | NA | URM6 | GenBank | No | 0.00 | 614 | dataset5 | 0 | 0 | NA | 0 | Ap2n-kaz |
| KF707718 | Ap2n | 2n | NA | URM5 | GenBank | No | 0.00 | 614 | dataset5 | 0 | 0 | NA | 0 | Ap2n-kaz |
| KF707719 | Ap2n | 2n | NA | URM4 | GenBank | No | 0.00 | 614 | dataset5 | 0 | 0 | NA | 0 | Ap2n-kaz |
| KF707720 | Ap2n | 2n | NA | ATA7 | GenBank | No | 0.00 | 614 | dataset5 | 0 | 0 | NA | 0 | Ap2n-kaz |
| KF707721 | Ap2n | 2n | NA | ATA6 | GenBank | No | 0.00 | 614 | dataset5 | 0 | 0 | NA | 0 | Ap2n-kaz |
| KF707722 | Ap2n | 2n | NA | ATA5 | GenBank | No | 0.00 | 614 | dataset5 | 0 | 0 | NA | 0 | Ap2n-kaz |
| KF707723 | Ap2n | 2n | NA | ATA4 | GenBank | No | 0.00 | 614 | dataset5 | 0 | 0 | NA | 0 | Ap2n-kaz |
| KF707724 | Ap2n | 2n | NA | ATA3 | GenBank | No | 0.00 | 614 | dataset5 | 0 | 0 | NA | 0 | Ap2n-kaz |
| KF707725 | Ap2n | 2n | NA | ATA2 | GenBank | No | 0.00 | 614 | dataset5 | 0 | 0 | NA | 0 | Ap2n-kaz |
| KF707726 | Ap2n | 2n | NA | ATA1 | GenBank | No | 0.00 | 614 | dataset5 | 0.06 | 0 | NA | 0 | Ap2n-urm |
| KF707727 | Ap2n | 2n | NA | IRA3 | GenBank | No | 0.00 | 614 | dataset5 | 0 | 0 | NA | 0 | Ap2n-kaz |
| KF707728 | Ap2n | 2n | NA | IRA4 | GenBank | No | 0.00 | 614 | dataset5 | 0 | 0 | NA | 0 | Ap2n-kaz |
| KF707729 | Ap2n | 2n | NA | IRA5 | GenBank | No | 0.00 | 614 | dataset5 | 0 | 0 | NA | 0 | Ap2n-kaz |
| KF707730 | Ap2n | 2n | NA | IRA12 | GenBank | No | 0.00 | 614 | dataset5 | 0 | 0 | NA | 0 | Ap2n-kaz |
| KF707731 | Ap2n | 2n | NA | IRA13 | GenBank | No | 0.00 | 614 | dataset5 | 0 | 0 | NA | 0 | Ap2n-kaz |
| KF707732 | Ap2n | 2n | NA | IRA2 | GenBank | No | 0.00 | 614 | dataset5 | 0 | 0 | NA | 0 | Ap2n-kaz |
| KF707733 | Ap2n | 2n | NA | IRA7 | GenBank | No | 0.00 | 614 | dataset5 | 0 | 0 | NA | 0 | Ap2n-kaz |
| KF707734 | Ap2n | 2n | NA | IRA8 | GenBank | No | 0.00 | 614 | dataset5 | 0 | 0 | NA | 0 | Ap2n-kaz |
| KF707735 | Ap2n | 2n | NA | IRA9 | GenBank | No | 0.00 | 614 | dataset5 | 0 | 0 | NA | 0 | Ap2n-kaz |
| KF707736 | Ap2n | 2n | NA | IRA6 | GenBank | No | 0.00 | 614 | dataset5 | 0 | 0 | NA | 0 | Ap2n-kaz |
| KF707737 | Ap2n | 2n | NA | IRA22 | GenBank | No | 0.00 | 614 | dataset5 | 0 | 0 | NA | 0 | Ap2n-kaz |
| KF707738 | Ap2n | 2n | NA | IRA15 | GenBank | No | 0.00 | 614 | dataset5 | 0 | 0 | NA | 0 | Ap2n-kaz |
| KF707739 | Ap2n | 2n | NA | IRA17 | GenBank | No | 0.00 | 614 | dataset5 | 0 | 0 | NA | 0 | Ap2n-kaz |
| KF707740 | Ap2n | 2n | NA | IRA18 | GenBank | No | 0.00 | 614 | dataset5 | 0 | 0 | NA | 0 | Ap2n-kaz |
| KF707741 | Ap2n | 2n | NA | IRA20 | GenBank | No | 0.00 | 614 | dataset5 | 0 | 0 | NA | 0 | Ap2n-kaz |
| KF707742 | Ap2n | 2n | NA | IRA14 | GenBank | No | 0.00 | 614 | dataset5 | 0 | 0 | NA | 0 | Ap2n-kaz |
| KF707743 | Ap2n | 2n | NA | IRA16 | GenBank | No | 0.00 | 614 | dataset5 | 0 | 0 | NA | 0 | Ap2n-kaz |
| KF707744 | Ap2n | 2n | NA | IRA19 | GenBank | No | 0.00 | 614 | dataset5 | 0 | 0 | NA | 0 | Ap2n-kaz |
| KF707745 | Ap2n | 2n | NA | IRA21 | GenBank | No | 0.00 | 614 | dataset5 | 0 | 0 | NA | 0 | Ap2n-kaz |
| KF707746 | Ap2n | 2n | NA | AIB3 | GenBank | No | 0.00 | 614 | dataset5 | 0 | 0 | 2 | 1 | Ap2n |
| KF707747 | Ap2n | 2n | NA | AIB9 | GenBank | No | 0.00 | 614 | dataset5 | 0 | 0 | NA | 0 | Ap2n-kaz |
| KF707748 | Ap2n | 2n | NA | AIB6 | GenBank | No | 0.00 | 614 | dataset5 | 0 | 0 | NA | 0 | Ap2n-kaz |
| KF707749 | Ap2n | 2n | NA | AIB10 | GenBank | No | 0.00 | 614 | dataset5 | 0 | 0 | NA | 0 | Ap2n-urm |
| KF707750 | Ap2n | 2n | NA | AIB4 | GenBank | No | 0.00 | 614 | dataset5 | 0 | 0 | NA | 0 | Ap2n-kaz |
| KF707751 | Ap2n | 2n | NA | AIB7 | GenBank | No | 0.00 | 614 | dataset5 | 0 | 0 | NA | 0 | Ap2n-urm |
| KF707752 | Ap2n | 2n | NA | AIB8 | GenBank | No | 0.00 | 614 | dataset5 | 0 | 0 | NA | 0 | Ap2n-urm |
| KF707753 | Ap2n | 2n | NA | AIB2 | GenBank | No | 0.00 | 614 | dataset5 | 0 | 0 | NA | 0 | Ap2n-kaz |
| KF707754 | Ap2n | 2n | NA | AIB1 | GenBank | No | 0.00 | 614 | dataset5 | 0 | 0 | NA | 0 | Ap2n-kaz |
| KF707755 | Ap2n | 2n | NA | GAH3 | GenBank | No | 0.00 | 614 | dataset5 | 0 | 0 | 1 | 0 | Ap2n-kaz |
| KF707756 | Ap2n | 2n | NA | GAH10 | GenBank | No | 0.00 | 614 | dataset5 | 0 | 0 | NA | 0 | Ap2n-kaz |
| KF707757 | Ap2n | 2n | NA | GAH5 | GenBank | No | 0.00 | 614 | dataset5 | 0 | 0 | NA | 0 | Ap2n-kaz |
| KF707758 | Ap2n | 2n | NA | GAH4 | GenBank | No | 0.00 | 614 | dataset5 | 0 | 0 | NA | 0 | Ap2n-kaz |
| KF707759 | Ap2n | 2n | NA | GAH1 | GenBank | No | 0.00 | 614 | dataset5 | 0 | 0 | NA | 0 | Ap2n-kaz |
| KF707760 | Ap2n | 2n | NA | GAH14 | GenBank | No | 0.00 | 614 | dataset5 | 0 | 0 | NA | 0 | Ap2n-kaz |
| KF707761 | Ap2n | 2n | NA | GAH6 | GenBank | No | 0.00 | 614 | dataset5 | 0 | 0 | NA | 0 | Ap2n-kaz |
| KF707762 | Ap2n | 2n | NA | GAH7 | GenBank | No | 0.00 | 614 | dataset5 | 0 | 0 | NA | 0 | Ap2n-kaz |
| KF707763 | Ap2n | 2n | NA | GAH8 | GenBank | No | 0.00 | 614 | dataset5 | 0 | 0 | NA | 0 | Ap2n-kaz |
| KF707764 | Ap2n | 2n | NA | GAH9 | GenBank | No | 0.00 | 614 | dataset5 | 0 | 0 | NA | 0 | Ap2n-kaz |
| KF707765 | Ap2n | 2n | NA | URM21 | GenBank | No | 0.00 | 614 | dataset5 | 0 | 0 | NA | 0 | Ap2n-kaz |
| KF707766 | Ap2n | 2n | NA | URM22 | GenBank | No | 0.00 | 614 | dataset5 | 0 | 0 | NA | 0 | Ap2n-kaz |
| KF707767 | Ap2n | 2n | NA | URM20 | GenBank | No | 0.00 | 614 | dataset5 | 0 | 0 | NA | 0 | Ap2n-kaz |
| KF707768 | Ap2n | 2n | NA | URM23 | GenBank | No | 0.00 | 614 | dataset5 | 0 | 0 | NA | 0 | Ap2n-kaz |
| KF707769 | Ap2n | 2n | NA | URM24 | GenBank | No | 0.00 | 614 | dataset5 | 0 | 0 | NA | 0 | Ap2n-kaz |
| KF707770 | Ap2n | 2n | NA | URM17 | GenBank | No | 0.00 | 614 | dataset5 | 0 | 0 | NA | 0 | Ap2n-kaz |
| KF707771 | Ap2n | 2n | NA | URM16 | GenBank | No | 0.00 | 614 | dataset5 | 0 | 0 | NA | 0 | Ap2n-kaz |
| KF707772 | Ap2n | 2n | NA | URM18 | GenBank | No | 0.00 | 614 | dataset5 | 0 | 0 | NA | 0 | Ap2n-kaz |
| KF707773 | Ap2n | 2n | NA | URM19 | GenBank | No | 0.00 | 614 | dataset5 | 0 | 0 | NA | 0 | Ap2n-kaz |
| KF707774 | Ap2n | 2n | NA | URM25 | GenBank | No | 0.00 | 614 | dataset5 | 0 | 0 | NA | 0 | Ap2n-kaz |
| KF707775 | Ap2n | 2n | NA | PAK14 | GenBank | No | 0.00 | 614 | dataset5 | 0 | 0 | NA | 0 | Ap2n-kaz |
| KF707776 | Ap2n | 2n | NA | PAK11 | GenBank | No | 0.00 | 614 | dataset5 | 0 | 0 | NA | 0 | Ap2n-kaz |
| KF707777 | Ap2n | 2n | NA | PAK12 | GenBank | No | 0.00 | 614 | dataset5 | 0 | 0 | NA | 0 | Ap2n-kaz |
| KF707778 | Ap2n | 2n | NA | PAK13 | GenBank | No | 0.00 | 614 | dataset5 | 0 | 0 | NA | 0 | Ap2n-kaz |
| KF707779 | Ap2n | 2n | NA | PAK2 | GenBank | No | 0.00 | 614 | dataset5 | 0 | 0 | NA | 0 | Ap2n-kaz |
| KF707780 | Ap2n | 2n | NA | PAK3 | GenBank | No | 0.00 | 614 | dataset5 | 0 | 0 | NA | 0 | Ap2n-kaz |
| KF707781 | Ap2n | 2n | NA | PAK4 | GenBank | No | 0.00 | 614 | dataset5 | 0 | 0 | NA | 0 | Ap2n-kaz |
| KF707782 | Ap2n | 2n | NA | PAK5 | GenBank | No | 0.00 | 614 | dataset5 | 0 | 0 | NA | 0 | Ap2n-kaz |
| KF707783 | Ap2n | 2n | NA | PAK6 | GenBank | No | 0.00 | 614 | dataset5 | 0 | 0 | NA | 0 | Ap2n-kaz |
| KF707784 | Ap2n | 2n | NA | PAK7 | GenBank | No | 0.00 | 614 | dataset5 | 0 | 0 | NA | 0 | Ap2n-kaz |
| KF707785 | Ap2n | 2n | NA | EGY1 | GenBank | No | 0.00 | 614 | dataset5 | 0 | 0 | NA | 0 | Ap2n-kaz |
| KF707786 | Ap2n | 2n | NA | EGY5 | GenBank | No | 0.00 | 614 | dataset5 | 0 | 0 | NA | 0 | Ap2n-kaz |
| KF707787 | Ap2n | 2n | NA | EGY10 | GenBank | No | 0.00 | 614 | dataset5 | 0 | 0 | NA | 0 | Ap2n-kaz |
| KF707788 | Ap2n | 2n | NA | EGY4 | GenBank | No | 0.00 | 614 | dataset5 | 0 | 0 | NA | 0 | Ap2n-kaz |
| KF707789 | Ap2n | 2n | NA | EGY9 | GenBank | No | 0.00 | 614 | dataset5 | 0 | 0 | NA | 0 | Ap2n-kaz |
| KF707790 | Ap2n | 2n | NA | ALB5 | GenBank | No | 0.00 | 614 | dataset5 | 0 | 0 | NA | 0 | Ap2n-kaz |
| KF707791 | Ap2n | 2n | NA | ALB9 | GenBank | No | 0.00 | 614 | dataset5 | 0 | 0 | NA | 0 | Ap2n-kaz |
| KF707792 | Ap2n | 2n | NA | ALB1 | GenBank | No | 0.00 | 614 | dataset5 | 0 | 0 | NA | 0 | Ap2n-kaz |
| KF707793 | Ap2n | 2n | NA | ALB2 | GenBank | No | 0.00 | 614 | dataset5 | 0 | 0 | NA | 0 | Ap2n-kaz |
| KF707794 | Ap2n | 2n | NA | ALB3 | GenBank | No | 0.00 | 614 | dataset5 | 0 | 0 | NA | 0 | Ap2n-kaz |
| KF707795 | Ap2n | 2n | NA | ALB4 | GenBank | No | 0.00 | 614 | dataset5 | 0 | 0 | NA | 0 | Ap2n-kaz |
| KF707796 | Ap2n | 2n | NA | ALB6 | GenBank | No | 0.00 | 614 | dataset5 | 0 | 0 | NA | 0 | Ap2n-kaz |
| KF707797 | Ap2n | 2n | NA | ALB7 | GenBank | No | 0.00 | 614 | dataset5 | 0 | 0 | NA | 0 | Ap2n-kaz |
| KF707798 | Ap2n | 2n | NA | ALB8 | GenBank | No | 0.00 | 614 | dataset5 | 0 | 0 | NA | 0 | Ap2n-kaz |
| KF707799 | Ap2n | 2n | NA | ALB10 | GenBank | No | 0.00 | 614 | dataset5 | 0 | 0 | NA | 0 | Ap2n-kaz |
| KF707800 | Ap2n | 2n | NA | ATA8 | GenBank | No | 0.00 | 614 | dataset5 | 0 | 0 | NA | 0 | Ap2n-kaz |
| KF707801 | Ap2n | 2n | NA | ATA15 | GenBank | No | 0.00 | 614 | dataset5 | 0 | 0 | NA | 0 | Ap2n-urm |
| KF707802 | Ap2n | 2n | NA | ATA10 | GenBank | No | 0.00 | 614 | dataset5 | 0 | 0 | NA | 0 | Ap2n-kaz |
| KF707803 | Ap2n | 2n | NA | ATA11 | GenBank | No | 0.00 | 614 | dataset5 | 0 | 0 | NA | 0 | Ap2n-kaz |
| KF707804 | Ap2n | 2n | NA | ATA12 | GenBank | No | 0.00 | 614 | dataset5 | 0 | 0 | NA | 0 | Ap2n-kaz |
| KF707805 | Ap2n | 2n | NA | KOY11 | GenBank | No | 0.00 | 614 | dataset5 | 0 | 0 | NA | 0 | Ap2n-kaz |
| KF707806 | Ap2n | 2n | NA | KOY17 | GenBank | No | 0.00 | 614 | dataset5 | 0 | 0 | NA | 0 | Ap2n-kaz |
| KF707807 | Ap2n | 2n | NA | KOY18 | GenBank | No | 0.00 | 614 | dataset5 | 0 | 0 | NA | 0 | Ap2n-kaz |
| KF707808 | Ap2n | 2n | NA | KOY14 | GenBank | No | 0.00 | 614 | dataset5 | 0 | 0 | NA | 0 | Ap2n-kaz |
| KF707809 | Ap2n | 2n | NA | KOY16 | GenBank | No | 0.00 | 614 | dataset5 | 0 | 0 | NA | 0 | Ap2n-kaz |
| KF707810 | Ap2n | 2n | NA | OYB10 | GenBank | No | 0.00 | 614 | dataset5 | 0 | 0 | NA | 0 | Ap2n-urm |
| KF707811 | Ap2n | 2n | NA | OYB3 | GenBank | No | 0.00 | 614 | dataset5 | 0 | 0 | NA | 0 | Ap2n-urm |
| KF707812 | Ap2n | 2n | NA | OYB4 | GenBank | No | 0.00 | 614 | dataset5 | 0 | 0 | NA | 0 | Ap2n-urm |
| KF707813 | Ap2n | 2n | NA | OYB5 | GenBank | No | 0.00 | 614 | dataset5 | 0 | 0 | NA | 0 | Ap2n-urm |
| KF707814 | Ap2n | 2n | NA | OYB6 | GenBank | No | 0.00 | 614 | dataset5 | 0 | 0 | NA | 0 | Ap2n-urm |
| KF707815 | Ap2n | 2n | NA | OYB9 | GenBank | No | 0.00 | 614 | dataset5 | 0 | 0 | NA | 0 | Ap2n-urm |
| KF707816 | Ap2n | 2n | NA | OYB1 | GenBank | No | 0.00 | 614 | dataset5 | 0 | 0 | NA | 0 | Ap2n-urm |
| KF707817 | Ap2n | 2n | NA | OYB13 | GenBank | No | 0.00 | 614 | dataset5 | 0 | 0 | NA | 0 | Ap2n-urm |
| KF707818 | Ap2n | 2n | NA | OYB7 | GenBank | No | 0.00 | 614 | dataset5 | 0 | 0 | NA | 0 | Ap2n-urm |
| KF707819 | Ap2n | 2n | NA | OYB8 | GenBank | No | 0.00 | 614 | dataset5 | 0 | 0 | NA | 0 | Ap2n-urm |
| KF707820 | Ap2n | 2n | NA | ARA1 | GenBank | No | 0.00 | 614 | dataset5 | 0 | 0 | NA | 0 | Ap2n-kaz |
| KF707821 | Ap2n | 2n | NA | ARA4 | GenBank | No | 0.00 | 614 | dataset5 | 0 | 0 | 1 | 0 | Ap2n-kaz |
| KF707822 | Ap2n | 2n | NA | ARA5 | GenBank | No | 0.00 | 614 | dataset5 | 0.06 | 0 | 1 | 0 | Ap2n-kaz |
| KF707823 | Ap2n | 2n | NA | ARA6 | GenBank | No | 0.00 | 614 | dataset5 | 0 | 0 | NA | 0 | Ap2n-kaz |
| KF707824 | Ap2n | 2n | NA | ARA7 | GenBank | No | 0.00 | 614 | dataset5 | 0 | 0 | NA | 0 | Ap2n-kaz |
| KF707825 | Ap2n | 2n | NA | ARA8 | GenBank | No | 0.00 | 614 | dataset5 | 0 | 0 | NA | 0 | Ap2n-kaz |
| KF707826 | Ap2n | 2n | NA | MAL2 | GenBank | No | 0.00 | 614 | dataset5 | 0 | 0 | NA | 0 | Ap2n-kaz |
| KF707827 | Ap2n | 2n | NA | MAL4 | GenBank | No | 0.00 | 614 | dataset5 | 0 | 0 | NA | 0 | Ap2n-kaz |
| KF707828 | Ap2n | 2n | NA | MAL1 | GenBank | No | 0.00 | 614 | dataset5 | 0 | 0 | NA | 0 | Ap2n-kaz |
| KF707829 | Ap2n | 2n | NA | MAL10 | GenBank | No | 0.00 | 614 | dataset5 | 0 | 0 | NA | 0 | Ap2n-kaz |
| KF707830 | Ap2n | 2n | NA | MAL3 | GenBank | No | 0.00 | 614 | dataset5 | 0 | 0 | NA | 0 | Ap2n-kaz |
| KF707831 | Ap2n | 2n | NA | MAL5 | GenBank | No | 0.00 | 614 | dataset5 | 0 | 0 | NA | 0 | Ap2n-kaz |
| KF707832 | Ap2n | 2n | NA | MAL6 | GenBank | No | 0.00 | 614 | dataset5 | 0 | 0 | NA | 0 | Ap2n-kaz |
| KF707833 | Ap2n | 2n | NA | MAL7 | GenBank | No | 0.00 | 614 | dataset5 | 0 | 0 | NA | 0 | Ap2n-kaz |
| KF707834 | Ap2n | 2n | NA | MAL8 | GenBank | No | 0.00 | 614 | dataset5 | 0 | 0 | NA | 0 | Ap2n-kaz |
| KF707835 | Ap2n | 2n | NA | MAL9 | GenBank | No | 0.00 | 614 | dataset5 | 0 | 0 | NA | 0 | Ap2n-kaz |
| KF707836 | Ap2n | 2n | NA | BOL1 | GenBank | No | 0.00 | 614 | dataset5 | 0 | 0 | NA | 0 | Ap2n-kaz |
| KF707837 | Ap2n | 2n | NA | BOL10 | GenBank | No | 0.00 | 614 | dataset5 | 0 | 0 | NA | 0 | Ap2n-kaz |
| KF707838 | Ap2n | 2n | NA | BOL2 | GenBank | No | 0.00 | 614 | dataset5 | 0 | 0 | NA | 0 | Ap2n-kaz |
| KF707839 | Ap2n | 2n | NA | BOL3 | GenBank | No | 0.00 | 614 | dataset5 | 0 | 0 | NA | 0 | Ap2n-kaz |
| KF707840 | Ap2n | 2n | NA | BOL4 | GenBank | No | 0.00 | 614 | dataset5 | 0 | 0 | NA | 0 | Ap2n-kaz |
| KF707841 | Ap2n | 2n | NA | BOL6 | GenBank | No | 0.00 | 614 | dataset5 | 0 | 0 | NA | 0 | Ap2n-kaz |
| KF707842 | Ap2n | 2n | NA | BOL7 | GenBank | No | 0.00 | 614 | dataset5 | 0 | 0 | NA | 0 | Ap2n-kaz |
| KF707843 | Ap2n | 2n | NA | BOL8 | GenBank | No | 0.00 | 614 | dataset5 | 0 | 0 | NA | 0 | Ap2n-kaz |
| KF707844 | Ap2n | 2n | NA | BOL9 | GenBank | No | 0.00 | 614 | dataset5 | 0 | 0 | NA | 0 | Ap2n-kaz |
| KF707845 | Ap2n | 2n | NA | LAG3 | GenBank | No | 0.00 | 614 | dataset5 | 0 | 0 | NA | 0 | Ap2n-kaz |
| KF707846 | Ap2n | 2n | NA | LAG2 | GenBank | No | 0.00 | 614 | dataset5 | 0 | 0 | NA | 0 | Ap2n-kaz |
| KF707847 | Ap2n | 2n | NA | LAG8 | GenBank | No | 0.00 | 614 | dataset5 | 1.36 | 1 | 1 | 1 | Ap2n |
| KF707848 | Ap2n | 2n | NA | LAG4 | GenBank | No | 0.00 | 614 | dataset5 | 1.36 | 1 | NA | 1 | Ap2n |
| KF707849 | Ap2n | 2n | NA | LAG6 | GenBank | No | 0.00 | 614 | dataset5 | 1.36 | 1 | NA | 1 | Ap2n |
| KF707850 | Ap2n | 2n | NA | LAG7 | GenBank | No | 0.00 | 614 | dataset5 | 0 | 0 | NA | 0 | Ap2n-kaz |
| KF707851 | Ap2n | 2n | NA | LAG10 | GenBank | No | 0.00 | 614 | dataset5 | 1.36 | 1 | NA | 1 | Ap2n |
| KF707852 | Ap2n | 2n | NA | LAG5 | GenBank | No | 0.00 | 614 | dataset5 | 0 | 0 | NA | 0 | Ap2n-kaz |
| KF707853 | Ap2n | 2n | NA | LAG1 | GenBank | No | 0.00 | 614 | dataset5 | 0 | 0 | NA | 0 | Ap2n-kaz |
| KF707854 | Ap2n | 2n | NA | LAG9 | GenBank | No | 0.00 | 614 | dataset5 | 1.36 | 1 | NA | 1 | Ap2n |
| KF707865 | Ap2n | 2n | NA | MOI10 | GenBank | No | 0.00 | 614 | dataset5 | 0 | 0 | NA | 0 | Ap2n-kaz |
| KF707866 | Ap2n | 2n | NA | MOI1 | GenBank | No | 0.00 | 614 | dataset5 | 0 | 0 | NA | 0 | Ap2n-kaz |
| KF707867 | Ap2n | 2n | NA | MOI2 | GenBank | No | 0.00 | 614 | dataset5 | 0 | 0 | NA | 0 | Ap2n-kaz |
| KF707868 | Ap2n | 2n | NA | MOI3 | GenBank | No | 0.00 | 614 | dataset5 | 0 | 0 | NA | 0 | Ap2n-kaz |
| KF707869 | Ap2n | 2n | NA | MOI4 | GenBank | No | 0.00 | 614 | dataset5 | 0 | 0 | NA | 0 | Ap2n-kaz |
| KF707870 | Ap2n | 2n | NA | MOI6 | GenBank | No | 0.00 | 614 | dataset5 | 0 | 0 | NA | 0 | Ap2n-kaz |
| KF707871 | Ap2n | 2n | NA | MOI5 | GenBank | No | 0.00 | 614 | dataset5 | 0 | 0 | 1 | 0 | Ap2n-kaz |
| KF707872 | Ap2n | 2n | NA | MOI7 | GenBank | No | 0.00 | 614 | dataset5 | 0 | 0 | NA | 0 | Ap2n-kaz |
| KF707873 | Ap2n | 2n | NA | MOI8 | GenBank | No | 0.00 | 614 | dataset5 | 0 | 0 | NA | 0 | Ap2n-kaz |
| KF707874 | Ap2n | 2n | NA | MOI9 | GenBank | No | 0.00 | 614 | dataset5 | 0 | 0 | NA | 0 | Ap2n-kaz |
| KC193638 | Ap2n | 2n | NA | rmMDS1 | GenBank | No | 0.00 | 617 | dataset5 | 0.09 | 0 | 0 | 0 | Ap2n-kaz |
| KC193639 | Ap2n | 2n | NA | rmMDS2 | GenBank | No | 0.00 | 617 | dataset5 | 0.09 | 0 | NA | 0 | Ap2n-kaz |
| KC193640 | Ap2n | 2n | NA | rmROC1 | GenBank | No | 0.00 | 617 | dataset5 | 0 | 0 | NA | 0 | Ap2n-kaz |
| KC193641 | Ap2n | 2n | NA | rmROC2 | GenBank | No | 0.00 | 617 | dataset5 | 0 | 0 | NA | 0 | Ap2n-kaz |
| KC193642 | Ap2n | 2n | NA | rmNOT1 | GenBank | No | 0.00 | 617 | dataset5 | 0 | 0 | NA | 0 | Ap2n-kaz |
| KC193643 | Ap2n | 2n | NA | rmNOT2 | GenBank | No | 0.00 | 617 | dataset5 | 0 | 0 | NA | 0 | Ap2n-kaz |
| KC193644 | Ap2n | 2n | NA | rmLAGK1 | GenBank | No | 0.00 | 617 | dataset5 | 0 | 0 | NA | 0 | Ap2n-kaz |
| KC193645 | Ap2n | 2n | NA | rmLAGK2 | GenBank | No | 0.00 | 617 | dataset5 | 0 | 0 | NA | 0 | Ap2n-kaz |
| KC193646 | Ap2n | 2n | NA | rmAIG1 | GenBank | No | 0.00 | 617 | dataset5 | 0 | 0 | NA | 0 | Ap2n-kaz |
| KC193647 | Ap2n | 2n | NA | rmAIG2 | GenBank | No | 0.00 | 617 | dataset5 | 0 | 0 | NA | 0 | Ap2n-kaz |
| KC193648 | Ap2n | 2n | NA | rmKOY1 | GenBank | No | 0.00 | 617 | dataset5 | 0 | 0 | NA | 0 | Ap2n-kaz |
| KC193649 | Ap2n | 2n | NA | rmKOY2 | GenBank | No | 0.00 | 617 | dataset5 | 0 | 0 | NA | 0 | Ap2n-kaz |
| KC193650 | Ap2n | 2n | NA | rmATA2 | GenBank | No | 0.00 | 617 | dataset5 | 0 | 0 | NA | 0 | Ap2n-kaz |
| KC193651 | Ap2n | 2n | NA | rmIRAQ1 | GenBank | No | 0.00 | 617 | dataset5 | 0 | 0 | NA | 0 | Ap2n-kaz |
| KC193652 | Ap2n | 2n | NA | rmIRAQ2 | GenBank | No | 0.00 | 617 | dataset5 | 0 | 0 | NA | 0 | Ap2n-kaz |
| KC193653 | Ap2n | 2n | NA | rmURM1 | GenBank | No | 0.00 | 617 | dataset5 | 0 | 0 | NA | 0 | Ap2n-kaz |
| KC193654 | Ap2n | 2n | NA | rmURM2 | GenBank | No | 0.00 | 617 | dataset5 | 0 | 0 | NA | 0 | Ap2n-kaz |
| KC193655 | Ap2n | 2n | NA | rmAIBI1 | GenBank | No | 0.00 | 617 | dataset5 | 0 | 0 | 1 | 0 | Ap2n-kaz |
| KC193656 | Ap2n | 2n | NA | rmAIBI2 | GenBank | No | 0.00 | 617 | dataset5 | 0 | 0 | NA | 0 | Ap2n-kaz |
| KC193657 | Ap2n | 2n | NA | rmXIAO1 | GenBank | No | 0.00 | 617 | dataset5 | 0 | 0 | 2 | 1 | Ap2n |
| KC193658 | Ap2n | 2n | NA | rmXIAO2 | GenBank | No | 0.00 | 617 | dataset5 | 0 | 0 | NA | 1 | Ap2n |
| KC193659 | Ap2n | 2n | NA | rmPAK1 | GenBank | No | 0.00 | 617 | dataset5 | 0 | 0 | 1 | 0 | Ap2n-kaz |
| KC193660 | Ap2n | 2n | NA | rmPAK2 | GenBank | No | 0.00 | 617 | dataset5 | 0 | 0 | NA | 0 | Ap2n-kaz |
| KC193661 | Ap2n | 2n | NA | rmMATA1 | GenBank | No | 0.00 | 617 | dataset5 | 0 | 0 | 3 | 1 | Ap2n |
| KC193662 | Ap2n | 2n | NA | rmMATA2 | GenBank | No | 0.00 | 617 | dataset5 | 0 | 0 | NA | 1 | Ap2n |
| KC193663 | Ap2n | 2n | NA | rmATA1 | GenBank | No | 0.00 | 617 | dataset5 | 0.06 | 0 | NA | 0 | Ap2n-urm |
| KC193664 | Ap2n | 2n | NA | rmKUJ1 | GenBank | No | 0.00 | 617 | dataset5 | 0 | 0 | 5 | 1 | Ap2n |
| KC193665 | Ap2n | 2n | NA | rmKUJ2 | GenBank | No | 0.00 | 617 | dataset5 | 0 | 0 | NA | 1 | Ap2n |
| KC193666 | Ap2n | 2n | NA | IRAQ2 | GenBank | No | 0.00 | 617 | dataset5 | 0 | 0 | NA | 0 | Ap2n-kaz |
| KC193668 | Ap2n | 2n | NA | LAGK1 | GenBank | No | 0.00 | 617 | dataset5 | 0 | 0 | NA | 0 | Ap2n-kaz |
| KC193670 | Ap2n | 2n | NA | AIG1 | GenBank | No | 0.00 | 617 | dataset5 | 0 | 0 | NA | 0 | Ap2n-kaz |
| KC193672 | Ap2n | 2n | NA | AIBI1 | GenBank | No | 0.00 | 617 | dataset5 | 0 | 0 | NA | 0 | Ap2n-kaz |
| KC193673 | Ap2n | 2n | NA | AIBI3 | GenBank | No | 0.00 | 617 | dataset5 | 0 | 0 | NA | 1 | Ap2n |
| KC193675 | Ap2n | 2n | NA | AIBI7 | GenBank | No | 0.00 | 617 | dataset5 | 0 | 0 | NA | 0 | Ap2n-urm |
| KC193676 | Ap2n | 2n | NA | LAGK4 | GenBank | No | 0.00 | 617 | dataset5 | 1.36 | 1 | NA | 1 | Ap2n |
| KC193677 | Ap2n | 2n | NA | MATA1 | GenBank | No | 0.00 | 617 | dataset5 | 0.06 | 0 | NA | 0 | Ap2n-urm |
| KU053797 | Ap2n | 2n | NA | DJA1 | GenBank | No | 0.00 | 625 | dataset5 | 0 | 0 | NA | 0 | Ap2n-urm |
| KU053798 | Ap2n | 2n | NA | DJA2 | GenBank | No | 0.00 | 625 | dataset5 | 0 | 0 | NA | 0 | Ap2n-urm |
| KU053799 | Ap2n | 2n | NA | DJA3 | GenBank | No | 0.00 | 625 | dataset5 | 0 | 0 | NA | 0 | Ap2n-urm |
| KU053800 | Ap2n | 2n | NA | DJA4 | GenBank | No | 0.00 | 625 | dataset5 | 0 | 0 | NA | 0 | Ap2n-urm |
| KU053801 | Ap2n | 2n | NA | DJA5 | GenBank | No | 0.00 | 625 | dataset5 | 0 | 0 | NA | 0 | Ap2n-urm |
| KU053802 | Ap2n | 2n | NA | DJA6 | GenBank | No | 0.00 | 625 | dataset5 | 0 | 0 | NA | 0 | Ap2n-urm |
| KU053803 | Ap2n | 2n | NA | SAK1 | GenBank | No | 0.00 | 625 | dataset5 | 0 | 0 | NA | 0 | Ap2n-urm |
| KU053804 | Ap2n | 2n | NA | SAK2 | GenBank | No | 0.00 | 625 | dataset5 | 0 | 0 | NA | 0 | Ap2n-urm |
| KU053805 | Ap2n | 2n | NA | SAK3 | GenBank | No | 0.00 | 625 | dataset5 | 0 | 0 | NA | 0 | Ap2n-urm |
| KU053806 | Ap2n | 2n | NA | SAK4 | GenBank | No | 0.00 | 625 | dataset5 | 0 | 0 | NA | 0 | Ap2n-urm |
| KU053807 | Ap2n | 2n | NA | SAK5 | GenBank | No | 0.00 | 625 | dataset5 | 0 | 0 | NA | 0 | Ap2n-urm |
| KU053808 | Ap2n | 2n | NA | DZH1 | GenBank | No | 0.00 | 625 | dataset5 | 0 | 0 | NA | 0 | Ap2n-urm |
| KU053809 | Ap2n | 2n | NA | DZH2 | GenBank | No | 0.00 | 625 | dataset5 | 0 | 0 | NA | 0 | Ap2n-urm |
| KU053810 | Ap2n | 2n | NA | DZH3 | GenBank | No | 0.00 | 625 | dataset5 | 0 | 0 | NA | 0 | Ap2n-urm |
| KU053811 | Ap2n | 2n | NA | DZH4 | GenBank | No | 0.00 | 625 | dataset5 | 0 | 0 | 1 | 0 | Ap2n-urm |
| KU053812 | Ap2n | 2n | NA | DZH5 | GenBank | No | 0.00 | 625 | dataset5 | 0 | 0 | NA | 0 | Ap2n-urm |
| KU053813 | Ap2n | 2n | NA | DZH6 | GenBank | No | 0.00 | 625 | dataset5 | 0 | 0 | NA | 0 | Ap2n-urm |
| KU053814 | Ap2n | 2n | NA | DZH7 | GenBank | No | 0.00 | 625 | dataset5 | 0 | 0 | NA | 0 | Ap2n-urm |
| KU053815 | Ap2n | 2n | NA | DZH8 | GenBank | No | 0.00 | 625 | dataset5 | 0 | 0 | NA | 0 | Ap2n-urm |
| KU053816 | Ap2n | 2n | NA | DZH9 | GenBank | No | 0.00 | 625 | dataset5 | 0 | 0 | NA | 0 | Ap2n-urm |
| KU053817 | Ap2n | 2n | NA | DZH10 | GenBank | No | 0.00 | 625 | dataset5 | 0 | 0 | NA | 0 | Ap2n-urm |
| KU053818 | Ap2n | 2n | NA | DZH11 | GenBank | No | 0.00 | 625 | dataset5 | 0 | 0 | NA | 0 | Ap2n-urm |
| KU053819 | Ap2n | 2n | NA | DZH12 | GenBank | No | 0.00 | 625 | dataset5 | 0 | 0 | 1 | 0 | Ap2n-urm |
| KF691148 | Ap2n | 2n | NA | BAM1 | GenBank | No | 0.00 | 561 | dataset5 | 0 | 0 | NA | 0 | Ap2n-kaz |
| KF691149 | Ap2n | 2n | NA | BAM2 | GenBank | No | 0.00 | 561 | dataset5 | 0 | 0 | NA | 0 | Ap2n-kaz |
| KF691150 | Ap2n | 2n | NA | BAM3 | GenBank | No | 0.00 | 561 | dataset5 | 0 | 0 | NA | 0 | Ap2n-kaz |
| KF691151 | Ap2n | 2n | NA | BAM4 | GenBank | No | 0.00 | 561 | dataset5 | 0 | 0 | NA | 0 | Ap2n-kaz |
| KF691152 | Ap2n | 2n | NA | BAM5 | GenBank | No | 0.00 | 561 | dataset5 | 0 | 0 | NA | 0 | Ap2n-kaz |
| KF691153 | Ap2n | 2n | NA | BAM6 | GenBank | No | 0.00 | 561 | dataset5 | 0 | 0 | NA | 0 | Ap2n-kaz |
| KF691166 | Ap2n | 2n | NA | CAN1 | GenBank | No | 0.00 | 561 | dataset5 | 0 | 0 | NA | 0 | Ap2n-kaz |
| KF691167 | Ap2n | 2n | NA | CAN2 | GenBank | No | 0.00 | 561 | dataset5 | 0 | 0 | NA | 0 | Ap2n-kaz |
| KF691168 | Ap2n | 2n | NA | CAN3 | GenBank | No | 0.00 | 561 | dataset5 | 0 | 0 | NA | 0 | Ap2n-kaz |
| KF691169 | Ap2n | 2n | NA | CAN4 | GenBank | No | 0.00 | 561 | dataset5 | 0 | 0 | NA | 0 | Ap2n-kaz |
| KF691170 | Ap2n | 2n | NA | CHE1 | GenBank | No | 0.00 | 561 | dataset5 | 0 | 0 | NA | 0 | Ap2n-kaz |
| KF691171 | Ap2n | 2n | NA | CHE2 | GenBank | No | 0.00 | 561 | dataset5 | 0 | 0 | NA | 0 | Ap2n-kaz |
| KF691172 | Ap2n | 2n | NA | CHE3 | GenBank | No | 0.00 | 561 | dataset5 | 0 | 0 | NA | 0 | Ap2n-kaz |
| KF691183 | Ap2n | 2n | NA | DLI8 | GenBank | No | 0.00 | 561 | dataset5 | 0 | 0 | NA | 0 | Ap2n-kaz |
| KF691187 | Ap2n | 2n | NA | DON1 | GenBank | No | 0.00 | 561 | dataset5 | 0 | 0 | NA | 0 | Ap2n-kaz |
| KF691188 | Ap2n | 2n | NA | DON2 | GenBank | No | 0.00 | 561 | dataset5 | 0 | 0 | NA | 0 | Ap2n-kaz |
| KF691189 | Ap2n | 2n | NA | DON3 | GenBank | No | 0.00 | 561 | dataset5 | 0 | 0 | NA | 0 | Ap2n-kaz |
| KF691199 | Ap2n | 2n | NA | GAH1.2 | GenBank | No | 0.00 | 561 | dataset5 | 0 | 0 | NA | 0 | Ap2n-kaz |
| KF691200 | Ap2n | 2n | NA | GAH2 | GenBank | No | 0.00 | 561 | dataset5 | 0 | 0 | NA | 0 | Ap2n-kaz |
| KF691201 | Ap2n | 2n | NA | GAH3.2 | GenBank | No | 0.00 | 561 | dataset5 | 0 | 0 | NA | 0 | Ap2n-kaz |
| KF691202 | Ap2n | 2n | NA | GAH4.2 | GenBank | No | 0.00 | 561 | dataset5 | 0 | 0 | NA | 0 | Ap2n-kaz |
| KF691203 | Ap2n | 2n | NA | GAH5.2 | GenBank | No | 0.00 | 561 | dataset5 | 0 | 0 | NA | 0 | Ap2n-kaz |
| KF691204 | Ap2n | 2n | NA | GAH6.2 | GenBank | No | 0.00 | 561 | dataset5 | 0 | 0 | NA | 0 | Ap2n-kaz |
| KF691208 | Ap2n | 2n | NA | HAN1 | GenBank | No | 0.00 | 561 | dataset5 | 0 | 0 | NA | 0 | Ap2n-kaz |
| KF691209 | Ap2n | 2n | NA | HAN2 | GenBank | No | 0.00 | 561 | dataset5 | 0 | 0 | NA | 0 | Ap2n-kaz |
| KF691210 | Ap2n | 2n | NA | HAN3 | GenBank | No | 0.00 | 561 | dataset5 | 0 | 0 | NA | 0 | Ap2n-kaz |
| KF691211 | Ap2n | 2n | NA | HAN4 | GenBank | No | 0.00 | 561 | dataset5 | 0 | 0 | NA | 0 | Ap2n-kaz |
| KF691212 | Ap2n | 2n | NA | HAN5 | GenBank | No | 0.00 | 561 | dataset5 | 0 | 0 | NA | 0 | Ap2n-kaz |
| KF691213 | Ap2n | 2n | NA | HAN6 | GenBank | No | 0.00 | 561 | dataset5 | 0 | 0 | 1 | 0 | Ap2n-kaz |
| KF691214 | Ap2n | 2n | NA | HAN7 | GenBank | No | 0.00 | 561 | dataset5 | 0 | 0 | NA | 0 | Ap2n-kaz |
| KF691224 | Ap2n | 2n | NA | LUA3 | GenBank | No | 0.00 | 561 | dataset5 | 0 | 0 | NA | 0 | Ap2n-kaz |
| KF691225 | Ap2n | 2n | NA | LUA4 | GenBank | No | 0.00 | 561 | dataset5 | 0 | 0 | NA | 0 | Ap2n-kaz |
| KF691226 | Ap2n | 2n | NA | LUA5 | GenBank | No | 0.00 | 561 | dataset5 | 0 | 0 | NA | 0 | Ap2n-kaz |
| KF691233 | Ap2n | 2n | NA | SHA1 | GenBank | No | 0.00 | 561 | dataset5 | 0 | 0 | NA | 0 | Ap2n-kaz |
| KF691234 | Ap2n | 2n | NA | SHA2 | GenBank | No | 0.00 | 561 | dataset5 | 0 | 0 | NA | 0 | Ap2n-kaz |
| KF691235 | Ap2n | 2n | NA | SHA3 | GenBank | No | 0.00 | 561 | dataset5 | 0 | 0 | NA | 0 | Ap2n-kaz |
| KF691236 | Ap2n | 2n | NA | SID1 | GenBank | No | 0.00 | 561 | dataset5 | 0.09 | 0 | NA | 0 | Ap2n-kaz |
| KF691238 | Ap2n | 2n | NA | SID3 | GenBank | No | 0.00 | 561 | dataset5 | 0.09 | 0 | NA | 0 | Ap2n-kaz |
| KF691265 | Ap2n | 2n | NA | XIA1 | GenBank | No | 0.00 | 561 | dataset5 | 0 | 0 | NA | 0 | Ap2n-kaz |
| KF691268 | Ap2n | 2n | NA | XIA4 | GenBank | No | 0.00 | 561 | dataset5 | 0 | 0 | NA | 0 | Ap2n-kaz |
| KF691287 | Ap2n | 2n | NA | YIN1 | GenBank | No | 0.00 | 561 | dataset5 | 0 | 0 | NA | 0 | Ap2n-kaz |
| KF691288 | Ap2n | 2n | NA | YIN2 | GenBank | No | 0.00 | 561 | dataset5 | 0 | 0 | NA | 0 | Ap2n-kaz |
| KF691289 | Ap2n | 2n | NA | YIN3 | GenBank | No | 0.00 | 561 | dataset5 | 0 | 0 | NA | 0 | Ap2n-kaz |
| KF691290 | Ap2n | 2n | NA | YIN4 | GenBank | No | 0.00 | 561 | dataset5 | 0 | 0 | NA | 0 | Ap2n-kaz |
| KF691333 | Ap2n | 2n | NA | INC1 | GenBank | No | 0.00 | 561 | dataset5 | 0 | 0 | NA | 0 | Ap2n-kaz |
| KF691334 | Ap2n | 2n | NA | INC2 | GenBank | No | 0.00 | 561 | dataset5 | 0 | 0 | NA | 0 | Ap2n-kaz |
| KF691335 | Ap2n | 2n | NA | INC3 | GenBank | No | 0.00 | 561 | dataset5 | 0 | 0 | NA | 0 | Ap2n-kaz |
| KF691336 | Ap2n | 2n | NA | INC4 | GenBank | No | 0.00 | 561 | dataset5 | 0 | 0 | NA | 0 | Ap2n-kaz |
| KF691337 | Ap2n | 2n | NA | INC5 | GenBank | No | 0.00 | 561 | dataset5 | 0 | 0 | NA | 0 | Ap2n-kaz |
| KF691338 | Ap2n | 2n | NA | LAGW1 | GenBank | No | 0.00 | 561 | dataset5 | 0 | 0 | NA | 0 | Ap2n-kaz |
| KF691339 | Ap2n | 2n | NA | LAGW2 | GenBank | No | 0.00 | 561 | dataset5 | 0 | 0 | NA | 0 | Ap2n-kaz |
| KF691340 | Ap2n | 2n | NA | LAGW3 | GenBank | No | 0.00 | 561 | dataset5 | 0 | 0 | NA | 0 | Ap2n-kaz |
| KF691341 | Ap2n | 2n | NA | LAGW4 | GenBank | No | 0.00 | 561 | dataset5 | 0 | 0 | NA | 0 | Ap2n-kaz |
| KF691342 | Ap2n | 2n | NA | LAGW5 | GenBank | No | 0.00 | 561 | dataset5 | 0 | 0 | NA | 0 | Ap2n-kaz |
| KF691343 | Ap2n | 2n | NA | LAGE1 | GenBank | No | 0.00 | 561 | dataset5 | 0 | 0 | NA | 0 | Ap2n-kaz |
| KF691344 | Ap2n | 2n | NA | LAGE2 | GenBank | No | 0.00 | 561 | dataset5 | 0 | 0 | NA | 0 | Ap2n-kaz |
| KF691345 | Ap2n | 2n | NA | LAGE3 | GenBank | No | 0.00 | 561 | dataset5 | 0 | 0 | NA | 0 | Ap2n-kaz |
| KF691346 | Ap2n | 2n | NA | MAHR1 | GenBank | No | 0.00 | 561 | dataset5 | 0 | 0 | NA | 0 | Ap2n-kaz |
| KF691348 | Ap2n | 2n | NA | MAHR3 | GenBank | No | 0.00 | 561 | dataset5 | 0 | 0 | NA | 0 | Ap2n-kaz |
| KF691357 | Ap2n | 2n | NA | MIG1 | GenBank | No | 0.00 | 561 | dataset5 | 0 | 0 | NA | 0 | Ap2n-kaz |
| KF691358 | Ap2n | 2n | NA | MIG2 | GenBank | No | 0.00 | 561 | dataset5 | 0 | 0 | NA | 0 | Ap2n-kaz |
| KF691359 | Ap2n | 2n | NA | MIG3 | GenBank | No | 0.00 | 561 | dataset5 | 0 | 0 | NA | 0 | Ap2n-kaz |
| KF691360 | Ap2n | 2n | NA | MIG4 | GenBank | No | 0.00 | 561 | dataset5 | 0 | 0 | 1 | 0 | Ap2n-kaz |
| KF691361 | Ap2n | 2n | NA | MIG5 | GenBank | No | 0.00 | 561 | dataset5 | 0 | 0 | NA | 0 | Ap2n-kaz |
| KF691367 | Ap2n | 2n | NA | QOM1 | GenBank | No | 0.00 | 561 | dataset5 | 0 | 0 | NA | 0 | Ap2n-kaz |
| KF691368 | Ap2n | 2n | NA | QOM2 | GenBank | No | 0.00 | 561 | dataset5 | 0 | 0 | NA | 0 | Ap2n-kaz |
| KF691369 | Ap2n | 2n | NA | QOM3 | GenBank | No | 0.00 | 561 | dataset5 | 0 | 0 | NA | 0 | Ap2n-kaz |
| KF691370 | Ap2n | 2n | NA | QOM4 | GenBank | No | 0.00 | 561 | dataset5 | 0 | 0 | NA | 0 | Ap2n-kaz |
| KF691371 | Ap2n | 2n | NA | QOM5 | GenBank | No | 0.00 | 561 | dataset5 | 0 | 0 | NA | 0 | Ap2n-kaz |
| KF691372 | Ap2n | 2n | NA | QOM6 | GenBank | No | 0.00 | 561 | dataset5 | 0 | 0 | NA | 0 | Ap2n-kaz |
| KF691373 | Ap2n | 2n | NA | ABG1 | GenBank | No | 0.00 | 561 | dataset5 | 0 | 0 | NA | 0 | Ap2n-kaz |
| KF691374 | Ap2n | 2n | NA | ABG2 | GenBank | No | 0.00 | 561 | dataset5 | 0 | 0 | NA | 0 | Ap2n-kaz |
| KF691375 | Ap2n | 2n | NA | ABG3 | GenBank | No | 0.00 | 561 | dataset5 | 0 | 0 | NA | 0 | Ap2n-kaz |
| KF691391 | Ap2n | 2n | NA | ARS1 | GenBank | No | 0.00 | 561 | dataset5 | 0 | 0 | NA | 0 | Ap2n-kaz |
| KF691392 | Ap2n | 2n | NA | ARS2 | GenBank | No | 0.00 | 561 | dataset5 | 0 | 0 | NA | 0 | Ap2n-kaz |
| KF691393 | Ap2n | 2n | NA | ARS3 | GenBank | No | 0.00 | 561 | dataset5 | 0 | 0 | NA | 0 | Ap2n-kaz |
| KF691394 | Ap2n | 2n | NA | ARS4 | GenBank | No | 0.00 | 561 | dataset5 | 0 | 0 | NA | 0 | Ap2n-kaz |
| KF691395 | Ap2n | 2n | NA | ARS5 | GenBank | No | 0.00 | 561 | dataset5 | 0 | 0 | NA | 0 | Ap2n-kaz |
| KF691396 | Ap2n | 2n | NA | ARS6 | GenBank | No | 0.00 | 561 | dataset5 | 0 | 0 | NA | 0 | Ap2n-kaz |
| KF691397 | Ap2n | 2n | NA | ARS7 | GenBank | No | 0.00 | 561 | dataset5 | 0 | 0 | NA | 0 | Ap2n-kaz |
| KF691398 | Ap2n | 2n | NA | ASS1 | GenBank | No | 0.00 | 561 | dataset5 | 0.09 | 0 | NA | 0 | Ap2n-kaz |
| KF691399 | Ap2n | 2n | NA | ASS2 | GenBank | No | 0.00 | 561 | dataset5 | 0.09 | 0 | NA | 0 | Ap2n-kaz |
| KF691400 | Ap2n | 2n | NA | ASS3 | GenBank | No | 0.00 | 561 | dataset5 | 0 | 0 | NA | 0 | Ap2n-kaz |
| KF691401 | Ap2n | 2n | NA | ASS4 | GenBank | No | 0.00 | 561 | dataset5 | 0.09 | 0 | NA | 0 | Ap2n-kaz |
| KF691402 | Ap2n | 2n | NA | ASS5 | GenBank | No | 0.00 | 561 | dataset5 | 0.09 | 0 | NA | 0 | Ap2n-kaz |
| KF691403 | Ap2n | 2n | NA | ASS6 | GenBank | No | 0.00 | 561 | dataset5 | 0.09 | 0 | NA | 0 | Ap2n-kaz |
| KF691404 | Ap2n | 2n | NA | KYZ1 | GenBank | No | 0.00 | 561 | dataset5 | 0 | 0 | NA | 0 | Ap2n-kaz |
| KF691405 | Ap2n | 2n | NA | KYZ2 | GenBank | No | 0.00 | 561 | dataset5 | 0 | 0 | NA | 0 | Ap2n-kaz |
| KF691406 | Ap2n | 2n | NA | KYZ3 | GenBank | No | 0.00 | 561 | dataset5 | 0 | 0 | NA | 0 | Ap2n-kaz |
| KF691407 | Ap2n | 2n | NA | KYZ4 | GenBank | No | 0.00 | 561 | dataset5 | 0 | 0 | NA | 0 | Ap2n-kaz |
| KF691408 | Ap2n | 2n | NA | KYZ5 | GenBank | No | 0.00 | 561 | dataset5 | 0 | 0 | NA | 0 | Ap2n-kaz |
| KF691409 | Ap2n | 2n | NA | NCS1 | GenBank | No | 0.00 | 561 | dataset5 | 0 | 0 | NA | 0 | Ap2n-kaz |
| KF691410 | Ap2n | 2n | NA | NCS2 | GenBank | No | 0.00 | 561 | dataset5 | 0 | 0 | NA | 0 | Ap2n-kaz |
| KF691411 | Ap2n | 2n | NA | NCS3 | GenBank | No | 0.00 | 561 | dataset5 | 0 | 0 | NA | 0 | Ap2n-kaz |
| KF691412 | Ap2n | 2n | NA | NCS4 | GenBank | No | 0.00 | 561 | dataset5 | 0 | 0 | NA | 0 | Ap2n-kaz |
| KF691413 | Ap2n | 2n | NA | NCS5 | GenBank | No | 0.00 | 561 | dataset5 | 0 | 0 | NA | 0 | Ap2n-kaz |
| KF691414 | Ap2n | 2n | NA | NCS6 | GenBank | No | 0.00 | 561 | dataset5 | 0 | 0 | NA | 0 | Ap2n-kaz |
| KF691415 | Ap2n | 2n | NA | PAV1 | GenBank | No | 0.00 | 561 | dataset5 | 0 | 0 | NA | 0 | Ap2n-kaz |
| KF691416 | Ap2n | 2n | NA | PAV2 | GenBank | No | 0.00 | 561 | dataset5 | 0 | 0 | NA | 0 | Ap2n-kaz |
| KF691417 | Ap2n | 2n | NA | PAV3 | GenBank | No | 0.00 | 561 | dataset5 | 0 | 0 | NA | 0 | Ap2n-kaz |
| KF691418 | Ap2n | 2n | NA | PAV4 | GenBank | No | 0.00 | 561 | dataset5 | 0 | 0 | NA | 0 | Ap2n-kaz |
| KF691419 | Ap2n | 2n | NA | PAV5 | GenBank | No | 0.00 | 561 | dataset5 | 0 | 0 | NA | 0 | Ap2n-kaz |
| KF691420 | Ap2n | 2n | NA | PAV6 | GenBank | No | 0.00 | 561 | dataset5 | 0 | 0 | NA | 0 | Ap2n-kaz |
| KF691421 | Ap2n | 2n | NA | TUZ1 | GenBank | No | 0.00 | 561 | dataset5 | 0 | 0 | NA | 0 | Ap2n-kaz |
| KF691422 | Ap2n | 2n | NA | TUZ2 | GenBank | No | 0.00 | 561 | dataset5 | 0 | 0 | NA | 0 | Ap2n-kaz |
| KF691423 | Ap2n | 2n | NA | TUZ3 | GenBank | No | 0.00 | 561 | dataset5 | 0 | 0 | NA | 0 | Ap2n-kaz |
| KF691424 | Ap2n | 2n | NA | TUZ4 | GenBank | No | 0.00 | 561 | dataset5 | 0 | 0 | NA | 0 | Ap2n-kaz |
| KF691425 | Ap2n | 2n | NA | TUZ5 | GenBank | No | 0.00 | 561 | dataset5 | 0 | 0 | NA | 0 | Ap2n-kaz |
| KF691426 | Ap2n | 2n | NA | TUZ6 | GenBank | No | 0.00 | 561 | dataset5 | 0 | 0 | NA | 0 | Ap2n-kaz |
| KF691427 | Ap2n | 2n | NA | TUZ7 | GenBank | No | 0.00 | 561 | dataset5 | 0 | 0 | NA | 0 | Ap2n-kaz |
| KF691428 | Ap2n | 2n | NA | TUZ8 | GenBank | No | 0.00 | 561 | dataset5 | 0 | 0 | NA | 0 | Ap2n-kaz |
| KF691429 | Ap2n | 2n | NA | TUZ9 | GenBank | No | 0.00 | 561 | dataset5 | 0 | 0 | NA | 0 | Ap2n-kaz |
| KF691430 | Ap2n | 2n | NA | TUZ10 | GenBank | No | 0.00 | 561 | dataset5 | 0 | 0 | NA | 0 | Ap2n-kaz |
| KF691431 | Ap2n | 2n | NA | TUZ11 | GenBank | No | 0.00 | 561 | dataset5 | 0 | 0 | NA | 0 | Ap2n-kaz |
| KF691432 | Ap2n | 2n | NA | TUZ12 | GenBank | No | 0.00 | 561 | dataset5 | 0 | 0 | NA | 0 | Ap2n-kaz |
| KF691433 | Ap2n | 2n | NA | TUZ13 | GenBank | No | 0.00 | 561 | dataset5 | 0 | 0 | NA | 0 | Ap2n-kaz |
| KF691434 | Ap2n | 2n | NA | TUZ14 | GenBank | No | 0.00 | 561 | dataset5 | 0 | 0 | NA | 0 | Ap2n-kaz |
| KF691442 | Ap2n | 2n | NA | KOC4 | GenBank | No | 0.00 | 561 | dataset5 | 0 | 0 | NA | 0 | Ap2n-kaz |
| KF691443 | Ap2n | 2n | NA | KOC5 | GenBank | No | 0.00 | 561 | dataset5 | 0 | 0 | NA | 0 | Ap2n-kaz |
| KF691444 | Ap2n | 2n | NA | KOC6 | GenBank | No | 0.00 | 561 | dataset5 | 0 | 0 | NA | 0 | Ap2n-kaz |
| KF691445 | Ap2n | 2n | NA | KOC7 | GenBank | No | 0.00 | 561 | dataset5 | 0 | 0 | NA | 0 | Ap2n-kaz |
| KF691447 | Ap2n | 2n | NA | KOC9 | GenBank | No | 0.00 | 561 | dataset5 | 0 | 0 | NA | 0 | Ap2n-kaz |
| KF691448 | Ap2n | 2n | NA | KOC10 | GenBank | No | 0.00 | 561 | dataset5 | 0 | 0 | NA | 0 | Ap2n-kaz |
| KF691455 | Ap2n | 2n | NA | BYA1 | GenBank | No | 0.00 | 561 | dataset5 | 0 | 0 | NA | 0 | Ap2n-kaz |
| KF691456 | Ap2n | 2n | NA | BYA2 | GenBank | No | 0.00 | 561 | dataset5 | 0 | 0 | NA | 0 | Ap2n-kaz |
| KF691457 | Ap2n | 2n | NA | BYA3 | GenBank | No | 0.00 | 561 | dataset5 | 0 | 0 | NA | 0 | Ap2n-kaz |
| KF691458 | Ap2n | 2n | NA | BYA4 | GenBank | No | 0.00 | 561 | dataset5 | 0 | 0 | NA | 0 | Ap2n-kaz |
| KF691459 | Ap2n | 2n | NA | BYA5 | GenBank | No | 0.00 | 561 | dataset5 | 0 | 0 | NA | 0 | Ap2n-kaz |
| KF691460 | Ap2n | 2n | NA | EBE1 | GenBank | No | 0.00 | 561 | dataset5 | 0 | 0 | NA | 0 | Ap2n-kaz |
| KF691461 | Ap2n | 2n | NA | EBE2 | GenBank | No | 0.00 | 561 | dataset5 | 0 | 0 | NA | 0 | Ap2n-kaz |
| KF691462 | Ap2n | 2n | NA | EBE3 | GenBank | No | 0.00 | 561 | dataset5 | 0 | 0 | NA | 0 | Ap2n-kaz |
| KF691463 | Ap2n | 2n | NA | EBE4 | GenBank | No | 0.00 | 561 | dataset5 | 0 | 0 | NA | 0 | Ap2n-kaz |
| KF691464 | Ap2n | 2n | NA | EBE5 | GenBank | No | 0.00 | 561 | dataset5 | 0 | 0 | NA | 0 | Ap2n-kaz |
| KF691465 | Ap2n | 2n | NA | EBE6 | GenBank | No | 0.00 | 561 | dataset5 | 0 | 0 | NA | 0 | Ap2n-kaz |
| KF691466 | Ap2n | 2n | NA | EBE7 | GenBank | No | 0.00 | 561 | dataset5 | 0 | 0 | NA | 0 | Ap2n-kaz |
| KF691467 | Ap2n | 2n | NA | GOR1 | GenBank | No | 0.00 | 561 | dataset5 | 0 | 0 | NA | 0 | Ap2n-kaz |
| KF691468 | Ap2n | 2n | NA | GOR2 | GenBank | No | 0.00 | 561 | dataset5 | 0 | 0 | NA | 0 | Ap2n-kaz |
| KF691469 | Ap2n | 2n | NA | GOR3 | GenBank | No | 0.00 | 561 | dataset5 | 0 | 0 | NA | 0 | Ap2n-kaz |
| KF691470 | Ap2n | 2n | NA | GOR4 | GenBank | No | 0.00 | 561 | dataset5 | 0 | 0 | NA | 0 | Ap2n-kaz |
| KF691471 | Ap2n | 2n | NA | GOR5 | GenBank | No | 0.00 | 561 | dataset5 | 0 | 0 | NA | 0 | Ap2n-kaz |
| KF691472 | Ap2n | 2n | NA | KUC1 | GenBank | No | 0.00 | 561 | dataset5 | 0 | 0 | NA | 0 | Ap2n-kaz |
| KF691473 | Ap2n | 2n | NA | KUC2 | GenBank | No | 0.00 | 561 | dataset5 | 0 | 0 | NA | 0 | Ap2n-kaz |
| KF691474 | Ap2n | 2n | NA | KUC3 | GenBank | No | 0.00 | 561 | dataset5 | 0 | 0 | NA | 0 | Ap2n-kaz |
| KF691475 | Ap2n | 2n | NA | KUL1 | GenBank | No | 0.00 | 561 | dataset5 | 0 | 0 | NA | 0 | Ap2n-kaz |
| KF691476 | Ap2n | 2n | NA | KUL2 | GenBank | No | 0.00 | 561 | dataset5 | 0 | 0 | NA | 0 | Ap2n-kaz |
| KF691477 | Ap2n | 2n | NA | KUL3 | GenBank | No | 0.00 | 561 | dataset5 | 0 | 0 | NA | 0 | Ap2n-kaz |
| KF691478 | Ap2n | 2n | NA | KUR1 | GenBank | No | 0.00 | 561 | dataset5 | 0 | 0 | NA | 0 | Ap2n-kaz |
| KF691479 | Ap2n | 2n | NA | KUR2 | GenBank | No | 0.00 | 561 | dataset5 | 0 | 0 | 1 | 0 | Ap2n-kaz |
| KF691480 | Ap2n | 2n | NA | KUR3 | GenBank | No | 0.00 | 561 | dataset5 | 0 | 0 | NA | 0 | Ap2n-kaz |
| KF691481 | Ap2n | 2n | NA | MME1 | GenBank | No | 0.00 | 561 | dataset5 | 0.16 | 1 | 0 | 1 | Ap2n |
| KF691482 | Ap2n | 2n | NA | MME2 | GenBank | No | 0.00 | 561 | dataset5 | 0 | 0 | NA | 0 | Ap2n-kaz |
| KF691483 | Ap2n | 2n | NA | MME3 | GenBank | No | 0.00 | 561 | dataset5 | 0 | 0 | 1 | 0 | Ap2n-kaz |
| KF691484 | Ap2n | 2n | NA | MME4 | GenBank | No | 0.00 | 561 | dataset5 | 0 | 0 | NA | 0 | Ap2n-kaz |
| KF691485 | Ap2n | 2n | NA | MYA1 | GenBank | No | 0.00 | 561 | dataset5 | 0 | 0 | NA | 0 | Ap2n-kaz |
| KF691486 | Ap2n | 2n | NA | MYA2 | GenBank | No | 0.00 | 561 | dataset5 | 0 | 0 | NA | 0 | Ap2n-kaz |
| KF691487 | Ap2n | 2n | NA | MYA3 | GenBank | No | 0.00 | 561 | dataset5 | 0 | 0 | NA | 0 | Ap2n-kaz |
| KF691488 | Ap2n | 2n | NA | MYA4 | GenBank | No | 0.00 | 561 | dataset5 | 0 | 0 | NA | 0 | Ap2n-kaz |
| KF691489 | Ap2n | 2n | NA | MYA5 | GenBank | No | 0.00 | 561 | dataset5 | 0 | 0 | NA | 0 | Ap2n-kaz |
| KF691490 | Ap2n | 2n | NA | MYA6 | GenBank | No | 0.00 | 561 | dataset5 | 0 | 0 | NA | 0 | Ap2n-kaz |
| KF691491 | Ap2n | 2n | NA | MYA7 | GenBank | No | 0.00 | 561 | dataset5 | 0 | 0 | NA | 0 | Ap2n-kaz |
| KF691492 | Ap2n | 2n | NA | MED1 | GenBank | No | 0.00 | 561 | dataset5 | 0 | 0 | NA | 0 | Ap2n-kaz |
| KF691493 | Ap2n | 2n | NA | MED2 | GenBank | No | 0.00 | 561 | dataset5 | 0 | 0 | NA | 0 | Ap2n-kaz |
| KF691494 | Ap2n | 2n | NA | MED3 | GenBank | No | 0.00 | 561 | dataset5 | 0 | 0 | NA | 0 | Ap2n-kaz |
| KF691495 | Ap2n | 2n | NA | VOS1 | GenBank | No | 0.00 | 561 | dataset5 | 0 | 0 | NA | 0 | Ap2n-kaz |
| KF691496 | Ap2n | 2n | NA | VOS2 | GenBank | No | 0.00 | 561 | dataset5 | 0 | 0 | NA | 0 | Ap2n-kaz |
| KF691497 | Ap2n | 2n | NA | VOS3 | GenBank | No | 0.00 | 561 | dataset5 | 0 | 0 | NA | 0 | Ap2n-kaz |
| KF691520 | Ap2n | 2n | NA | CAM1 | GenBank | No | 0.00 | 561 | dataset5 | 0.06 | 0 | NA | 0 | Ap2n-urm |
| KF691521 | Ap2n | 2n | NA | CAM2 | GenBank | No | 0.00 | 561 | dataset5 | 0 | 0 | NA | 0 | Ap2n-kaz |
| KF691522 | Ap2n | 2n | NA | CAM3 | GenBank | No | 0.00 | 561 | dataset5 | 0.06 | 0 | NA | 0 | Ap2n-urm |
| KF691523 | Ap2n | 2n | NA | CAM4 | GenBank | No | 0.00 | 561 | dataset5 | 0.06 | 0 | NA | 0 | Ap2n-urm |
| KF691524 | Ap2n | 2n | NA | CAM5 | GenBank | No | 0.00 | 561 | dataset5 | 0.06 | 0 | NA | 0 | Ap2n-urm |
| KF691525 | Ap2n | 2n | NA | CAM6 | GenBank | No | 0.00 | 561 | dataset5 | 0.06 | 0 | NA | 0 | Ap2n-urm |
| KF691526 | Ap2n | 2n | NA | CAM7 | GenBank | No | 0.00 | 561 | dataset5 | 0 | 0 | 6 | 1 | Ap2n |
| KF691527 | Ap2n | 2n | NA | CAM8 | GenBank | No | 0.00 | 561 | dataset5 | 0.06 | 0 | 2 | 1 | Ap2n |
| KF691528 | Ap2n | 2n | NA | CAM9 | GenBank | No | 0.00 | 561 | dataset5 | 0.06 | 0 | NA | 0 | Ap2n-urm |
| KF691529 | Ap2n | 2n | NA | CAM10 | GenBank | No | 0.00 | 561 | dataset5 | 0.06 | 0 | NA | 0 | Ap2n-urm |
| KF691530 | Ap2n | 2n | NA | KBG1 | GenBank | No | 0.00 | 561 | dataset5 | 0 | 0 | NA | 0 | Ap2n-kaz |
| KF691531 | Ap2n | 2n | NA | KBG2 | GenBank | No | 0.00 | 561 | dataset5 | 0 | 0 | NA | 0 | Ap2n-kaz |
| KF691532 | Ap2n | 2n | NA | KBG3 | GenBank | No | 0.00 | 561 | dataset5 | 0 | 0 | NA | 0 | Ap2n-kaz |
| KF691534 | Ap2n | 2n | NA | KBG5 | GenBank | No | 0.00 | 561 | dataset5 | 0 | 0 | NA | 0 | Ap2n-kaz |
| KF691547 | Ap2n | 2n | NA | CAA1 | GenBank | No | 0.00 | 561 | dataset5 | 0.04 | 0 | 0 | 0 | Ap2n-kaz |
| KF691548 | Ap2n | 2n | NA | CAA2 | GenBank | No | 0.00 | 561 | dataset5 | 0 | 0 | NA | 0 | Ap2n-kaz |
| KF691549 | Ap2n | 2n | NA | CAA3 | GenBank | No | 0.00 | 561 | dataset5 | 0 | 0 | NA | 0 | Ap2n-kaz |
| KF691550 | Ap2n | 2n | NA | CAA4 | GenBank | No | 0.00 | 561 | dataset5 | 0 | 0 | NA | 0 | Ap2n-kaz |
| KF691551 | Ap2n | 2n | NA | CAA5 | GenBank | No | 0.00 | 561 | dataset5 | 0 | 0 | NA | 0 | Ap2n-kaz |
| KF691552 | Ap2n | 2n | NA | CAA6 | GenBank | No | 0.00 | 561 | dataset5 | 0 | 0 | NA | 0 | Ap2n-kaz |
| KF691553 | Ap2n | 2n | NA | CAA7 | GenBank | No | 0.00 | 561 | dataset5 | 0 | 0 | NA | 0 | Ap2n-kaz |
| KF691554 | Ap2n | 2n | NA | CAA8 | GenBank | No | 0.00 | 561 | dataset5 | 0.04 | 0 | 0 | 0 | Ap2n-kaz |
| KF691555 | Ap2n | 2n | NA | CAA9 | GenBank | No | 0.00 | 561 | dataset5 | 0 | 0 | NA | 0 | Ap2n-kaz |
| KP090297 | Ap2n | 2n | NA | ELA1 | GenBank | No | 0.00 | 557 | dataset5 | 0 | 0 | NA | 0 | Ap2n-kaz |
| KP090298 | Ap2n | 2n | NA | ELA2 | GenBank | No | 0.00 | 557 | dataset5 | 0 | 0 | NA | 0 | Ap2n-kaz |
| KP090299 | Ap2n | 2n | NA | ELA3 | GenBank | No | 0.00 | 557 | dataset5 | 0 | 0 | NA | 0 | Ap2n-kaz |
| KP090300 | Ap2n | 2n | NA | ELA4 | GenBank | No | 0.00 | 557 | dataset5 | 0 | 0 | NA | 0 | Ap2n-kaz |
| KP090301 | Ap2n | 2n | NA | ELA5 | GenBank | No | 0.00 | 557 | dataset5 | 0 | 0 | NA | 0 | Ap2n-kaz |
| KP090302 | Ap2n | 2n | NA | ELA6 | GenBank | No | 0.00 | 557 | dataset5 | 0 | 0 | NA | 0 | Ap2n-kaz |
| KP090303 | Ap2n | 2n | NA | ELM1 | GenBank | No | 0.00 | 557 | dataset5 | 0 | 0 | NA | 0 | Ap2n-kaz |
| KP090304 | Ap2n | 2n | NA | ELM2 | GenBank | No | 0.00 | 557 | dataset5 | 0 | 0 | NA | 0 | Ap2n-kaz |
| KP090305 | Ap2n | 2n | NA | ELM3 | GenBank | No | 0.00 | 557 | dataset5 | 0 | 0 | NA | 0 | Ap2n-kaz |
| KP090306 | Ap2n | 2n | NA | ELM4 | GenBank | No | 0.00 | 557 | dataset5 | 0 | 0 | NA | 0 | Ap2n-kaz |
| KP090307 | Ap2n | 2n | NA | ELM5 | GenBank | No | 0.00 | 557 | dataset5 | 0 | 0 | NA | 0 | Ap2n-kaz |
| KP090308 | Ap2n | 2n | NA | ELM6 | GenBank | No | 0.00 | 557 | dataset5 | 0 | 0 | NA | 0 | Ap2n-kaz |
| KP090309 | Ap2n | 2n | NA | ELM7 | GenBank | No | 0.00 | 557 | dataset5 | 0 | 0 | NA | 0 | Ap2n-kaz |
| KP090310 | Ap2n | 2n | NA | ELM8 | GenBank | No | 0.00 | 557 | dataset5 | 0 | 0 | NA | 0 | Ap2n-kaz |
| KP090311 | Ap2n | 2n | NA | ELM9 | GenBank | No | 0.00 | 557 | dataset5 | 0 | 0 | NA | 0 | Ap2n-kaz |
| KP090312 | Ap2n | 2n | NA | SAI1 | GenBank | No | 0.00 | 557 | dataset5 | 0 | 0 | NA | 0 | Ap2n-kaz |
| KP090313 | Ap2n | 2n | NA | SAI2 | GenBank | No | 0.00 | 557 | dataset5 | 0 | 0 | NA | 0 | Ap2n-kaz |
| KP090314 | Ap2n | 2n | NA | SAI3 | GenBank | No | 0.00 | 557 | dataset5 | 0 | 0 | NA | 0 | Ap2n-kaz |
| KP090315 | Ap2n | 2n | NA | SAI4 | GenBank | No | 0.00 | 557 | dataset5 | 0 | 0 | NA | 0 | Ap2n-kaz |
| KP090316 | Ap2n | 2n | NA | SAI5 | GenBank | No | 0.00 | 557 | dataset5 | 0 | 0 | NA | 0 | Ap2n-kaz |
| KP090317 | Ap2n | 2n | NA | SAI6 | GenBank | No | 0.00 | 557 | dataset5 | 0 | 0 | NA | 0 | Ap2n-kaz |
| KU183949 | Ap2n | 2n | NA | BRK12n | GenBank | No | 0.00 | 578 | dataset5 | 0 | 0 | NA | 0 | Ap2n-kaz |
| KU183950 | Ap2n | 2n | NA | BRK22n | GenBank | No | 0.00 | 578 | dataset5 | 0 | 0 | NA | 0 | Ap2n-kaz |
| KU183951 | Ap2n | 2n | NA | BRK32n | GenBank | No | 0.00 | 578 | dataset5 | 0 | 0 | 2 | 1 | Ap2n |
| KU183952 | Ap2n | 2n | NA | BRK42n | GenBank | No | 0.00 | 578 | dataset5 | 0 | 0 | NA | 0 | Ap2n-kaz |
| KU183953 | Ap2n | 2n | NA | BRK52n | GenBank | No | 0.00 | 578 | dataset5 | 0 | 0 | NA | 0 | Ap2n-kaz |
| KU183961 | Ap2n | 2n | NA | GAH12n | GenBank | No | 0.00 | 554 | dataset5 | 0 | 0 | NA | 0 | Ap2n-kaz |
| KU183962 | Ap2n | 2n | NA | GAH22n | GenBank | No | 0.00 | 554 | dataset5 | 0 | 0 | NA | 0 | Ap2n-kaz |
| KU183963 | Ap2n | 2n | NA | GAH32n | GenBank | No | 0.00 | 554 | dataset5 | 0 | 0 | NA | 0 | Ap2n-kaz |
| KU183964 | Ap2n | 2n | NA | GAH42n | GenBank | No | 0.00 | 554 | dataset5 | 0 | 0 | NA | 0 | Ap2n-kaz |
| KU183965 | Ap2n | 2n | NA | GAH52n | GenBank | No | 0.00 | 554 | dataset5 | 0 | 0 | NA | 0 | Ap2n-kaz |
| KU183966 | Ap2n | 2n | NA | GAH62n | GenBank | No | 0.00 | 554 | dataset5 | 0 | 0 | NA | 0 | Ap2n-kaz |
| KU183967 | Ap2n | 2n | NA | GAH72n | GenBank | No | 0.00 | 554 | dataset5 | 0 | 0 | NA | 0 | Ap2n-kaz |
| KU183978 | Ap2n | 2n | NA | AQ12n | GenBank | No | 0.00 | 574 | dataset5 | 0.04 | 0 | NA | 0 | Ap2n-urm |
| KU183979 | Ap2n | 2n | NA | AQ22n | GenBank | No | 0.00 | 573 | dataset5 | 0.04 | 0 | NA | 0 | Ap2n-urm |
| KU183980 | Ap2n | 2n | NA | AQ32n | GenBank | No | 0.00 | 571 | dataset5 | 0.04 | 0 | NA | 0 | Ap2n-urm |
| KU183981 | Ap2n | 2n | NA | AQ42n | GenBank | No | 0.00 | 574 | dataset5 | 0.04 | 0 | NA | 0 | Ap2n-urm |
| KU183982 | Ap2n | 2n | NA | AQ52n | GenBank | No | 0.00 | 570 | dataset5 | 0.04 | 0 | NA | 0 | Ap2n-urm |
| KU183983 | Ap2n | 2n | NA | AB12n | GenBank | No | 0.00 | 574 | dataset5 | 0 | 0 | 1 | 0 | Ap2n-kaz |
| KU183984 | Ap2n | 2n | NA | AB22n | GenBank | No | 0.00 | 570 | dataset5 | 0 | 0 | NA | 0 | Ap2n-kaz |
| KU183985 | Ap2n | 2n | NA | AB32n | GenBank | No | 0.00 | 574 | dataset5 | 0 | 0 | NA | 0 | Ap2n-kaz |
| KU183986 | Ap2n | 2n | NA | AB42n | GenBank | No | 0.00 | 574 | dataset5 | 0 | 0 | NA | 0 | Ap2n-kaz |
| KU183987 | Ap2n | 2n | NA | AB52n | GenBank | No | 0.00 | 570 | dataset5 | 0 | 0 | NA | 0 | Ap2n-kaz |
| AY953368 | Ap2n | 2n | NA | Aus1 | GenBank | No | 0.00 | 531 | dataset6 | NA | NA | NA | 0 | Ap2n-kaz |
| AY953369 | Ap2n | 2n | NA | Aus2 | GenBank | No | 0.00 | 531 | dataset6 | NA | NA | NA | 0 | Ap2n-kaz |
| FJ790946 | Ap2n | 2n | NA | haplotype1 | GenBank | No | 0.00 | 604 | dataset5 | 0 | 0 | NA | 0 | Ap2n-kaz |
| FJ790947 | Ap2n | 2n | NA | haplotype2 | GenBank | No | 0.00 | 604 | dataset5 | 0 | 0 | NA | 0 | Ap2n-kaz |
| FJ790948 | Ap2n | 2n | NA | haplotype3 | GenBank | No | 0.00 | 604 | dataset5 | 0 | 0 | NA | 0 | Ap2n-kaz |
| KX925405 | Ap2n | 2n | NA | APAR.1 | GenBank | No | 0.00 | 1412 | dataset6 | NA | NA | NA | 1 | Ap2n |
| KX925406 | Ap2n | 2n | NA | APAR.2 | GenBank | No | 0.00 | 1413 | dataset6 | NA | NA | NA | 1 | Ap2n |
| KX925407 | Ap2n | 2n | NA | APAR.3 | GenBank | No | 0.00 | 1406 | dataset6 | NA | NA | NA | 0 | Ap2n-kaz |
| NA | Ap3n | 3n | ANK | Ap3nANK1 | This study | No | 0.00 | 521 | dataset2 | NA | NA | NA | 0 | Ap3n |
| NA | Ap3n | 3n | ANK | Ap3nANK7 | This study | No | 0.00 | 521 | dataset2 | NA | NA | NA | 0 | Ap3n |
| NA | Ap3n | 3n | ANK | Ap3nANK2 | This study | No | 0.00 | 627 | dataset2 | NA | NA | NA | 0 | Ap3n |
| MT791797 | Ap3n | 3n | ANK | Ap3nANK3 | This study | No | 0.00 | 627 | dataset1 | 0 | 0 | NA | 0 | Ap3n |
| NA | Ap3n | 3n | ANK | Ap3nANK4 | This study | No | 0.01 | 620 | dataset2 | NA | NA | NA | 0 | Ap3n |
| MT791798 | Ap3n | 3n | ANK | Ap3nANK5 | This study | No | 0.00 | 627 | dataset1 | 0 | 0 | NA | 0 | Ap3n |
| NA | Ap3n | 3n | ANK | Ap3nANK6 | This study | No | 0.01 | 601 | dataset2 | NA | NA | NA | 1 | Ap3n |
| MT791799 | Ap3n | 3n | ANK | Ap3nANK8 | This study | No | 0.00 | 627 | dataset1 | 0.06 | 0 | 1 | 0 | Ap3n |
| MT791800 | Ap3n | 3n | ANK | Ap3nANK9 | This study | No | 0.00 | 627 | dataset1 | 0 | 0 | NA | 0 | Ap3n |
| NA | Ap3n | 3n | IZM | Ap3nIZM2.1 | This study | No | 0.00 | 627 | dataset1 | 0 | 0 | 3 | 1 | Ap3n |
| NA | Ap3n | 3n | IZM | Ap3nIZM1 | This study | No | 0.01 | 597 | dataset2 | NA | NA | NA | 0 | Ap3n |
| NA | Ap3n | 3n | IZM | Ap3nIZM3 | This study | No | 0.04 | 625 | dataset2 | NA | NA | NA | 1 | Ap3n |
| NA | Ap3n | 3n | IZM | Ap3nIZM4.1 | This study | No | 0.02 | 547 | dataset2 | NA | NA | NA | 0 | Ap3n |
| NA | Ap3n | 3n | IZM | Ap3nIZM2.2 | This study | No | 0.00 | 637 | dataset2 | NA | NA | NA | 1 | Ap3n |
| MT791802 | Ap3n | 3n | NAR | Ap3nNAR1 | This study | No | 0.00 | 633 | dataset1 | 0 | 0 | NA | 0 | Ap3n |
| NA | Ap3n | 3n | NAR | Ap3nNAR2 | This study | No | 0.00 | 635 | dataset2 | NA | NA | NA | 0 | Ap3n |
| MT791803 | Ap3n | 3n | NAR | Ap3nNAR3 | This study | No | 0.00 | 633 | dataset1 | 0 | 0 | 1 | 0 | Ap3n |
| NA | Ap3n | 3n | IZM | Ap3nIZM4.2 | This study | No | 0.00 | 515 | dataset2 | NA | NA | NA | 1 | Ap3n |
| MT791804 | Ap3n | 3n | NAR | Ap3nNAR5 | This study | No | 0.00 | 621 | dataset1 | 0 | 0 | NA | 0 | Ap3n |
| MT791801 | Ap3n | 3n | ANK | Ap3nANK10 | This study | Yes | 0.00 | 600 | dataset3 | 0.06 | 0 | NA | 0 | Ap3n |
| KP090318 | Ap3n | 3n | NA | ANK1 | GenBank | No | 0.00 | 557 | dataset5 | 0.04 | 0 | 0 | 0 | Ap3n |
| KP090319 | Ap3n | 3n | NA | ANK2 | GenBank | No | 0.00 | 557 | dataset5 | 0 | 0 | NA | 0 | Ap3n |
| KP090320 | Ap3n | 3n | NA | ANK3 | GenBank | No | 0.00 | 557 | dataset5 | 0 | 0 | NA | 0 | Ap3n |
| KP090321 | Ap3n | 3n | NA | ANK4 | GenBank | No | 0.00 | 557 | dataset5 | 0.04 | 0 | NA | 0 | Ap3n |
| KP090322 | Ap3n | 3n | NA | ANK5 | GenBank | No | 0.00 | 557 | dataset5 | 0 | 0 | NA | 0 | Ap3n |
| KP090323 | Ap3n | 3n | NA | ANK6 | GenBank | No | 0.00 | 557 | dataset5 | 0 | 0 | NA | 0 | Ap3n |
| KP090324 | Ap3n | 3n | NA | ANK7 | GenBank | No | 0.00 | 557 | dataset5 | 0 | 0 | NA | 0 | Ap3n |
| KU183988 | Ap3n | 3n | NA | AB13n | GenBank | No | 0.00 | 578 | dataset5 | 0 | 0 | NA | 0 | Ap3n |
| KU183989 | Ap3n | 3n | NA | AB23n | GenBank | No | 0.00 | 578 | dataset5 | 0 | 0 | NA | 0 | Ap3n |
| KU183990 | Ap3n | 3n | NA | AB33n | GenBank | No | 0.00 | 578 | dataset5 | 0 | 0 | NA | 0 | Ap3n |
| KU183991 | Ap3n | 3n | NA | AB43n | GenBank | No | 0.00 | 578 | dataset5 | 0 | 0 | NA | 0 | Ap3n |
| KU183992 | Ap3n | 3n | NA | AB53n | GenBank | No | 0.00 | 578 | dataset5 | 0 | 0 | NA | 0 | Ap3n |
| NA | Ap4n | 4n | IZM | numt3Ap4nIZM1 | This study | No | 0.01 | 582 | dataset2 | NA | NA | NA | 1 | Ap4n |
| NA | Ap4n | 4n | IZM | Ap4nIZM2 | This study | No | 0.01 | 593 | dataset2 | NA | NA | NA | 1 | Ap4n |
| NA | Ap4n | 4n | LAV | Ap4nLAV1 | This study | No | 0.01 | 550 | dataset2 | NA | NA | NA | 1 | Ap4n |
| NA | Ap4n | 4n | LAV | Ap4nLAV2 | This study | No | 0.01 | 498 | dataset2 | NA | NA | NA | 1 | Ap4n |
| NA | Ap4n | 4n | LAV | Ap4nLAV3 | This study | No | 0.01 | 508 | dataset2 | NA | NA | NA | 1 | Ap4n |
| NA | Ap4n | 4n | LAV | Ap4nLAV4.1 | This study | No | 0.01 | 595 | dataset2 | NA | NA | NA | 1 | Ap4n |
| NA | Ap4n | 4n | LAV | Ap4nLAV5 | This study | No | 0.01 | 568 | dataset2 | NA | NA | NA | 1 | Ap4n |
| NA | Ap4n | 4n | LAV | Ap4nLAV6.1 | This study | No | 0.01 | 550 | dataset2 | NA | NA | NA | 1 | Ap4n |
| NA | Ap4n | 4n | TEK | Ap4nTEK1 | This study | No | 0.02 | 559 | dataset2 | NA | NA | NA | 1 | Ap4n |
| NA | Ap4n | 4n | TEK | Ap4nTEK2 | This study | No | 0.02 | 559 | dataset2 | NA | NA | NA | 1 | Ap4n |
| NA | Ap4n | 4n | TEK | Ap4nTEK3.1 | This study | No | 0.02 | 559 | dataset2 | NA | NA | NA | 1 | Ap4n |
| NA | Ap4n | 4n | TEK | Ap4nTEK4.1 | This study | No | 0.01 | 510 | dataset2 | NA | NA | NA | 1 | Ap4n |
| NA | Ap4n | 4n | TEK | Ap4nTEK5 | This study | No | 0.02 | 495 | dataset2 | NA | NA | NA | 1 | Ap4n |
| NA | Ap4n | 4n | TEK | Ap4nTEK6 | This study | No | 0.03 | 513 | dataset2 | NA | NA | NA | 1 | Ap4n |
| NA | Ap4n | 4n | TEK | Ap4nTEK7 | This study | No | 0.02 | 517 | dataset2 | NA | NA | NA | 1 | Ap4n |
| MT791805 | Ap4n | 4n | ARS | Ap4nARS1 | This study | No | 0.00 | 688 | dataset1 | 0.04 | 0 | 1 | 0 | Ap4n |
| MT791806 | Ap4n | 4n | ARS | Ap4nARS2 | This study | No | 0.00 | 686 | dataset1 | 0.04 | 0 | NA | 0 | Ap4n |
| MT791807 | Ap4n | 4n | ARS | Ap4nARS3 | This study | No | 0.00 | 687 | dataset1 | 0.04 | 0 | NA | 0 | Ap4n |
| MT791808 | Ap4n | 4n | ARS | Ap4nARS4 | This study | No | 0.00 | 687 | dataset1 | 0.04 | 0 | NA | 0 | Ap4n |
| NA | Ap4n | 4n | BET | Ap4nBET1 | This study | No | 0.00 | 685 | dataset2 | NA | NA | NA | 0 | Ap4n |
| MT791809 | Ap4n | 4n | BET | Ap4nBET2 | This study | No | 0.00 | 683 | dataset1 | 0.04 | 0 | NA | 0 | Ap4n |
| NA | Ap4n | 4n | BET | Ap4nBET3 | This study | No | 0.00 | 685 | dataset2 | NA | NA | NA | 0 | Ap4n |
| MT791810 | Ap4n | 4n | IMO | Ap4nIMO1 | This study | No | 0.00 | 681 | dataset1 | 0.04 | 0 | NA | 0 | Ap4n |
| MT791811 | Ap4n | 4n | IMO | Ap4nIMO2 | This study | No | 0.00 | 652 | dataset1 | 0.04 | 0 | NA | 0 | Ap4n |
| MT791812 | Ap4n | 4n | IMO | Ap4nIMO3 | This study | No | 0.00 | 684 | dataset1 | 0.04 | 0 | NA | 0 | Ap4n |
| NA | Ap4n | 4n | IMO | Ap4nIMO4 | This study | No | 0.00 | 650 | dataset2 | NA | NA | NA | 0 | Ap4n |
| NA | Ap4n | 4n | LAV | Ap4nLAV4.2 | This study | No | 0.02 | 490 | dataset2 | NA | NA | NA | 1 | Ap4n |
| NA | Ap4n | 4n | TEK | Ap4nTEK4.2 | This study | No | 0.02 | 490 | dataset2 | NA | NA | NA | 1 | Ap4n |
| NA | Ap4n | 4n | LAV | Ap4nLAV6.2 | This study | No | 0.02 | 305 | dataset2 | NA | NA | NA | 1 | Ap4n |
| NA | Ap4n | 4n | TEK | Ap4nTEK3.2 | This study | No | 0.02 | 490 | dataset2 | NA | NA | NA | 1 | Ap4n |
| MT792817 | Ap4n | 4n | IZM | numt1Ap4nIZM1 | This study | Yes | 0.00 | 650 | dataset3 | 3.27 | 1 | 23 | 1 | Ap4n |
| MT791813 | Ap4n | 4n | IZM | Ap4nIZM1 | This study | Yes | 0.00 | 655 | dataset3 | 0.04 | 0 | 0 | 0 | Ap4n |
| MT792818 | Ap4n | 4n | IZM | numt2Ap4nIZM1 | This study | Yes | 0.00 | 637 | dataset3 | 0 | 0 | 1 | 1 | Ap4n |
| MT792819 | Ap4n | 4n | TEK | numt1Ap4nTEK3 | This study | Yes | 0.00 | 638 | dataset3 | 0.04 | 0 | 5 | 1 | Ap4n |
| MT792820 | Ap4n | 4n | TEK | numt2Ap4nTEK3 | This study | Yes | 0.00 | 638 | dataset3 | 3.76 | 1 | 16 | 1 | Ap4n |
| KU183954 | Ap4n | 4n | NA | BRK14n | GenBank | No | 0.00 | 578 | dataset5 | 0.04 | 0 | NA | 0 | Ap4n |
| KU183955 | Ap4n | 4n | NA | BRK24n | GenBank | No | 0.00 | 578 | dataset5 | 4.61 | 1 | 0 | 1 | Ap4n |
| KU183956 | Ap4n | 4n | NA | BRK34n | GenBank | No | 0.00 | 578 | dataset5 | 0.04 | 0 | 1 | 0 | Ap4n |
| KU183957 | Ap4n | 4n | NA | BRK44n | GenBank | No | 0.00 | 579 | dataset5 | 0.04 | 0 | 1 | 0 | Ap4n |
| KU183972 | Ap4n | 4n | NA | HOH14n | GenBank | Yes | 0.00 | 578 | dataset5 | 0.04 | 0 | 5 | 1 | Ap4n |
| KU183973 | Ap4n | 4n | NA | HOH34n | GenBank | Yes | 0.00 | 578 | dataset5 | 0.04 | 0 | 8 | 1 | Ap4n |
| KU183974 | Ap4n | 4n | NA | HOH44n | GenBank | Yes | 0.00 | 554 | dataset5 | 0.04 | 0 | 6 | 1 | Ap4n |
| KU183975 | Ap4n | 4n | NA | HOH54n | GenBank | Yes | 0.00 | 554 | dataset5 | 0.04 | 0 | 3 | 1 | Ap4n |
| KU183976 | Ap4n | 4n | NA | HOH64n | GenBank | Yes | 0.00 | 554 | dataset5 | 0.04 | 0 | NA | 1 | Ap4n |
| KU183977 | Ap4n | 4n | NA | HOH74n | GenBank | Yes | 0.00 | 554 | dataset5 | 0.04 | 0 | NA | 1 | Ap4n |
| NA | Ap5n | 5n | ATA2 | Ap5nATA21 | This study | No | 0.00 | 550 | dataset2 | NA | NA | NA | 1 | Ap5n |
| NA | Ap5n | 5n | ATA2 | Ap5nATA22 | This study | No | 0.01 | 568 | dataset2 | NA | NA | NA | 1 | Ap5n |
| NA | Ap5n | 5n | ATA2 | Ap5nATA23 | This study | No | 0.00 | 559 | dataset2 | NA | NA | NA | 1 | Ap5n |
| NA | Ap5n | 5n | ATA2 | Ap5nATA24 | This study | No | 0.00 | 566 | dataset2 | NA | NA | NA | 1 | Ap5n |
| NA | Ap5n | 5n | IZM | Ap5nIZM3 | This study | No | 0.00 | 568 | dataset2 | NA | NA | NA | 1 | Ap5n |
| NA | Ap5n | 5n | IZM | Ap5nIZM5 | This study | No | 0.01 | 559 | dataset2 | NA | NA | NA | 1 | Ap5n |
| NA | Ap5n | 5n | BUJ | Ap5nBUJ2.1 | This study | No | 0.01 | 562 | dataset2 | NA | NA | NA | 1 | Ap5n |
| NA | Ap5n | 5n | BUJ | Ap5nBUJ3 | This study | No | 0.00 | 686 | dataset2 | NA | NA | NA | 0 | Ap5n |
| MT791814 | Ap5n | 5n | BUJ | Ap5nBUJ4 | This study | No | 0.00 | 686 | dataset1 | 0.04 | 0 | 1 | 0 | Ap5n |
| NA | Ap5n | 5n | BUJ | Ap5nBUJ1 | This study | No | 0.01 | 686 | dataset2 | NA | NA | NA | 0 | Ap5n |
| NA | Ap5n | 5n | CIT | Ap5nCIT1 | This study | No | 0.01 | 683 | dataset2 | NA | NA | NA | 0 | Ap5n |
| NA | Ap5n | 5n | CIT | Ap5nCIT2 | This study | No | 0.02 | 490 | dataset2 | NA | NA | NA | 0 | Ap5n |
| NA | Ap5n | 5n | CIT | Ap5nCIT3 | This study | No | 0.01 | 678 | dataset2 | NA | NA | NA | 0 | Ap5n |
| NA | Ap5n | 5n | CIT | Ap5nCIT4 | This study | No | 0.00 | 682 | dataset2 | NA | NA | NA | 0 | Ap5n |
| NA | Ap5n | 5n | DON | Ap5nDON1 | This study | No | 0.03 | 497 | dataset2 | NA | NA | NA | 1 | Ap5n |
| NA | Ap5n | 5n | DON | Ap5nDON2.1 | This study | No | 0.01 | 671 | dataset2 | NA | NA | NA | 0 | Ap5n |
| NA | Ap5n | 5n | IZM | Ap5nIZM1.1 | This study | No | 0.03 | 464 | dataset2 | NA | NA | NA | 1 | Ap5n |
| NA | Ap5n | 5n | IZM | Ap5nIZM2 | This study | No | 0.05 | 479 | dataset2 | NA | NA | NA | 1 | Ap5n |
| NA | Ap5n | 5n | IZM | Ap5nIZM4 | This study | No | 0.04 | 494 | dataset2 | NA | NA | NA | 1 | Ap5n |
| NA | Ap5n | 5n | ATA1 | Ap5nATA11 | This study | No | 0.04 | 359 | dataset2 | NA | NA | NA | 1 | Ap5n |
| NA | Ap5n | 5n | ATA1 | Ap5nATA15 | This study | No | 0.01 | 507 | dataset2 | NA | NA | NA | 1 | Ap5n |
| NA | Ap5n | 5n | ATA1 | Ap5nATA13 | This study | No | 0.01 | 451 | dataset2 | NA | NA | NA | 1 | Ap5n |
| NA | Ap5n | 5n | BUJ | Ap5nBUJ10 | This study | No | 0.03 | 489 | dataset2 | NA | NA | NA | 1 | Ap5n |
| NA | Ap5n | 5n | BUJ | Ap5nBUJ12 | This study | No | 0.01 | 510 | dataset2 | NA | NA | NA | 1 | Ap5n |
| NA | Ap5n | 5n | BUJ | Ap5nBUJ6 | This study | No | 0.02 | 490 | dataset2 | NA | NA | NA | 1 | Ap5n |
| NA | Ap5n | 5n | BUJ | numtAp5nBUJ8 | This study | No | 0.03 | 494 | dataset2 | NA | NA | NA | 1 | Ap5n |
| NA | Ap5n | 5n | CIT | Ap5nCIT5 | This study | No | 0.00 | 533 | dataset2 | NA | NA | NA | 1 | Ap5n |
| NA | Ap5n | 5n | ATA1 | Ap5nATA14 | This study | No | 0.00 | 520 | dataset2 | NA | NA | NA | 1 | Ap5n |
| NA | Ap5n | 5n | ATA1 | Ap5nATA12 | This study | No | 0.05 | 454 | dataset2 | NA | NA | NA | 1 | Ap5n |
| NA | Ap5n | 5n | BUJ | Ap5nBUJ2.2 | This study | No | 0.01 | 509 | dataset2 | NA | NA | NA | 1 | Ap5n |
| NA | Ap5n | 5n | BUJ | Ap5nBUJ11 | This study | No | 0.02 | 506 | dataset2 | NA | NA | NA | 1 | Ap5n |
| NA | Ap5n | 5n | BUJ | Ap5nBUJ13 | This study | No | 0.01 | 501 | dataset2 | NA | NA | NA | 1 | Ap5n |
| NA | Ap5n | 5n | BUJ | Ap5nBUJ5 | This study | No | 0.00 | 510 | dataset2 | NA | NA | NA | 1 | Ap5n |
| NA | Ap5n | 5n | BUJ | Ap5nBUJ7 | This study | No | 0.03 | 488 | dataset2 | NA | NA | NA | 1 | Ap5n |
| NA | Ap5n | 5n | BUJ | Ap5nBUJ9 | This study | No | 0.01 | 496 | dataset2 | NA | NA | NA | 1 | Ap5n |
| NA | Ap5n | 5n | CIT | Ap5nCIT6 | This study | No | 0.00 | 494 | dataset2 | NA | NA | NA | 1 | Ap5n |
| NA | Ap5n | 5n | IZM | Ap5nIZM1.2 | This study | No | 0.00 | 495 | dataset2 | NA | NA | NA | 1 | Ap5n |
| NA | Ap5n | 5n | DON | Ap5nDON2.2 | This study | No | 0.02 | 487 | dataset2 | NA | NA | NA | 0 | Ap5n |
| MT792834 | Ap5n | 5n | IZM | numt1Ap5nIZM2 | This study | Yes | 0.00 | 650 | dataset3 | 5.4 | 1 | 2 | 1 | Ap5n |
| MT792835 | Ap5n | 5n | IZM | numt2Ap5nIZM2 | This study | Yes | 0.00 | 621 | dataset3 | 1.22 | 1 | 0 | 1 | Ap5n |
| MT792836 | Ap5n | 5n | IZM | numt3Ap5nIZM2 | This study | Yes | 0.00 | 621 | dataset3 | 5.7 | 1 | 20 | 1 | Ap5n |
| MT792837 | Ap5n | 5n | IZM | numt4Ap5nIZM2 | This study | Yes | 0.00 | 621 | dataset3 | 5.09 | 1 | 7 | 1 | Ap5n |
| MT792838 | Ap5n | 5n | IZM | numt5Ap5nIZM2 | This study | Yes | 0.00 | 630 | dataset3 | 5.34 | 1 | 4 | 1 | Ap5n |
| MT792839 | Ap5n | 5n | IZM | numt6Ap5nIZM2 | This study | Yes | 0.00 | 637 | dataset3 | 2.81 | 1 | 3 | 1 | Ap5n |
| MT792840 | Ap5n | 5n | IZM | numt7Ap5nIZM2 | This study | Yes | 0.00 | 637 | dataset3 | 4.42 | 1 | 1 | 1 | Ap5n |
| MT792841 | Ap5n | 5n | IZM | numt8Ap5nIZM2 | This study | Yes | 0.00 | 638 | dataset3 | 0.24 | 1 | 14 | 1 | Ap5n |
| MT792827 | Ap5n | 5n | CIT | numt1Ap5nCIT6 | This study | Yes | 0.00 | 638 | dataset3 | 0 | 0 | 0 | 1 | Ap5n |
| MT792828 | Ap5n | 5n | CIT | numt2Ap5nCIT6 | This study | Yes | 0.00 | 608 | dataset3 | 1.36 | 1 | 4 | 1 | Ap5n |
| MT792829 | Ap5n | 5n | CIT | numt3Ap5nCIT6 | This study | Yes | 0.00 | 608 | dataset3 | 0 | 0 | 8 | 1 | Ap5n |
| MT792830 | Ap5n | 5n | CIT | numt4Ap5nCIT6 | This study | Yes | 0.00 | 650 | dataset3 | 4.61 | 1 | 15 | 1 | Ap5n |
| MT792821 | Ap5n | 5n | BUJ | numt6Ap5nBUJ2 | This study | Yes | 0.00 | 606 | dataset3 | 0 | 0 | NA | 1 | Ap5n |
| MT792822 | Ap5n | 5n | BUJ | numt2Ap5nBUJ2 | This study | Yes | 0.00 | 638 | dataset3 | 0.1 | 0 | 5 | 1 | Ap5n |
| MT792823 | Ap5n | 5n | BUJ | numt3Ap5nBUJ2 | This study | Yes | 0.00 | 642 | dataset3 | 0 | 0 | 1 | 1 | Ap5n |
| MT792824 | Ap5n | 5n | BUJ | numt4Ap5nBUJ2 | This study | Yes | 0.00 | 606 | dataset3 | 2.64 | 1 | 24 | 1 | Ap5n |
| MT792825 | Ap5n | 5n | BUJ | numt5Ap5nBUJ2 | This study | Yes | 0.00 | 606 | dataset3 | 6.09 | 1 | 6 | 1 | Ap5n |
| MT792826 | Ap5n | 5n | BUJ | numt1Ap5nBUJ2 | This study | Yes | 0.00 | 637 | dataset3 | 0 | 0 | NA | 1 | Ap5n |
| MT792832 | Ap5n | 5n | DON | numt1Ap5nDON2 | This study | Yes | 0.00 | 638 | dataset3 | 3.15 | 1 | 1 | 1 | Ap5n |
| MT792833 | Ap5n | 5n | DON | numt2Ap5nDON2 | This study | Yes | 0.00 | 621 | dataset3 | 1.4 | 1 | 3 | 1 | Ap5n |
| MT791816 | Ap5n | 5n | DON | Ap5nDON2 | This study | Yes | 0.00 | 637 | dataset3 | 0.04 | 0 | NA | 0 | Ap5n |
| MT791815 | Ap5n | 5n | CIT | Ap5nCIT7 | This study | Yes | 0.00 | 553 | dataset3 | 0.04 | 0 | NA | 0 | Ap5n |
| MT792831 | Ap5n | 5n | CIT | numt1Ap5nCIT7 | This study | Yes | 0.00 | 553 | dataset3 | 1.26 | 1 | 1 | 1 | Ap5n |
| KU183959 | Ap5n | 5n | NA | BRK35n | GenBank | Yes | 0.00 | 578 | dataset5 | 0.04 | 0 | NA | 0 | Ap5n |
| KU183960 | Ap5n | 5n | NA | BRK75n | GenBank | Yes | 0.00 | 578 | dataset5 | 0.04 | 0 | NA | 0 | Ap5n |
| KU183968 | Ap5n | 5n | NA | YGH15n | GenBank | No | 0.00 | 554 | dataset5 | 14.41 | 1 | 14 | 1 | Ap5n |
| KU183969 | Ap5n | 5n | NA | YGH25n | GenBank | No | 0.00 | 554 | dataset5 | 9.11 | 1 | 17 | 1 | Ap5n |
| KU183970 | Ap5n | 5n | NA | YGH35n | GenBank | No | 0.00 | 554 | dataset5 | 10.33 | 1 | 17 | 1 | Ap5n |
| KU183971 | Ap5n | 5n | NA | YGH45n | GenBank | No | 0.00 | 554 | dataset5 | 1.98 | 1 | 6 | 1 | Ap5n |
| MT791817 | Asin | 2n | DON | AsinDON1 | This study | No | 0.00 | 686 | dataset1 | 0.04 | 0 | 1 | 0 | Asin |
| MT791820 | Asin | 2n | YIM | AsinYIM1 | This study | No | 0.00 | 626 | dataset1 | 0.04 | 0 | 1 | 0 | Asin |
| NA | Asin | 2n | YIM | AsinYIM3 | This study | No | 0.02 | 637 | dataset2 | NA | NA | NA | 0 | Asin |
| NA | Asin | 2n | XIE | AsinXIE1 | This study | No | 0.01 | 594 | dataset2 | NA | NA | NA | 0 | Asin |
| NA | Asin | 2n | XIE | AsinXIE8 | This study | No | 0.00 | 639 | dataset2 | NA | NA | NA | 0 | Asin |
| NA | Asin | 2n | XIE | AsinXIE3 | This study | No | 0.00 | 638 | dataset2 | NA | NA | NA | 0 | Asin |
| MT791818 | Asin | 2n | XIE | AsinXIE6 | This study | No | 0.00 | 633 | dataset1 | 0.04 | 0 | 1 | 0 | Asin |
| NA | Asin | 2n | XIE | AsinXIE5 | This study | No | 0.05 | 354 | dataset2 | NA | NA | NA | 1 | Asin |
| MT791819 | Asin | 2n | XIE | AsinXIE7 | This study | No | 0.00 | 638 | dataset1 | 0.04 | 0 | 0 | 0 | Asin |
| NA | Asin | 2n | XIE | AsinXIE2 | This study | No | 0.03 | 457 | dataset2 | NA | NA | NA | 1 | Asin |
| NA | Asin | 2n | XIE | AsinXIE4 | This study | No | 0.01 | 633 | dataset2 | NA | NA | NA | 1 | Asin |
| DQ119648 | Asin | 2n | NA | AsinicaYCHou | GenBank | No | 0.00 | 658 | dataset5 | 0.04 | 0 | 1 | 0 | Asin |
| DQ119650 | Asin | 2n | NA | AsinicaSQZHou | GenBank | No | 0.00 | 658 | dataset5 | 0.04 | 0 | 1 | 0 | Asin |
| DQ119649 | Asin | 2n | NA | AsinicaNLHou | GenBank | No | 0.00 | 658 | dataset5 | 1.4 | 1 | 0 | 1 | Asin |
| LC195586 | Asin | 2n | NA | TU13 | GenBank | No | 0.00 | 658 | dataset5 | 0.04 | 0 | 5 | 1 | Asin |
| MK069595 | Asin | 2n | NA | AsARC1166 | GenBank | No | 0.00 | 688 | dataset5 | 0.04 | 0 | 1 | 0 | Asin |
| MG572075 | Asin | 2n | NA | Yuncheng2016 | GenBank | No | 0.00 | 1409 | dataset6 | NA | NA | NA | 1 | Asin |
| MG572077 | Asin | 2n | NA | Haolebaoqing1997 | GenBank | No | 0.00 | 1407 | dataset6 | NA | NA | NA | 0 | Asin |
| MG572079 | Asin | 2n | NA | Yimeng1991 | GenBank | No | 0.00 | 1406 | dataset6 | NA | NA | NA | 0 | Asin |
| MG572080 | Asin | 2n | NA | Bameng2000 | GenBank | No | 0.00 | 1394 | dataset6 | NA | NA | NA | 0 | Asin |
| MG572081 | Asin | 2n | NA | Bayanhu1994 | GenBank | No | 0.00 | 1406 | dataset6 | NA | NA | NA | 0 | Asin |
| EF615591 | Asin | 2n | NA | ARC1188 | GenBank | No | 0.00 | 622 | dataset6 | NA | NA | NA | 0 | Asin |
| EF615592 | Asin | 2n | NA | ARC1317 | GenBank | No | 0.00 | 622 | dataset6 | NA | NA | NA | 0 | Asin |
| KF707885 | Asin | 2n | NA | ASYUN3 | GenBank | No | 0.00 | 614 | dataset5 | 0.04 | 0 | 2 | 1 | Asin |
| KF707886 | Asin | 2n | NA | ASYUN1 | GenBank | No | 0.00 | 614 | dataset5 | 0.04 | 0 | 2 | 1 | Asin |
| KF707887 | Asin | 2n | NA | ASYUN5 | GenBank | No | 0.00 | 614 | dataset5 | 0.04 | 0 | NA | 1 | Asin |
| KF707888 | Asin | 2n | NA | ASYUN6 | GenBank | No | 0.00 | 614 | dataset5 | 0.04 | 0 | NA | 1 | Asin |
| KF707889 | Asin | 2n | NA | ASYUN8 | GenBank | No | 0.00 | 614 | dataset5 | 0.04 | 0 | NA | 1 | Asin |
| KF707890 | Asin | 2n | NA | ASYUN9 | GenBank | No | 0.00 | 614 | dataset5 | 0.04 | 0 | NA | 1 | Asin |
| KF707891 | Asin | 2n | NA | ASYUN10 | GenBank | No | 0.00 | 614 | dataset5 | 0.04 | 0 | NA | 1 | Asin |
| KF707892 | Asin | 2n | NA | ASYUN2 | GenBank | No | 0.00 | 614 | dataset5 | 0.04 | 0 | NA | 1 | Asin |
| KF707893 | Asin | 2n | NA | ASYUN4 | GenBank | No | 0.00 | 614 | dataset5 | 0.04 | 0 | NA | 1 | Asin |
| KF707894 | Asin | 2n | NA | ASYUN7 | GenBank | No | 0.00 | 614 | dataset5 | 0.04 | 0 | NA | 1 | Asin |
| KF691157 | Asin | 2n | NA | BEI1 | GenBank | No | 0.00 | 561 | dataset5 | 0.29 | 1 | 6 | 1 | Asin |
| KF691158 | Asin | 2n | NA | BEI2 | GenBank | No | 0.00 | 561 | dataset5 | 0.29 | 1 | 6 | 1 | Asin |
| KF691269 | Asin | 2n | NA | XIE1 | GenBank | No | 0.00 | 561 | dataset5 | 0.13 | 1 | 2 | 1 | Asin |
| KF691270 | Asin | 2n | NA | XIE2 | GenBank | No | 0.00 | 561 | dataset5 | 0.19 | 1 | 1 | 1 | Asin |
| KF691271 | Asin | 2n | NA | XIE3 | GenBank | No | 0.00 | 561 | dataset5 | 0.19 | 1 | 1 | 1 | Asin |
| KF691272 | Asin | 2n | NA | XIE4 | GenBank | No | 0.00 | 561 | dataset5 | 0.04 | 0 | 1 | 0 | Asin |
| KF691273 | Asin | 2n | NA | XIE5 | GenBank | No | 0.00 | 561 | dataset5 | 0.04 | 0 | 1 | 0 | Asin |
| KF691274 | Asin | 2n | NA | XIE6 | GenBank | No | 0.00 | 561 | dataset5 | 0.04 | 0 | 1 | 0 | Asin |
| KF691275 | Asin | 2n | NA | XIE7 | GenBank | No | 0.00 | 561 | dataset5 | 0.04 | 0 | 1 | 0 | Asin |
| KF691276 | Asin | 2n | NA | XIE8 | GenBank | No | 0.00 | 561 | dataset5 | 0.04 | 0 | NA | 0 | Asin |
| KF691277 | Asin | 2n | NA | XIE9 | GenBank | No | 0.00 | 561 | dataset5 | 0.04 | 0 | NA | 0 | Asin |
| KF691298 | Asin | 2n | NA | YUN1 | GenBank | No | 0.00 | 561 | dataset5 | 0.04 | 0 | NA | 0 | Asin |
| KF691299 | Asin | 2n | NA | YUN2 | GenBank | No | 0.00 | 561 | dataset5 | 0.04 | 0 | NA | 0 | Asin |
| KF691300 | Asin | 2n | NA | YUN3 | GenBank | No | 0.00 | 561 | dataset5 | 0.04 | 0 | NA | 0 | Asin |
| KF691301 | Asin | 2n | NA | YUN4 | GenBank | No | 0.00 | 561 | dataset5 | 1.4 | 1 | 1 | 1 | Asin |
| KF691302 | Asin | 2n | NA | YUN5 | GenBank | No | 0.00 | 561 | dataset5 | 0.04 | 0 | NA | 0 | Asin |
| KX925413 | Asin | 2n | NA | ASIN.1 | GenBank | No | 0.00 | 1406 | dataset6 | NA | NA | NA | 0 | Asin |
| KX925414 | Asin | 2n | NA | ASIN.2 | GenBank | No | 0.00 | 1396 | dataset6 | NA | NA | NA | 1 | Asin |
| KX925415 | Asin | 2n | NA | ASIN.3 | GenBank | No | 0.00 | 1393 | dataset6 | NA | NA | NA | 1 | Asin |
| KX925416 | Asin | 2n | NA | ASIN.4 | GenBank | No | 0.00 | 1413 | dataset6 | NA | NA | NA | 0 | Asin |
| MT791821 | Atib | 2n | LAG | AtibLAG13 | This study | No | 0.00 | 653 | dataset1 | 0 | 0 | 1 | 0 | Atib |
| NA | Atib | 2n | LAG | AtibLAG1 | This study | No | 0.00 | 553 | dataset2 | NA | NA | NA | 0 | Atib |
| MT791834 | Atib | 2n | LAG | AtibLAG2 | This study | No | 0.00 | 544 | dataset1 | 0 | 0 | NA | 0 | Atib |
| MT791822 | Atib | 2n | LAG | AtibLAG3 | This study | No | 0.00 | 638 | dataset1 | 0 | 0 | NA | 0 | Atib |
| MT791823 | Atib | 2n | LAG | AtibLAG4 | This study | No | 0.00 | 673 | dataset1 | 0 | 0 | NA | 0 | Atib |
| MT791824 | Atib | 2n | LAG | AtibLAG5 | This study | No | 0.00 | 662 | dataset1 | 0 | 0 | NA | 0 | Atib |
| MT791825 | Atib | 2n | LAG | AtibLAG6 | This study | No | 0.00 | 686 | dataset1 | 0 | 0 | NA | 0 | Atib |
| NA | Atib | 2n | LAG | AtibLAG7 | This study | No | 0.01 | 560 | dataset2 | NA | NA | NA | 1 | Atib |
| MT791826 | Atib | 2n | LAG | AtibLAG8 | This study | No | 0.00 | 637 | dataset1 | 0 | 0 | NA | 0 | Atib |
| MT791827 | Atib | 2n | LAG | AtibLAG9 | This study | No | 0.00 | 657 | dataset1 | 0 | 0 | NA | 0 | Atib |
| MT791828 | Atib | 2n | LAG | AtibLAG10 | This study | No | 0.00 | 584 | dataset1 | 0 | 0 | NA | 0 | Atib |
| MT791829 | Atib | 2n | LAG | AtibLAG11 | This study | No | 0.00 | 637 | dataset1 | 0 | 0 | NA | 0 | Atib |
| NA | Atib | 2n | LAG | AtibLAG12 | This study | No | 0.01 | 568 | dataset2 | NA | NA | NA | 0 | Atib |
| MT791830 | Atib | 2n | LAG | AtibLAG15 | This study | No | 0.00 | 636 | dataset1 | 0 | 0 | NA | 0 | Atib |
| NA | Atib | 2n | LAG | AtibLAG16 | This study | No | 0.01 | 592 | dataset2 | NA | NA | NA | 1 | Atib |
| NA | Atib | 2n | LAG | AtibLAG17 | This study | No | 0.00 | 631 | dataset2 | NA | NA | NA | 0 | Atib |
| MT791831 | Atib | 2n | LAG | AtibLAG18 | This study | No | 0.00 | 633 | dataset1 | 0 | 0 | NA | 0 | Atib |
| NA | Atib | 2n | LAG | AtibLAG19 | This study | No | 0.00 | 621 | dataset2 | NA | NA | NA | 0 | Atib |
| MT791832 | Atib | 2n | LAG | AtibLAG20 | This study | No | 0.00 | 642 | dataset1 | 0 | 0 | NA | 0 | Atib |
| NA | Atib | 2n | LAG | AtibLAG21 | This study | No | 0.00 | 633 | dataset2 | NA | NA | NA | 0 | Atib |
| NA | Atib | 2n | LAG | AtibLAG23 | This study | No | 0.01 | 558 | dataset2 | NA | NA | NA | 1 | Atib |
| MT791833 | Atib | 2n | LAG | AtibLAG24 | This study | Yes | 0.00 | 540 | dataset3 | 0.02 | 0 | 1 | 0 | Atib |
| MT792842 | Atib | 2n | LAG | numt1AtibLAG24 | This study | Yes | 0.00 | 540 | dataset3 | 1.84 | 1 | 1 | 1 | Atib |
| MT792843 | Atib | 2n | LAG | numt2AtibLAG24 | This study | Yes | 0.00 | 540 | dataset3 | 0.02 | 0 | 1 | 1 | Atib |
| DQ119652 | Atib | 2n | NA | AspQXCHou | GenBank | No | 0.00 | 658 | dataset6 | NA | NA | NA | 0 | Atib |
| JQ975177 | Atib | 2n | NA | AtARC1609 | GenBank | No | 0.00 | 688 | dataset5 | 0 | 0 | 1 | 0 | Atib |
| JQ975178 | Atib | 2n | NA | AtARC1610 | GenBank | No | 0.00 | 688 | dataset5 | 0 | 0 | NA | 0 | Atib |
| MG572067 | Atib | 2n | NA | Yanabengcuo2002 | GenBank | No | 0.00 | 1413 | dataset6 | NA | NA | NA | 0 | Atib |
| MG572068 | Atib | 2n | NA | Xizang1997 | GenBank | No | 0.00 | 1413 | dataset6 | NA | NA | NA | 0 | Atib |
| MG572069 | Atib | 2n | NA | Anduoshuang1998 | GenBank | No | 0.00 | 1418 | dataset6 | NA | NA | NA | 0 | Atib |
| MG572070 | Atib | 2n | NA | Naqudongqiao1996 | GenBank | No | 0.00 | 1418 | dataset6 | NA | NA | NA | 0 | Atib |
| MG572071 | Atib | 2n | NA | Qixianghu2002 | GenBank | No | 0.00 | 1418 | dataset6 | NA | NA | NA | 0 | Atib |
| EF615584 | Atib | 2n | NA | ARC1609 | GenBank | No | 0.00 | 622 | dataset6 | NA | NA | NA | 0 | Atib |
| EF615585 | Atib | 2n | NA | ARC1610 | GenBank | No | 0.00 | 622 | dataset6 | NA | NA | NA | 0 | Atib |
| EF615586 | Atib | 2n | NA | ARC1611 | GenBank | No | 0.00 | 622 | dataset6 | NA | NA | NA | 0 | Atib |
| EF615587 | Atib | 2n | NA | ARC1348 | GenBank | No | 0.00 | 718 | dataset5 | 1.99 | 1 | 7 | 1 | Atib |
| EF615588 | Atib | 2n | NA | ARC1524 | GenBank | No | 0.00 | 739 | dataset5 | 1.99 | 1 | NA | 1 | Atib |
| KF707855 | Atib | 2n | NA | ATLAG1 | GenBank | No | 0.00 | 614 | dataset5 | 0 | 0 | NA | 0 | Atib |
| KF707856 | Atib | 2n | NA | ATLAG2 | GenBank | No | 0.00 | 614 | dataset5 | 0 | 0 | NA | 0 | Atib |
| KF707857 | Atib | 2n | NA | ATLAG3 | GenBank | No | 0.00 | 614 | dataset5 | 0 | 0 | NA | 0 | Atib |
| KF707858 | Atib | 2n | NA | ATLAG7 | GenBank | No | 0.00 | 614 | dataset5 | 0 | 0 | NA | 0 | Atib |
| KF707859 | Atib | 2n | NA | ATLAG10 | GenBank | No | 0.00 | 614 | dataset5 | 0 | 0 | NA | 0 | Atib |
| KF707860 | Atib | 2n | NA | ATLAG12 | GenBank | No | 0.00 | 614 | dataset5 | 0 | 0 | NA | 0 | Atib |
| KF707861 | Atib | 2n | NA | ATLAG8 | GenBank | No | 0.00 | 614 | dataset5 | 0 | 0 | NA | 0 | Atib |
| KF707862 | Atib | 2n | NA | ATLAG4 | GenBank | No | 0.00 | 614 | dataset5 | 0 | 0 | NA | 0 | Atib |
| KF707863 | Atib | 2n | NA | ATLAG5 | GenBank | No | 0.00 | 614 | dataset5 | 0 | 0 | NA | 0 | Atib |
| KF707864 | Atib | 2n | NA | ATLAG6 | GenBank | No | 0.00 | 614 | dataset5 | 0 | 0 | NA | 0 | Atib |
| KF707895 | Atib | 2n | NA | ATGAI4 | GenBank | No | 0.00 | 614 | dataset5 | 0 | 0 | NA | 0 | Atib |
| KF707896 | Atib | 2n | NA | ATGAI1 | GenBank | No | 0.00 | 614 | dataset5 | 0 | 0 | NA | 0 | Atib |
| KF707897 | Atib | 2n | NA | ATGAI5 | GenBank | No | 0.00 | 614 | dataset5 | 0 | 0 | NA | 0 | Atib |
| KF707898 | Atib | 2n | NA | ATGAI2 | GenBank | No | 0.00 | 614 | dataset5 | 0 | 0 | NA | 0 | Atib |
| KF707899 | Atib | 2n | NA | ATGAI3 | GenBank | No | 0.00 | 614 | dataset5 | 0 | 0 | NA | 0 | Atib |
| KF707900 | Atib | 2n | NA | ATHAY1 | GenBank | No | 0.00 | 614 | dataset5 | 0 | 0 | 1 | 0 | Atib |
| KF707901 | Atib | 2n | NA | ATHAY2 | GenBank | No | 0.00 | 614 | dataset5 | 0 | 0 | NA | 0 | Atib |
| KF707902 | Atib | 2n | NA | ATHAY3 | GenBank | No | 0.00 | 614 | dataset5 | 0 | 0 | NA | 0 | Atib |
| KF707903 | Atib | 2n | NA | ATHAY4 | GenBank | No | 0.00 | 614 | dataset5 | 0 | 0 | NA | 0 | Atib |
| KF707904 | Atib | 2n | NA | ATHAY5 | GenBank | No | 0.00 | 614 | dataset5 | 0 | 0 | 1 | 0 | Atib |
| KF707905 | Atib | 2n | NA | ATHAY6 | GenBank | No | 0.00 | 614 | dataset5 | 0 | 0 | NA | 0 | Atib |
| KF707906 | Atib | 2n | NA | ATHAY8 | GenBank | No | 0.00 | 614 | dataset5 | 0 | 0 | 1 | 0 | Atib |
| KF707907 | Atib | 2n | NA | ATHAY9 | GenBank | No | 0.00 | 614 | dataset5 | 0 | 0 | NA | 0 | Atib |
| KF707908 | Atib | 2n | NA | ATHAY10 | GenBank | No | 0.00 | 614 | dataset5 | 0 | 0 | NA | 0 | Atib |
| KF707909 | Atib | 2n | NA | ATJIN1 | GenBank | No | 0.00 | 614 | dataset5 | 0 | 0 | NA | 0 | Atib |
| KF707910 | Atib | 2n | NA | ATJIN2 | GenBank | No | 0.00 | 614 | dataset5 | 0 | 0 | 2 | 1 | Atib |
| KF707911 | Atib | 2n | NA | ATJIN3 | GenBank | No | 0.00 | 614 | dataset5 | 0 | 0 | NA | 0 | Atib |
| KF707912 | Atib | 2n | NA | ATJIN4 | GenBank | No | 0.00 | 614 | dataset5 | 0 | 0 | 1 | 0 | Atib |
| KF707913 | Atib | 2n | NA | ATJIN5 | GenBank | No | 0.00 | 614 | dataset5 | 0 | 0 | NA | 0 | Atib |
| KF707914 | Atib | 2n | NA | ATJIN6 | GenBank | No | 0.00 | 614 | dataset5 | 0 | 0 | NA | 0 | Atib |
| KF707915 | Atib | 2n | NA | ATJIN7 | GenBank | No | 0.00 | 614 | dataset5 | 0 | 0 | NA | 0 | Atib |
| KF707916 | Atib | 2n | NA | ATJIN8 | GenBank | No | 0.00 | 614 | dataset5 | 0 | 0 | NA | 0 | Atib |
| KF707917 | Atib | 2n | NA | ATJIN9 | GenBank | No | 0.00 | 614 | dataset5 | 0 | 0 | NA | 0 | Atib |
| KF707918 | Atib | 2n | NA | ATJIN10 | GenBank | No | 0.00 | 614 | dataset5 | 0 | 0 | NA | 0 | Atib |
| KF707919 | Atib | 2n | NA | ATLAG13 | GenBank | No | 0.00 | 614 | dataset5 | 0 | 0 | NA | 0 | Atib |
| KF707920 | Atib | 2n | NA | ATLAG14 | GenBank | No | 0.00 | 614 | dataset5 | 0 | 0 | NA | 0 | Atib |
| KF707921 | Atib | 2n | NA | ATLAG15 | GenBank | No | 0.00 | 614 | dataset5 | 0 | 0 | NA | 0 | Atib |
| KF707922 | Atib | 2n | NA | ATLAG16 | GenBank | No | 0.00 | 614 | dataset5 | 0 | 0 | NA | 0 | Atib |
| KF707923 | Atib | 2n | NA | ATLAG17 | GenBank | No | 0.00 | 614 | dataset5 | 0 | 0 | 1 | 0 | Atib |
| KF707924 | Atib | 2n | NA | ATLAG18 | GenBank | No | 0.00 | 614 | dataset5 | 0 | 0 | NA | 0 | Atib |
| KF707925 | Atib | 2n | NA | ATLAG19 | GenBank | No | 0.00 | 614 | dataset5 | 0 | 0 | NA | 0 | Atib |
| KF707926 | Atib | 2n | NA | ATLAG20 | GenBank | No | 0.00 | 614 | dataset5 | 0 | 0 | NA | 0 | Atib |
| KF707927 | Atib | 2n | NA | ATLAG21 | GenBank | No | 0.00 | 614 | dataset5 | 0 | 0 | 1 | 0 | Atib |
| KF707928 | Atib | 2n | NA | ATLAG22 | GenBank | No | 0.00 | 614 | dataset5 | 0 | 0 | 0 | 0 | Atib |
| KF691215 | Atib | 2n | NA | JIN1 | GenBank | No | 0.00 | 561 | dataset5 | 0 | 0 | 2 | 1 | Atib |
| KF691216 | Atib | 2n | NA | JIN2 | GenBank | No | 0.00 | 561 | dataset5 | 0 | 0 | 1 | 0 | Atib |
| KF691217 | Atib | 2n | NA | JIN3 | GenBank | No | 0.00 | 561 | dataset5 | 0 | 0 | 2 | 1 | Atib |
| KF691218 | Atib | 2n | NA | JIN4 | GenBank | No | 0.00 | 561 | dataset5 | 0 | 0 | 2 | 1 | Atib |
| KF691245 | Atib | 2n | NA | TIB11 | GenBank | No | 0.00 | 561 | dataset5 | 0 | 0 | 0 | 0 | Atib |
| KF691246 | Atib | 2n | NA | TIB12 | GenBank | No | 0.00 | 561 | dataset5 | 0 | 0 | 1 | 0 | Atib |
| KF691247 | Atib | 2n | NA | TIB13 | GenBank | No | 0.00 | 561 | dataset5 | 0 | 0 | NA | 0 | Atib |
| KF691248 | Atib | 2n | NA | TIB14 | GenBank | No | 0.00 | 561 | dataset5 | 0 | 0 | 1 | 0 | Atib |
| KF691249 | Atib | 2n | NA | TIB15 | GenBank | No | 0.00 | 561 | dataset5 | 0 | 0 | NA | 0 | Atib |
| KF691316 | Atib | 2n | NA | TIB21 | GenBank | No | 0.00 | 561 | dataset5 | 0 | 0 | 4 | 1 | Atib |
| KF691317 | Atib | 2n | NA | TIB22 | GenBank | No | 0.00 | 561 | dataset5 | 0 | 0 | 2 | 1 | Atib |
| KF691318 | Atib | 2n | NA | TIB23 | GenBank | No | 0.00 | 561 | dataset5 | 0 | 0 | 1 | 0 | Atib |
| KX925408 | Atib | 2n | NA | ATIB.1 | GenBank | No | 0.00 | 1412 | dataset6 | NA | NA | NA | 0 | Atib |
| KX925409 | Atib | 2n | NA | ATIB.2 | GenBank | No | 0.00 | 1405 | dataset6 | NA | NA | NA | 1 | Atib |
| MT791835 | Aurm | 2n | URM | AurmURM1 | This study | No | 0.00 | 685 | dataset1 | 0 | 0 | 1 | 0 | Aurm |
| MT791836 | Aurm | 2n | URM | AurmURM2 | This study | No | 0.00 | 686 | dataset1 | 0 | 0 | 1 | 0 | Aurm |
| MT791837 | Aurm | 2n | URM | AurmURM3 | This study | No | 0.00 | 650 | dataset1 | 0 | 0 | 1 | 0 | Aurm |
| MT791838 | Aurm | 2n | URM | AurmURM4 | This study | No | 0.00 | 686 | dataset1 | 0 | 0 | NA | 0 | Aurm |
| MT791839 | Aurm | 2n | URM | AurmURM5 | This study | No | 0.00 | 663 | dataset1 | 0 | 0 | 1 | 0 | Aurm |
| NA | Aurm | 2n | URM | AurmURM7 | This study | No | 0.00 | 685 | dataset2 | NA | NA | NA | 0 | Aurm |
| MT791840 | Aurm | 2n | URM | AurmURM8 | This study | No | 0.00 | 615 | dataset1 | 0 | 0 | NA | 0 | Aurm |
| NA | Aurm | 2n | URM | AurmURM10 | This study | No | 0.00 | 526 | dataset2 | NA | NA | NA | 0 | Aurm |
| MT791841 | Aurm | 2n | URM | AurmURM11 | This study | No | 0.00 | 617 | dataset1 | 0 | 0 | 0 | 0 | Aurm |
| NA | Aurm | 2n | URM | AurmURM12 | This study | No | 0.01 | 627 | dataset2 | NA | NA | NA | 1 | Aurm |
| MT791842 | Aurm | 2n | URM | AurmURM13 | This study | Yes | 0.00 | 581 | dataset3 | 0 | 0 | 1 | 0 | Aurm |
| MT792844 | Aurm | 2n | URM | numt1AurmURM13 | This study | Yes | 0.00 | 563 | dataset3 | 0 | 0 | 1 | 1 | Aurm |
| MT792845 | Aurm | 2n | URM | numt2AurmURM13 | This study | Yes | 0.00 | 571 | dataset3 | 0.06 | 0 | 0 | 1 | Aurm |
| DQ119651 | Aurm | 2n | NA | AurmHou | GenBank | No | 0.00 | 658 | dataset5 | 0 | 0 | NA | 0 | Aurm |
| JQ975176 | Aurm | 2n | NA | AuARC1227 | GenBank | No | 0.00 | 688 | dataset5 | 0 | 0 | 4 | 1 | Aurm |
| KF707681 | Aurm | 2n | NA | AUURM10 | GenBank | No | 0.00 | 614 | dataset5 | 0 | 0 | 1 | 1 | Aurm |
| KF707682 | Aurm | 2n | NA | AUURM9 | GenBank | No | 0.00 | 614 | dataset5 | 0 | 0 | NA | 0 | Aurm |
| KF707683 | Aurm | 2n | NA | AUURM8 | GenBank | No | 0.00 | 614 | dataset5 | 0 | 0 | 1 | 0 | Aurm |
| KF707684 | Aurm | 2n | NA | AUURM3 | GenBank | No | 0.00 | 614 | dataset5 | 0 | 0 | NA | 0 | Aurm |
| KF707685 | Aurm | 2n | NA | AUURM12 | GenBank | No | 0.00 | 614 | dataset5 | 0 | 0 | 3 | 1 | Aurm |
| KF707686 | Aurm | 2n | NA | AUURM11 | GenBank | No | 0.00 | 614 | dataset5 | 0.18 | 1 | 1 | 1 | Aurm |
| KF707687 | Aurm | 2n | NA | AUURM7 | GenBank | No | 0.00 | 614 | dataset5 | 0 | 0 | 3 | 1 | Aurm |
| KF707688 | Aurm | 2n | NA | AUURM5 | GenBank | No | 0.00 | 614 | dataset5 | 0 | 0 | 3 | 1 | Aurm |
| KF707689 | Aurm | 2n | NA | AUURM4 | GenBank | No | 0.00 | 614 | dataset5 | 0 | 0 | NA | 0 | Aurm |
| KF707690 | Aurm | 2n | NA | AUURM2 | GenBank | No | 0.00 | 614 | dataset5 | 0 | 0 | 1 | 0 | Aurm |
| KF707691 | Aurm | 2n | NA | AUKOY12 | GenBank | No | 0.00 | 614 | dataset5 | 0 | 0 | 1 | 0 | Aurm |
| KF707692 | Aurm | 2n | NA | AUKOY11 | GenBank | No | 0.00 | 614 | dataset5 | 0 | 0 | 2 | 1 | Aurm |
| KF707693 | Aurm | 2n | NA | AUKOY9 | GenBank | No | 0.00 | 614 | dataset5 | 0 | 0 | NA | 0 | Aurm |
| KF707694 | Aurm | 2n | NA | AUKOY8 | GenBank | No | 0.00 | 614 | dataset5 | 0 | 0 | NA | 1 | Aurm |
| KF707695 | Aurm | 2n | NA | AUKOY5 | GenBank | No | 0.00 | 614 | dataset5 | 0 | 0 | NA | 0 | Aurm |
| KF707696 | Aurm | 2n | NA | AUKOY4 | GenBank | No | 0.00 | 614 | dataset5 | 0 | 0 | NA | 1 | Aurm |
| KF707697 | Aurm | 2n | NA | AUKOY3 | GenBank | No | 0.00 | 614 | dataset5 | 0 | 0 | NA | 0 | Aurm |
| KF707698 | Aurm | 2n | NA | AUKOY2 | GenBank | No | 0.00 | 614 | dataset5 | 0 | 0 | NA | 0 | Aurm |
| KF707699 | Aurm | 2n | NA | AUKOY1 | GenBank | No | 0.00 | 614 | dataset5 | 0 | 0 | NA | 1 | Aurm |
| KF707875 | Aurm | 2n | NA | AUURM16 | GenBank | No | 0.00 | 614 | dataset5 | 0 | 0 | NA | 0 | Aurm |
| KF707876 | Aurm | 2n | NA | AUURM17 | GenBank | No | 0.00 | 614 | dataset5 | 0 | 0 | 1 | 0 | Aurm |
| KF707877 | Aurm | 2n | NA | AUURM18 | GenBank | No | 0.00 | 614 | dataset5 | 0 | 0 | NA | 0 | Aurm |
| KF707878 | Aurm | 2n | NA | AUURM19 | GenBank | No | 0.00 | 614 | dataset5 | 0 | 0 | 2 | 1 | Aurm |
| KF707879 | Aurm | 2n | NA | AUURM20 | GenBank | No | 0.00 | 614 | dataset5 | 0 | 0 | NA | 0 | Aurm |
| KF707880 | Aurm | 2n | NA | AUURM21 | GenBank | No | 0.00 | 614 | dataset5 | 0 | 0 | NA | 0 | Aurm |
| KF707881 | Aurm | 2n | NA | AUURM22 | GenBank | No | 0.00 | 614 | dataset5 | 0 | 0 | NA | 0 | Aurm |
| KF707882 | Aurm | 2n | NA | AUURM23 | GenBank | No | 0.00 | 614 | dataset5 | 0 | 0 | NA | 1 | Aurm |
| KF707883 | Aurm | 2n | NA | AUURM24 | GenBank | No | 0.00 | 614 | dataset5 | 0 | 0 | NA | 0 | Aurm |
| KF707884 | Aurm | 2n | NA | AUURM25 | GenBank | No | 0.00 | 614 | dataset5 | 0 | 0 | 1 | 0 | Aurm |
| KF691533 | Aurm | 2n | NA | KBG4 | GenBank | No | 0.00 | 561 | dataset5 | 0 | 0 | 1 | 0 | Aurm |
| JX512748 | Aurm | 2n | NA | NC21 | GenBank | No | 0.00 | 561 | dataset5 | 0 | 0 | NA | 0 | Aurm |
| JX512749 | Aurm | 2n | NA | NC22 | GenBank | No | 0.00 | 561 | dataset6 | NA | NA | NA | 0 | Aurm |
| JX512750 | Aurm | 2n | NA | NC23 | GenBank | No | 0.00 | 561 | dataset5 | 0 | 0 | 1 | 0 | Aurm |
| JX512751 | Aurm | 2n | NA | NC11 | GenBank | No | 0.00 | 561 | dataset6 | NA | NA | NA | 1 | Aurm |
| JX512752 | Aurm | 2n | NA | NC12 | GenBank | No | 0.00 | 561 | dataset5 | 0 | 0 | 2 | 1 | Aurm |
| JX512753 | Aurm | 2n | NA | NC13 | GenBank | No | 0.00 | 561 | dataset6 | NA | NA | NA | 0 | Aurm |
| JX512754 | Aurm | 2n | NA | NE1 | GenBank | No | 0.00 | 561 | dataset5 | 0 | 0 | 1 | 0 | Aurm |
| JX512755 | Aurm | 2n | NA | NE2 | GenBank | No | 0.00 | 561 | dataset5 | 0 | 0 | NA | 0 | Aurm |
| JX512756 | Aurm | 2n | NA | NE3 | GenBank | No | 0.00 | 561 | dataset5 | 0 | 0 | NA | 0 | Aurm |
| JX512757 | Aurm | 2n | NA | NE4 | GenBank | No | 0.00 | 561 | dataset5 | 0 | 0 | 1 | 0 | Aurm |
| JX512758 | Aurm | 2n | NA | NE5 | GenBank | No | 0.00 | 561 | dataset5 | 0 | 0 | NA | 0 | Aurm |
| JX512759 | Aurm | 2n | NA | NE6 | GenBank | No | 0.00 | 561 | dataset5 | 0.06 | 0 | 1 | 0 | Aurm |
| JX512760 | Aurm | 2n | NA | NW1 | GenBank | No | 0.00 | 561 | dataset5 | 0 | 0 | 3 | 1 | Aurm |
| JX512761 | Aurm | 2n | NA | NW2 | GenBank | No | 0.00 | 561 | dataset5 | 0 | 0 | 1 | 0 | Aurm |
| JX512762 | Aurm | 2n | NA | NW3 | GenBank | No | 0.00 | 561 | dataset5 | 0 | 0 | NA | 0 | Aurm |
| JX512763 | Aurm | 2n | NA | NW4 | GenBank | No | 0.00 | 561 | dataset5 | 0 | 0 | 2 | 1 | Aurm |
| JX512764 | Aurm | 2n | NA | NW5 | GenBank | No | 0.00 | 561 | dataset5 | 0 | 0 | NA | 0 | Aurm |
| JX512765 | Aurm | 2n | NA | NW6 | GenBank | No | 0.00 | 561 | dataset5 | 0 | 0 | 2 | 1 | Aurm |
| JX512766 | Aurm | 2n | NA | ME11 | GenBank | No | 0.00 | 561 | dataset5 | 0 | 0 | NA | 0 | Aurm |
| JX512767 | Aurm | 2n | NA | ME12 | GenBank | No | 0.00 | 561 | dataset5 | 0 | 0 | 2 | 1 | Aurm |
| JX512768 | Aurm | 2n | NA | ME21 | GenBank | No | 0.00 | 561 | dataset5 | 0 | 0 | 3 | 1 | Aurm |
| JX512769 | Aurm | 2n | NA | ME22 | GenBank | No | 0.00 | 561 | dataset5 | 0 | 0 | NA | 0 | Aurm |
| JX512770 | Aurm | 2n | NA | ME23 | GenBank | No | 0.00 | 561 | dataset5 | 0 | 0 | 1 | 0 | Aurm |
| JX512771 | Aurm | 2n | NA | ME24 | GenBank | No | 0.00 | 561 | dataset5 | 0 | 0 | NA | 0 | Aurm |
| JX512772 | Aurm | 2n | NA | ME25 | GenBank | No | 0.00 | 561 | dataset5 | 0 | 0 | 0 | 0 | Aurm |
| JX512773 | Aurm | 2n | NA | ME26 | GenBank | No | 0.00 | 561 | dataset5 | 0 | 0 | 1 | 0 | Aurm |
| JX512774 | Aurm | 2n | NA | MW11 | GenBank | No | 0.00 | 561 | dataset5 | 0 | 0 | 1 | 0 | Aurm |
| JX512775 | Aurm | 2n | NA | MW12 | GenBank | No | 0.00 | 561 | dataset5 | 0 | 0 | 1 | 0 | Aurm |
| JX512776 | Aurm | 2n | NA | MW13 | GenBank | No | 0.00 | 561 | dataset5 | 0 | 0 | NA | 0 | Aurm |
| JX512777 | Aurm | 2n | NA | MW14 | GenBank | No | 0.00 | 561 | dataset5 | 0 | 0 | 1 | 0 | Aurm |
| JX512778 | Aurm | 2n | NA | MW15 | GenBank | No | 0.00 | 561 | dataset5 | 0 | 0 | NA | 0 | Aurm |
| JX512779 | Aurm | 2n | NA | MW16 | GenBank | No | 0.00 | 561 | dataset5 | 0 | 0 | 1 | 0 | Aurm |
| JX512780 | Aurm | 2n | NA | MW21 | GenBank | No | 0.00 | 561 | dataset5 | 0 | 0 | NA | 0 | Aurm |
| JX512781 | Aurm | 2n | NA | MW22 | GenBank | No | 0.00 | 561 | dataset5 | 0 | 0 | 1 | 0 | Aurm |
| JX512782 | Aurm | 2n | NA | MW23 | GenBank | No | 0.00 | 561 | dataset5 | 0 | 0 | 4 | 1 | Aurm |
| JX512783 | Aurm | 2n | NA | MW24 | GenBank | No | 0.00 | 561 | dataset5 | 0 | 0 | NA | 0 | Aurm |
| JX512784 | Aurm | 2n | NA | MW25 | GenBank | No | 0.00 | 561 | dataset5 | 0 | 0 | 3 | 1 | Aurm |
| JX512785 | Aurm | 2n | NA | SE21 | GenBank | No | 0.00 | 561 | dataset5 | 0 | 0 | 1 | 0 | Aurm |
| JX512786 | Aurm | 2n | NA | SE22 | GenBank | No | 0.00 | 561 | dataset5 | 0 | 0 | 2 | 1 | Aurm |
| JX512787 | Aurm | 2n | NA | SE23 | GenBank | No | 0.00 | 561 | dataset5 | 0 | 0 | 2 | 1 | Aurm |
| JX512788 | Aurm | 2n | NA | SE24 | GenBank | No | 0.00 | 561 | dataset5 | 0 | 0 | NA | 0 | Aurm |
| JX512789 | Aurm | 2n | NA | SE25 | GenBank | No | 0.00 | 561 | dataset5 | 0 | 0 | 2 | 1 | Aurm |
| JX512790 | Aurm | 2n | NA | SE26 | GenBank | No | 0.00 | 561 | dataset5 | 0 | 0 | NA | 0 | Aurm |
| JX512791 | Aurm | 2n | NA | SE31 | GenBank | No | 0.00 | 561 | dataset5 | 0 | 0 | NA | 0 | Aurm |
| JX512792 | Aurm | 2n | NA | SE32 | GenBank | No | 0.00 | 561 | dataset6 | NA | NA | NA | 0 | Aurm |
| JX512793 | Aurm | 2n | NA | SE33 | GenBank | No | 0.01 | 561 | dataset6 | NA | NA | NA | 0 | Aurm |
| JX512794 | Aurm | 2n | NA | SC11 | GenBank | No | 0.00 | 561 | dataset5 | 0 | 0 | 0 | 0 | Aurm |
| JX512795 | Aurm | 2n | NA | SC12 | GenBank | No | 0.00 | 561 | dataset5 | 0 | 0 | NA | 0 | Aurm |
| JX512796 | Aurm | 2n | NA | SC13 | GenBank | No | 0.00 | 561 | dataset5 | 0 | 0 | NA | 0 | Aurm |
| JX512797 | Aurm | 2n | NA | SC21 | GenBank | No | 0.00 | 561 | dataset5 | 0 | 0 | 1 | 0 | Aurm |
| JX512798 | Aurm | 2n | NA | SC22 | GenBank | No | 0.00 | 561 | dataset5 | 0.18 | 1 | 1 | 1 | Aurm |
| JX512799 | Aurm | 2n | NA | SC23 | GenBank | No | 0.00 | 561 | dataset6 | NA | NA | NA | 0 | Aurm |
| JX512800 | Aurm | 2n | NA | SC31 | GenBank | No | 0.00 | 561 | dataset5 | 0 | 0 | 1 | 0 | Aurm |
| JX512801 | Aurm | 2n | NA | SC32 | GenBank | No | 0.00 | 561 | dataset5 | 0 | 0 | NA | 0 | Aurm |
| JX512802 | Aurm | 2n | NA | SC33 | GenBank | No | 0.00 | 561 | dataset5 | 0 | 0 | 1 | 0 | Aurm |
| JX512803 | Aurm | 2n | NA | SC34 | GenBank | No | 0.00 | 561 | dataset5 | 0 | 0 | NA | 0 | Aurm |
| JX512804 | Aurm | 2n | NA | SE11 | GenBank | No | 0.00 | 561 | dataset5 | 0 | 0 | NA | 0 | Aurm |
| JX512805 | Aurm | 2n | NA | SE12 | GenBank | No | 0.00 | 561 | dataset5 | 0 | 0 | NA | 0 | Aurm |
| JX512806 | Aurm | 2n | NA | SE13 | GenBank | No | 0.00 | 561 | dataset5 | 0 | 0 | 1 | 0 | Aurm |
| JX512807 | Aurm | 2n | NA | SW1 | GenBank | No | 0.00 | 561 | dataset5 | 0 | 0 | 2 | 1 | Aurm |
| JX512808 | Aurm | 2n | NA | SW2 | GenBank | No | 0.00 | 561 | dataset5 | 0 | 0 | NA | 0 | Aurm |
| MK682320 | Aurm | 2n | NA | URM1994.1 | GenBank | No | 0.00 | 631 | dataset5 | 0 | 0 | 1 | 0 | Aurm |
| MK682321 | Aurm | 2n | NA | URM1994.2 | GenBank | No | 0.00 | 631 | dataset5 | 0 | 0 | 1 | 0 | Aurm |
| MK682322 | Aurm | 2n | NA | URM1994.3 | GenBank | No | 0.00 | 631 | dataset5 | 0 | 0 | 1 | 0 | Aurm |
| MK682323 | Aurm | 2n | NA | URM1994.4 | GenBank | No | 0.00 | 631 | dataset5 | 0 | 0 | 1 | 0 | Aurm |
| MK682324 | Aurm | 2n | NA | URM1994.5 | GenBank | No | 0.00 | 631 | dataset5 | 0 | 0 | 1 | 0 | Aurm |
| MK682325 | Aurm | 2n | NA | URM1994.6 | GenBank | No | 0.00 | 631 | dataset5 | 0 | 0 | 4 | 1 | Aurm |
| MK682326 | Aurm | 2n | NA | URM1994.7 | GenBank | No | 0.00 | 631 | dataset5 | 0 | 0 | 3 | 1 | Aurm |
| MK682327 | Aurm | 2n | NA | URM1994.8 | GenBank | No | 0.00 | 631 | dataset5 | 0 | 0 | 0 | 0 | Aurm |
| MK682328 | Aurm | 2n | NA | URM1994.9 | GenBank | No | 0.00 | 631 | dataset5 | 0 | 0 | 3 | 1 | Aurm |
| MK682329 | Aurm | 2n | NA | URM1994.10 | GenBank | No | 0.00 | 631 | dataset5 | 0 | 0 | NA | 0 | Aurm |
| MK682330 | Aurm | 2n | NA | URM1994.11 | GenBank | No | 0.00 | 631 | dataset5 | 0 | 0 | 5 | 1 | Aurm |
| MK682331 | Aurm | 2n | NA | URM1994.12 | GenBank | No | 0.00 | 631 | dataset5 | 0.04 | 0 | 0 | 0 | Aurm |
| MK682332 | Aurm | 2n | NA | URM1994.13 | GenBank | No | 0.00 | 631 | dataset5 | 1.36 | 1 | 1 | 1 | Aurm |
| MK682333 | Aurm | 2n | NA | URM1994.14 | GenBank | No | 0.00 | 631 | dataset5 | 0 | 0 | NA | 0 | Aurm |
| MK682334 | Aurm | 2n | NA | URM1994.15 | GenBank | No | 0.00 | 631 | dataset5 | 0 | 0 | 2 | 1 | Aurm |
| MK682335 | Aurm | 2n | NA | URM1994.16 | GenBank | No | 0.00 | 631 | dataset5 | 0 | 0 | 1 | 0 | Aurm |
| MK682336 | Aurm | 2n | NA | URM1994.17 | GenBank | No | 0.00 | 631 | dataset5 | 0 | 0 | NA | 0 | Aurm |
| MK682337 | Aurm | 2n | NA | URM1994.18 | GenBank | No | 0.00 | 631 | dataset5 | 0 | 0 | 3 | 1 | Aurm |
| MK682338 | Aurm | 2n | NA | URM1994.19 | GenBank | No | 0.00 | 631 | dataset5 | 0 | 0 | NA | 0 | Aurm |
| MK682339 | Aurm | 2n | NA | URM1994.20 | GenBank | No | 0.00 | 631 | dataset5 | 0 | 0 | NA | 0 | Aurm |
| MK682340 | Aurm | 2n | NA | URM1994.21 | GenBank | No | 0.00 | 631 | dataset5 | 0 | 0 | 6 | 1 | Aurm |
| MK682341 | Aurm | 2n | NA | URM1994.22 | GenBank | No | 0.00 | 631 | dataset5 | 0 | 0 | 1 | 0 | Aurm |
| MK682342 | Aurm | 2n | NA | URM1994.23 | GenBank | No | 0.00 | 631 | dataset5 | 0 | 0 | 1 | 0 | Aurm |
| MK682343 | Aurm | 2n | NA | URM1994.24 | GenBank | No | 0.00 | 631 | dataset5 | 0 | 0 | 2 | 1 | Aurm |
| MK682344 | Aurm | 2n | NA | URM1994.25 | GenBank | No | 0.00 | 631 | dataset5 | 0 | 0 | NA | 1 | Aurm |
| MK682345 | Aurm | 2n | NA | URM1994.26 | GenBank | No | 0.00 | 631 | dataset5 | 0 | 0 | 1 | 0 | Aurm |
| MK682346 | Aurm | 2n | NA | URM1994.27 | GenBank | No | 0.00 | 631 | dataset5 | 0 | 0 | NA | 0 | Aurm |
| MK682347 | Aurm | 2n | NA | URM1994.28 | GenBank | No | 0.00 | 631 | dataset5 | 0 | 0 | NA | 0 | Aurm |
| MK682348 | Aurm | 2n | NA | URM1994.29 | GenBank | No | 0.00 | 631 | dataset5 | 0 | 0 | NA | 0 | Aurm |
| MK682349 | Aurm | 2n | NA | URM1994.30 | GenBank | No | 0.00 | 631 | dataset5 | 0 | 0 | 2 | 1 | Aurm |
| MK682350 | Aurm | 2n | NA | URM2004.1 | GenBank | No | 0.00 | 631 | dataset5 | 0 | 0 | 1 | 0 | Aurm |
| MK682351 | Aurm | 2n | NA | URM2004.2 | GenBank | No | 0.00 | 631 | dataset5 | 0 | 0 | NA | 0 | Aurm |
| MK682352 | Aurm | 2n | NA | URM2004.3 | GenBank | No | 0.00 | 631 | dataset5 | 0 | 0 | 2 | 1 | Aurm |
| MK682353 | Aurm | 2n | NA | URM2004.4 | GenBank | No | 0.00 | 631 | dataset5 | 0 | 0 | NA | 0 | Aurm |
| MK682354 | Aurm | 2n | NA | URM2004.5 | GenBank | No | 0.00 | 631 | dataset5 | 0 | 0 | 1 | 0 | Aurm |
| MK682355 | Aurm | 2n | NA | URM2004.6 | GenBank | No | 0.00 | 631 | dataset5 | 0 | 0 | 4 | 1 | Aurm |
| MK682356 | Aurm | 2n | NA | URM2004.7 | GenBank | No | 0.00 | 631 | dataset5 | 0 | 0 | NA | 0 | Aurm |
| MK682357 | Aurm | 2n | NA | URM2004.8 | GenBank | No | 0.00 | 631 | dataset5 | 0 | 0 | 2 | 1 | Aurm |
| MK682358 | Aurm | 2n | NA | URM2004.9 | GenBank | No | 0.00 | 631 | dataset5 | 0 | 0 | NA | 0 | Aurm |
| MK682359 | Aurm | 2n | NA | URM2004.10 | GenBank | No | 0.00 | 631 | dataset5 | 0 | 0 | NA | 0 | Aurm |
| MK682360 | Aurm | 2n | NA | URM2004.11 | GenBank | No | 0.00 | 631 | dataset5 | 0 | 0 | 2 | 1 | Aurm |
| MK682361 | Aurm | 2n | NA | URM2004.12 | GenBank | No | 0.00 | 631 | dataset5 | 0 | 0 | 1 | 0 | Aurm |
| MK682362 | Aurm | 2n | NA | URM2004.13 | GenBank | No | 0.00 | 631 | dataset5 | 0 | 0 | 1 | 0 | Aurm |
| MK682363 | Aurm | 2n | NA | URM2004.14 | GenBank | No | 0.00 | 631 | dataset5 | 0 | 0 | NA | 0 | Aurm |
| MK682364 | Aurm | 2n | NA | URM2004.15 | GenBank | No | 0.00 | 631 | dataset5 | 0 | 0 | 2 | 1 | Aurm |
| MK682365 | Aurm | 2n | NA | URM2004.16 | GenBank | No | 0.00 | 631 | dataset5 | 0 | 0 | NA | 0 | Aurm |
| MK682366 | Aurm | 2n | NA | URM2004.17 | GenBank | No | 0.00 | 631 | dataset5 | 0 | 0 | NA | 0 | Aurm |
| MK682367 | Aurm | 2n | NA | URM2004.18 | GenBank | No | 0.00 | 631 | dataset5 | 0 | 0 | 1 | 0 | Aurm |
| MK682368 | Aurm | 2n | NA | URM2004.19 | GenBank | No | 0.00 | 631 | dataset5 | 0 | 0 | NA | 0 | Aurm |
| MK682369 | Aurm | 2n | NA | URM2004.20 | GenBank | No | 0.00 | 631 | dataset5 | 0 | 0 | NA | 0 | Aurm |
| MK682370 | Aurm | 2n | NA | URM2004.21 | GenBank | No | 0.00 | 631 | dataset5 | 0 | 0 | 1 | 0 | Aurm |
| MK682371 | Aurm | 2n | NA | URM2004.22 | GenBank | No | 0.00 | 631 | dataset5 | 0 | 0 | 1 | 0 | Aurm |
| MK682372 | Aurm | 2n | NA | URM2004.23 | GenBank | No | 0.00 | 631 | dataset5 | 0.08 | 0 | 0 | 0 | Aurm |
| MK682373 | Aurm | 2n | NA | URM2004.24 | GenBank | No | 0.00 | 631 | dataset5 | 0 | 0 | 1 | 0 | Aurm |
| MK682374 | Aurm | 2n | NA | URM2004.25 | GenBank | No | 0.00 | 631 | dataset5 | 0 | 0 | NA | 0 | Aurm |
| MK682375 | Aurm | 2n | NA | URM2004.26 | GenBank | No | 0.00 | 631 | dataset5 | 0 | 0 | 2 | 1 | Aurm |
| MK682376 | Aurm | 2n | NA | URM2004.27 | GenBank | No | 0.00 | 631 | dataset5 | 0 | 0 | 2 | 1 | Aurm |
| MK682377 | Aurm | 2n | NA | URM2004.28 | GenBank | No | 0.00 | 631 | dataset5 | 0 | 0 | NA | 0 | Aurm |
| MK682378 | Aurm | 2n | NA | URM2004.29 | GenBank | No | 0.00 | 631 | dataset5 | 0 | 0 | NA | 1 | Aurm |
| MK682379 | Aurm | 2n | NA | URM2004.30 | GenBank | No | 0.00 | 631 | dataset5 | 0 | 0 | NA | 0 | Aurm |

**Table S5. List of the 1196 sequences collapsed into the 123 COI haplotypes whose network is represented in Fig. 5** (after removal of putative COI-numts and chimeras, except that from *Asin* from East Siberia). 399 sequences (hereafter “CEFE sequences”) were obtained for this study either directly (dataset1 and dataset2) or following a cloning step (dataset3). 797 additional sequences were obtained from the GenBank database (“GenBank” sequences) whose species and ploidy correspond to their original assignment. We retained respectively 237 CEFE and 748 GenBank putative COI sequences for further analyses (i.e. sequences with any ambiguous site and including less that 538 sites within the final alignment were removed). After excluding the 32 known numts, we found among the remaining 953 putative COI sequences 3 CEFE and 100 GenBank sequences that were potential numts or chimeras (assigned based on polarity changes in the amino acid sequence or on an unusually high number of mutations shared with another unrelated sequence of the dataset). Only sequences of high quality (dataset1 and dataset3) were deposited into GenBank.
